# The evolution of multi-component weapons in the superfamily of leaf-footed bugs

**DOI:** 10.1101/2023.04.24.538071

**Authors:** Christine W. Miller, Rebecca T. Kimball, Michael Forthman

## Abstract

Sexually selected weapons, such as the antlers of deer, claws of crabs, and tusks of beaked whales, are strikingly diverse across taxa and even within groups of closely related species. Phylogenetic comparative studies have typically taken a simplified approach to investigating the evolution of weapon diversity, examining the gains and losses of entire weapons, major shifts in size or type, or changes in location. Less understood is how individual weapon components evolve and assemble into a complete weapon. We addressed this question by examining weapon evolution in the diverse, multi-component hind-leg and body weapons of leaf-footed bugs, Superfamily Coreoidea (Hemiptera: Heteroptera). Male leaf-footed bugs use their weapons to fight for access to mating territories. We used a large multilocus dataset comprised of ultraconserved element loci for 248 species and inferred evolutionary transitions among component states using ancestral state estimation. We found that weapons added components over time with some evidence of a cyclical evolutionary pattern — gains of components followed by losses and then gains again. Further, we found that certain trait combinations evolved repeatedly across the phylogeny. This work reveals the remarkable and dynamic evolution of weapon form in the leaf-footed bugs. It also highlights that multi-component weapons may be especially useful in providing insights into the evolutionary interplay of form and function.

**TEASER TEXT:** For centuries, humans have been fascinated by the morphological weapons animals use to engage in battle. The diversity of sexually selected weapons is surprising, with considerable variation across even closely related groups of animals. Studies are needed that take a detailed view of the components that comprise weapons and the evolutionary assembly of these components into a complete structure. Here, we reconstruct the evolution of a multi-component weapon in a superfamily of insects. Male leaf-footed bugs use spiky, enlarged hind legs to wrestle over mating territories. We measured 15 putative weapon components across 248 species, using phylogenetic comparative analyses. We found that the number of weapon components generally increased over time, with many gains and losses of components along the way. We found that certain components were more likely to evolve with others, suggesting that specific trait combinations might be especially functional in battle. This work highlights that evolutionary studies of complex, multi-component weapons may be useful for reconstructing the evolutionary assembly of weapons and the interplay of form and function.

## INTRODUCTION

From the fearsome tusks of prehistoric elephants to the branching “antlers” of antler flies, sexually selected weapons are as diverse as they are captivating. Along with stunning variability across taxa, morphological weapons are highly diverse within groups of closely related species (Emlen 2008). For example, head weapons on extinct and extant members of the giraffe family include small ossicones, armored helmets, large flat expansions in the shape of butterfly wings, and paddle-like headgear that resembles the antlers of moose (Wang et al. 2022). The fascinating diversity of animal weapons can aid understanding of the evolutionary interplay of form and function because weapons have been selected to perform in physical combat (McCullough et al. 2016). For this reason, even small, intricate differences in morphology may reveal differences in fighting behavior across species and be tied to meaningful fitness consequences within species. A focus on weapon size alone can overlook the functional elements that contribute to fighting success (as described in Dennenmoser and Christy 2013; Palaoro and Peixoto 2022). Yet, the focus of most phylogenetic comparative studies of weapon morphology has been confined to major changes in weapon size or type, gains and losses of entire weapons, and shifts in location on the body (Emlen et al. 2005; Heinze et al. 2005; Hosoya and Araya 2005; Schutze et al. 2007; Dalebout et al. 2008; Cabrera and Stankowich 2020). Work that considers the individual components of weapons can inform unresolved evolutionary questions about weapon assembly. For example, such work can address whether extra components are added to weapons over time (Geist 1966) as might be expected in an evolutionary arms race (Emlen 2008) or run-away sexual selection (Moore et al. 2022). Further, it can examine the extent to which certain sets of weapon components appear together repeatedly across the phylogeny, suggesting a coordinated function during battle.

Combat is often a full-body sport. Males engaged in male-male competition frequently launch their entire bodies towards rivals. Sexually selected weapons often serve as the contact points in battle, while supporting traits such as the muscles contract, the feet or other structures anchor the body to the substrate, and the nervous system processes information and adjusts movements accordingly. For the purposes of this study, the morphological weapons of sexual selection are defined as the elaborated structures that frequently contact other individuals during intrasexual contests. These weapons may be quite simple, such as the head horns of dik-diks, the spurs of galliform birds, or the fangs of musk deer. Yet, some animals can become highly weaponized possessing weapon components stretching across vast areas of the body (e.g., Miyatake 1997; Miller and Emlen 2010; Dennenmoser and Christy 2013; O’Brien et al. 2017; Figure 1). We can consider these weapon systems when one or more joints (fulcrums) are co-opted, and when multiple components operate in a coordinated fashion, heightening weapon functionality.

**Figure 1.**
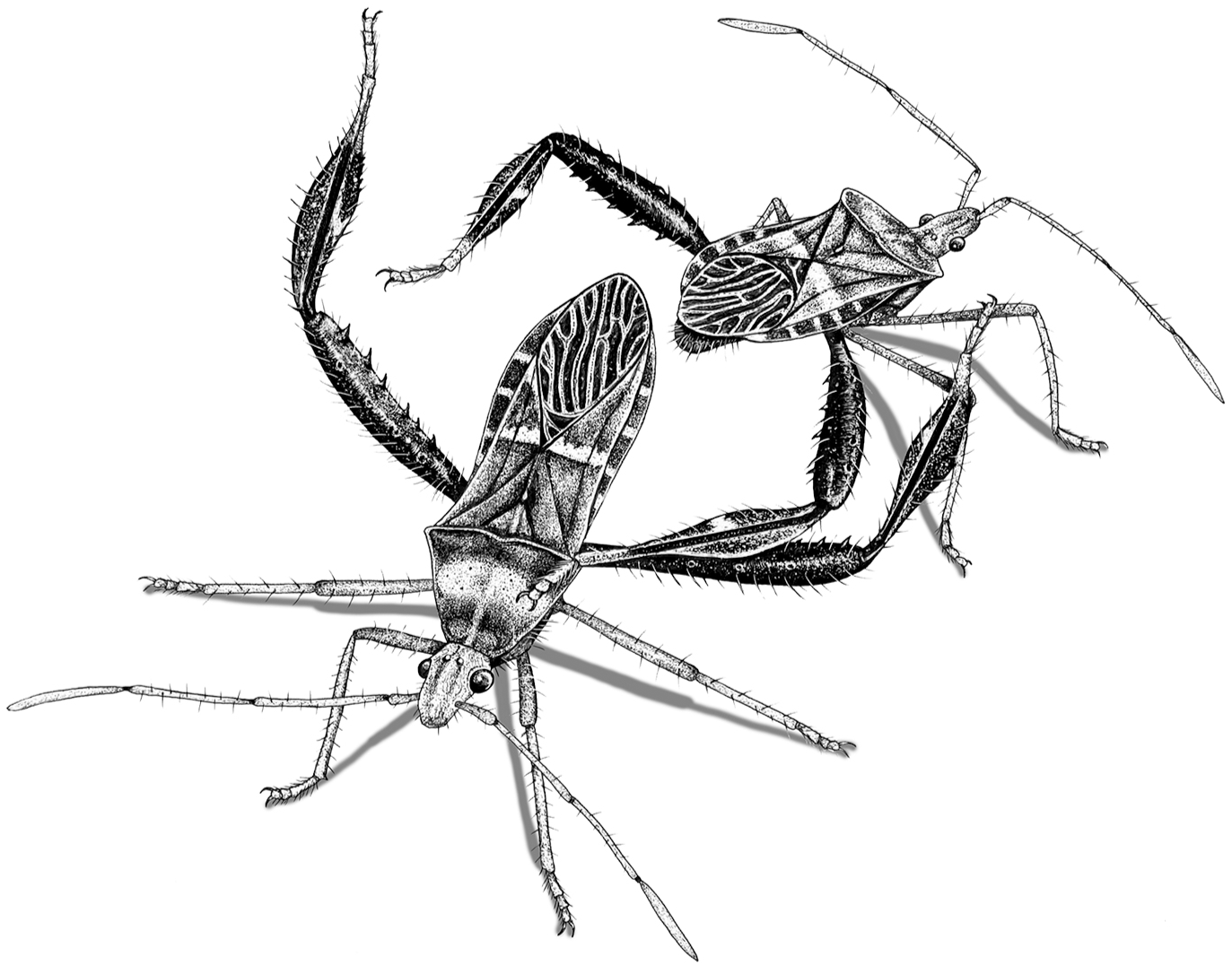
Leaf-footed cactus bugs, *Narnia femorata* Stål, 1892, engaged in an end-to-end battle. As seen here, many male leaf-footed bugs jockey into position, then press the spines on their femur into their opponent’s body. Males of this species show enlarged hind legs with spines and flags, including seven of the 15 weapon components studied (Table 1; Figure 4). Illustration by David J. Tuss.

**Table 1.**
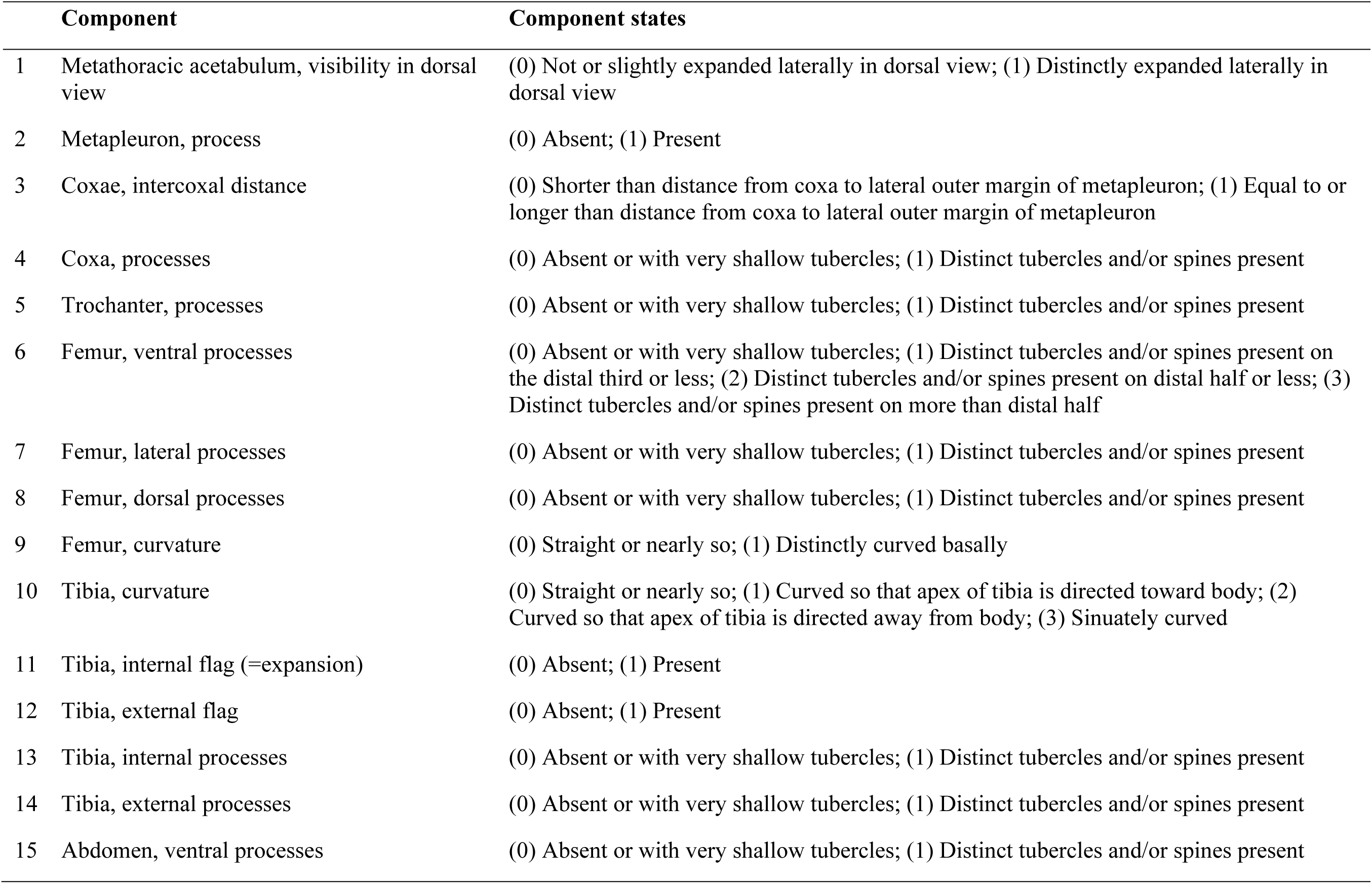
Morphological components and component state coding. Leg components correspond only to the hind legs (Components #3– 14).

**Table 2.**
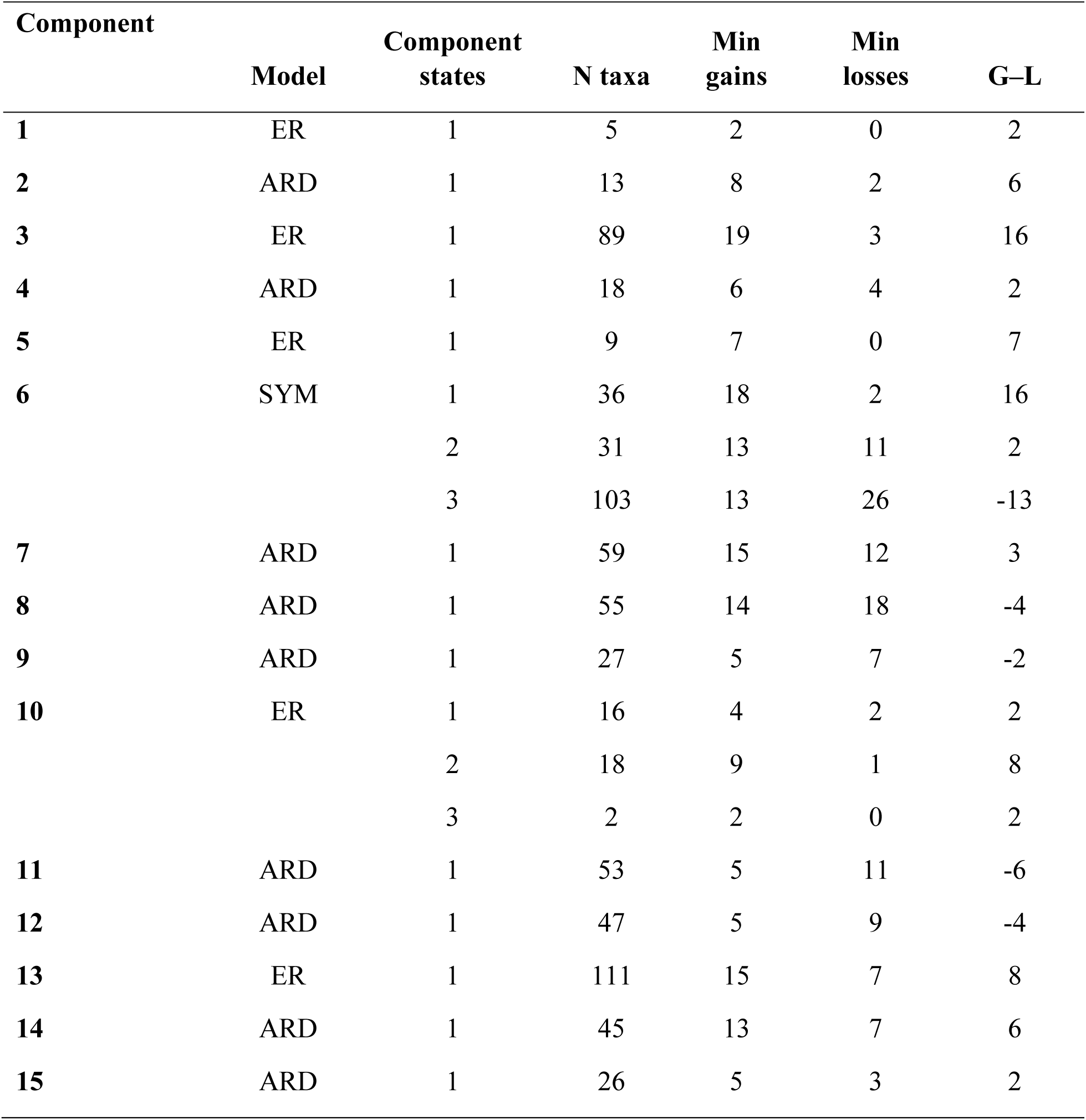
Summary of the minimum number of component state gains and losses across different ultrametric trees (component state 0 not reported). For results specific the 50p ML, 50p MSC, and 50p25mi MSC ultrametric trees, see Table S4. Abbreviations: ARD, all rates different model; ER, equal rates model; G–L, gains minus losses; Min, minimum; N, sample size; SYM, symmetric model.

Although weapon systems have been rarely studied in biomechanical detail outside the crustaceans (Sneddon et al. 2000; Levinton and Allen 2005; Dennenmoser and Christy 2013; Bywater et al. 2015), these systems may enable versatile combat maneuvers to exploit the prevailing context and may increase the capacity for rival manipulation, enabling a male to shift and hold a rival in position as he is pinched, punctured, or crushed. Since weapon components are often extremely tough, they may also serve as armor to protect the animal from bodily injury during combat. For all these reasons and more, weapon systems warrant further study. Additionally, examining the evolution of multi-component weapons provides outstanding opportunities to trace the assembly and disassembly of weapons over time.

We examined the evolution of 15 components of a weapon system found in a fascinating group of armed insects, the leaf-footed bugs and allies (Hemiptera: Coreoidea; Figures 1, 2). This group includes ∼3,300 species in five extant families, and it is one of only a few animal groups that produce weapons on the hind legs (Rico-Guevara and Hurme 2019). In some species, the hind legs are slim and non-elaborated (Figure 2A); males in these species typically do not engage in male-male combat. In other species, the hind legs exhibit extreme modifications including weapon components such as robust spines, club-like expansions, flags, and serrations (Figure 2B-G; CoreoideaSFTeam 2022). In fact, it is the striking, elaborated hind legs that give the common name “leaf-footed bugs” to the Coreidae, the largest family within the Coreoidea. For simplicity, we will hereafter refer to this superfamily as the “leaf-footed bugs”.

**Figure 2.**
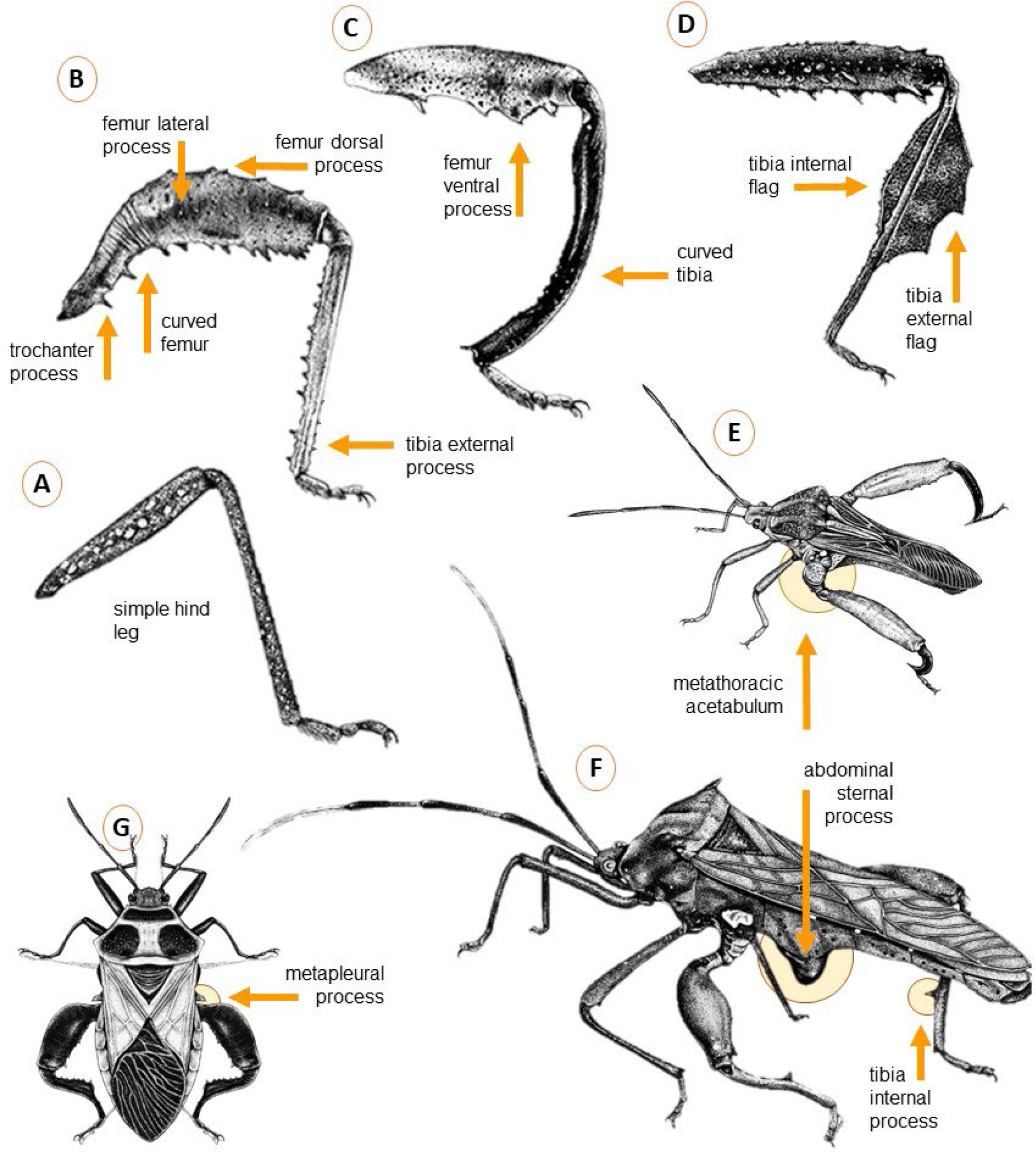
Illustrations of diverse hind leg shapes (A–D) and body shapes (E–G) in the leaf-footed bug superfamily. Arrows and text show many of the weapon components examined. Featured are the hind legs of: (A) *Anasa tristis* (De Geer, 1773), (B) *Camptischium clavipes* (Fabricius, 1803), (C) *Hyalymenus subinermis* Van Duzee, 1923, and (D) *Leptoglossus gonagra* (Fabricius, 1775). Full-body specimens include: (E) *Alcocerniella limonensis* Brailovsky, 1999 (F) *Mictis longicornis* Westwood, 1842(G) *Sagotylus confluens* (Say, 1832). Illustrations by David J. Tuss.

### The leaf-footed bugs: weapon morphology and behavior

The morphological elaborations on leaf-footed bug hind legs (Figure 2B-D) can, at times, extend to parts of the thorax and abdomen (Figure 2E-G). Where such elaboration exists, it is typically greater in males and is used in fighting (e.g., Figure 1). Male fighting maneuvers are varied; males may lunge, kick, squeeze, slap, pierce, and tear at their rivals (Figure 1; Mitchell 1980; Fujisaki 1981; Miyatake 1993, 1995, 1997; Eberhard 1998; Okada et al. 2011; Tatarnic and Spence 2013; Nolen et al. 2017; Emberts et al. 2021; Emberts and Wiens 2021). Some species engage in escalated combat in a ventral-ventral position when hanging from a plant surface (e.g., the crusader bug, *Mictis profana* (Fabricius, 1803); Tatarnic and Spence 2013), while others squeeze each other end-to-end (e.g., the heliconia bug, *Leptoscelis tricolor* Westwood, 1842; Miller and Emlen 2010). Successful males typically establish territories on plant resources that females need for feeding and laying eggs. Species that fight differently likely experience selection differently on their form, leading to the evolution of varied weapon components.

Indeed, many mictine species (such as *Mictis profana* and *Mictis longicornis* Westwood, 1842) possess a bizarre ventral horn—an abdominal sternal process—that is jabbed at the other male’s horn during ventral-ventral contests (Figure 2F; Tatarnic and Spence 2013). Fighting injuries are common in leaf-footed bugs and can include legs that are severed or missing, punctures to the legs and abdomen (G. Raina, *in prep*), and torn or punctured wings (G. Raina*, in prep*; Emberts et al. 2021; Emberts and Wiens 2021).

As in elk (Metz et al. 2018) and many other species (Rojas and Burdfield-Steel 2017; Lane 2018), the sexually selected weapons of leaf-footed bugs have functions beyond male-male competition. For example, they serve an important role in locomotion, and they are involved in predator defense. When attacked, some leaf-footed bugs squeeze attackers with their hind legs (Author, *personal observation*), in some cases drawing blood (U. Somjee, *personal communication*). Interestingly, the hind legs of leaf-footed bugs do not appear to be used in mate choice; indeed, experimentally blinded female *Riptortus pedestris* (Fabricius, 1775) do not show differences in mating behavior (Numata et al. 1986), and in the absence of male-male dynamics, female *Narnia femorata* show no reluctance to mate with a male missing a hind limb (Authors et al., *unpublished data*). Instead, chemical cues (Numata et al. 1986; Aldrich 1988; Wang and Millar 2000) and tactile/auditory cues (e.g., vibration, Numata et al. 1986; tapping, Miller 2008; and stridulation, Shestakov 2009) may be more influential in mate choice. Females rarely fight; when they do, it is typically with less intensity than males, and the conflict appears to center on feeding or oviposition sites (Eberhard 1998; Author, *personal observation*). Our focus in this study is the evolution of male weapon morphology, with work forthcoming on the evolution of sexual dimorphism in this superfamily.

Here, we provide the first phylogenetic analysis to investigate the evolution of male weapons across the leaf-footed bugs and one of the first studies across taxa that addresses the separate evolution of multiple weapon components (see also, Chow et al. 2021). We capitalized upon the multi-component nature of the leaf-footed bug weapon by focusing on discrete components, rather than simplifying or generalizing body form. We used 243 ingroup taxa from the insect families Alydidae, Coreidae, and Rhopalidae, as well as five outgroup taxa in the Pentatomomorpha (Table S1). Our sampling included a diverse representation of male hind leg morphologies within leaf-footed bugs, as well as of the many subfamilies and tribes found across major biogeographic regions. We inferred a phylogeny of the superfamily using ultraconserved element (UCE) loci. We then investigated the evolutionary lability of each of these components with ancestral state estimation (ASE). We asked: 1) How evolutionary labile are weapons and their components? 2) Does the number of weapon components increase over time? 3) Do certain components co-occur repeatedly, suggesting a coordinated function during battle?

## METHODS

### Selection of morphological components for study

Numerous leaf-footed bugs, such as *Jadera haematoloma* (Herrich-Schäffer, 1847), *Savius diversicornis* (Westwood, 1842), and *Anasa tristis* (De Geer, 1773), have simple, streamlined bodies and legs (e.g., Figure 2A, Table 1). The females in such species are typically larger than males and exhibit a rounded abdomen, but sex differences in morphology are otherwise minor. Male-male competition has not been reported in these species. Males of other leaf-footed bug species show modifications to this simple body plan including sharp spines, curves, and flags (Figure 2B-G), and these characters are often associated with fighting (Figure 1). Our goal with this study was to understand the evolution of such character elaborations. We selected 15 characters that are commonly modified, vary widely in their expression, and are straightforward to score objectively and reliably. Hereafter, we refer to the characters as weapon components.

The components, except for the metathoracic acetabulum (Figure 2E; Component #1), are typically sexually dimorphic when they are elaborated. Indeed, most of these components directly contact, and even injure, rivals during competition. Yet, much is unknown. Behavior has been documented in only a fraction of the thousands of leaf-footed bug species. Further, not all species with morphological elaboration engage in male-male competition (e.g., *Leptoglossus phyllopus* [Linnaeus, 1767], Mitchell 1980; *Anisoscelis alipes* Guérin-Méneville, 1833, Longbottom et al. 2022). We embrace the rich spectrum in morphology and behavior across leaf-footed bugs, acknowledging that most traits in most species have multiple uses and that the uses vary across the phylogeny. Indeed, our ultimate hope is to encourage work that examines the evolutionary interplay of morphology and behavior in this intriguing group of insects.

### Molecular data collection and phylogenetic inferences

For 216 taxa, we retrieved UCE sequence capture data from Forthman et al. (2019); Kieran et al. (2019); Emberts et al. (2020); Forthman et al. (2020); Forthman et al. (2022b); Miller et al. (2022) (Table S1). We also downloaded genome sequences of *Halyomorpha halys* (Stål, 1855) (Pentatomidae) and *Oncopeltus fasciatus* (Dallas, 1852) (Lygaeidae) from NCBI to extract UCE sequences from scaffolds. We generated new sequence data for 30 taxa following DNA extraction, isolation, and library construction approaches described in Forthman et al. (2019, 2020, 2022b). In short, sequence capture was done using baits designed from two pentatomomorphan taxa (Faircloth 2017; see Forthman et al. 2019) and using the touchdown capture protocol from Forthman et al. (2022a). Enriched library pools were combined into a single pool in equimolar amounts prior to sequencing on a single Illumina HiSeq3000 lane (2×100) at the University of Florida’s Interdisciplinary Center for Biotechnology Research.

Sequence reads were demultiplexed, adapter-trimmed, deduplicated, error-corrected, and assembled into contigs following Forthman et al. (2022a). We used PHYLUCE v1.7.0 (Faircloth 2016) to identify UCE loci from assembled contigs following (Forthman et al. 2019; Forthman et al. 2020; Forthman et al. 2022a). We also used the PHYLUCE to align UCE baits to two genome sequences (*Halyomorpha halys* and *Oncopeltus fasciatus*) and extract UCE loci with 500 bp of flanking nucleotides. A summary regarding newly generated read, contig, and UCE data are given in Table S2.

Loci were aligned individually with PHYLUCE using the --mafft setting (Katoh et al. 2002; Katoh and Standley 2013), and locus alignments were trimmed using trimAl v1.2 (Capella-Gutiérrez et al. 2009). Locus alignments with at least 50% and 70% of the total taxa were selected for analysis (referred to as “50p” and “70p” datasets, respectively). We also subsampled each of these datasets for the 25% most parsimony-informative loci (referred to as “25mi”), resulting in four datasets: 50p, 50p25mi, 70p, 70p25mi (see Table S3 for a summary of informative sites and number of UCE loci in each dataset).

For the 50p and 70p datasets, we concatenated locus alignments for maximum likelihood (ML) phylogenetic analysis, using the best model of sequence evolution and partitioning scheme identified by IQ-Tree v2.1.2 (Minh et al. 2020). For each dataset, ten separate partitioned ML analyses (Chernomor et al. 2016) were performed, with support measured by 1000 ultrafast bootstrap replicates (Hoang et al. 2018) and 1000 Shimodaira-Hasegawa-like approximate likelihood ratio test (sh-alrt) replicates (Guindon et al. 2010). We also inferred species trees for all four datasets under the multispecies coalescent (MSC) model to account for gene tree discordance due to incomplete lineage sorting (Degnan and Rosenberg 2006; Kubatko and Degnan 2007; Degnan and Rosenberg 2009; Roch and Steel 2015). We included the 50p25mi and 70p25mi datasets for species tree inference given that filtering for more informative loci has been shown to improve topological and branch lengths (in coalescent units) estimates in MSC analyses (Mirarab et al. 2014a; Hosner et al. 2016; Meiklejohn et al. 2016; Sayyari and Mirarab 2016; Sayyari et al. 2017; Forthman et al. 2022b). We estimated gene trees using the best-fit model of sequence evolution for each locus alignment using IQ-Tree, with near-zero branch lengths collapsed. Species trees were inferred from optimal gene trees using ASTRAL-III v5.7.7 (Mirarab et al. 2014b; Sayyari and Mirarab 2016; Zhang et al. 2018). We assessed clade support using local posterior probabilities (Sayyari and Mirarab 2016).

Prior to ASE, we transformed our 50p ML and 50p and 50p25mi MSC trees into ultrametric trees. First, we used IQ-Tree to estimate branch lengths as units of substitutions on the 50p and 50p25mi MSC topologies. We pruned outgroup taxa and used the *chronos* function in the *ape* package v5.6.1(Paradis and Schliep 2019) with R v4.1.2 (R Core Team 2021) to generate ultrametric trees under four models (correlated, discrete, relaxed, clock) and four values of lambda (0, 0.1, 1, 10). For more specific details on our molecular data collection and phylogenetic inferences, see Supplementary Materials.

### Ancestral state estimation

For ASE, 13 components were coded as binary, while two (Components #6 and #10) were treated as multistate components. For Component #6, we assigned three categories for the distribution of the ventral femoral processes, when present. Coding ventral femoral processes as a multistate component rather than a binary present/absent component allowed us to explore whether species with more elaborated hind legs often have a more extensive distribution of spines and tubercles on the ventral surface of the femur compared to species with less elaborated legs. Similarly, for Component #10, the tibia can exhibit four distinctive categories of curvature, and we treated these as separate states to explore patterns of gains and losses relative to other components. We primarily coded trait data from available specimen material, but in some cases, data was retrieved from type images and/or taxonomic descriptions of sampled species.

We used the *rayDISC* function in the R package *corHMM* v2.7 (Beaulieu et al. 2013) to estimate ancestral states for each component on the 50p ML and 50p and 50p25mi MSC ultrametric trees (70p and 70p25mi datasets were not included as these resulted in the same topologies and similar branch lengths [see Results and Figures S1–S6]; see Supplementary Methods for further details on generating ultrametric trees). We performed the analyses using marginal reconstruction and tested among the equal rates (ER), symmetric rates (SYM) (for multistate components only), and all rates different (ARD) models. To determine the best model for each component, we compared the Akaike Information Criterion corrected for small sample size (AICc) (Hurvich and Tsai 1989). We chose a difference of at least two among AICc values as a cut-off for determining a significant increase in model fit. When two models did not differ significantly in their AICc values, we selected the simplest model.

### Co-occurring and correlated components

To evaluate whether two putative weapon components co-occurred more or less than expected, we co-opted a probabilistic model of species co-occurrence using the R package *cooccur* v1.3 (Veech 2013; Griffith et al. 2016). For this analysis, we treated components as “species” and species as “sites” to determine whether two components were significantly associated with one another and whether such associations were positive or negative. This analysis requires data to be treated as binary presence-absence components. As such, we converted our two multistate components into binary ones (Component #6: ventral femoral processes [0] absent vs. [1] present; Component #10: tibia [0] straight vs. [1] curved) prior to analyses. For components that were not coded as absent or present (i.e., Components #1, #3, #9, and #10), we treated the plesiomorphic state based on ASE results as “absent”. Lastly, we excluded sites (i.e., species) that had missing data for at least one component.

Because *cooccur* does not account for phylogenetic non-independence among components, we also used the *fitPagel* function in the R package *phytools* v1.0 (Revell 2012) to determine whether two co-occurring components were correlated (i.e., we used *coocur* results as an *a priori* criterion for testing component pairs with *fitPagel*). This function tests for significant correlations between two binary components using Pagel’s (1994) method.

We note that correlated trait tests can produce misleading, significant associations when at least one component has a single evolutionary origin (see the “Darwin’s Scenario” and Figure 1 in Maddison and FitzJohn 2015). To address this situation *a priori*, we excluded significant *cooccur* component pairs from *fitPagel* testing if at least one component had only one evolutionary gain based on ASE results.

Correlated traits tests should produce significant results when there are replicated patterns of origins for both components throughout the phylogeny (see the “replicated co-distribution” and “replicated bursts” and Figure 1 in Maddison and FitzJohn 2015). However, while we can expect strong, statistical associations between component pairs when replicated patterns of origins for these components occur in nearby clades, this could be a *potentially* misleading result; the rate of origin for one component could potentially be explained by a third, unrelated component originating in a slightly larger clade that includes closely related “subclades” having independent gains of the second component (i.e., potential for unreplicated effects within lineages; Figure S7A; Maddison and FitzJohn 2015). The greater the number of evolutionary gains of component and the more dispersed the replicated evolutionary patterns are in the phylogeny, there is less concern of spurious associations (Maddison and FitzJohn 2015). Thus, we also assessed if both components often occurred within the same clades that were relatively dispersed throughout the phylogeny by comparing the ASE results of both components. We excluded significant *cooccur* component pairs from *fitPagel* testing if one component originated within clades that were in relatively close proximity on the phylogenetic tree (e.g., Figure S7A).

For the remaining component pairs, we used the same component matrix as for the *cooccur* analysis, excluded taxa with missing data for at least one component, and pruned ultrametric trees accordingly. For each test of a pair of components, we used marginal reconstruction with either the ER or ARD model of evolution (SYM = ER model with binary components) based on the best model found in our previous ASE analyses with *rayDISC*; if there were two different models for a pair of components (e.g., ER model for Component A and ARD model for Component B), we selected the most complex model for the *fitPagel* test given the use of a simpler model could potentially bias results due to model misspecification (Swofford et al. 2001; Lemmon and Moriarty 2004). We then ran three analyses for each pair of components, with two analyses having a different component treated as the dependent variable and one analysis having both components treated as an interdependent variable. A likelihood-ratio test was used to determine whether the null hypothesis (i.e., two components are independent of one another) can be rejected. We adjusted *p*-values with the Benjamini-Hochberg correction for multiple comparisons (Yang et al. 1994; Benjamini and Hochberg 1995), with the statistical significance threshold set to *p* < 0.05.

While our focus was to determine if co-occurrences among components were statistically supported by correlated traits test rather than on the directionality of component dependencies, we evaluated *a posteriori* whether the patterns of component evolution based on ASE results matched predictions of significant dependency models. For example, when a *fitPagel* test supported a single dependency model, we assessed whether the origin of one component generally preceded or evolved simultaneously with the other as would be predicted by the significant model. When patterns of evolution did not support the significant model(s) but appeared to support a non-significant model, we considered the correlated traits test to have produced a false positive result and did not report the two components as correlated in the absence of a significant alternative model (e.g., see Figure S7B).

We report all *fitPagel* results in the supplementary, and for significant *fitPagel* results, side-by-side comparisons of the corresponding ASE results are provided in Data File 1. However, in the Results section, we only report those statistically significant results we considered to be acceptable based on the criteria discussed above.

## RESULTS

### Familial to triable level phylogenetic relationships

Phylogenetic relationships among the families and subfamilies of leaf-footed bugs (Figures 3, S1–S6) were congruent with recent phylogenomic studies, which have supported the non-monophyly of Alydidae, Coreidae, Coreinae, and Meropachyinae (Forthman et al., 2019, 2020, 2022b; Emberts et al., 2020; Miller et al., 2022). Relationships among the tribes of Coreinae + Meropachyinae were also largely congruent with results of Forthman et al. (2020), however we found several differences. For example, we did not recover a monophyletic Acanthocorini or Dasynini. While we found high support for Clade E (Figure 3) as the sister group of Clade D in our MSC analyses (Figures S3–S6; congruent with Forthman et al. [2020]), our ML analyses found Clade E to be the sister group of Clade F + Clade G (Figures 3, S1, S2), also with high support. The phylogenetic position of Clade F was also unstable across our analyses (Figures 3, S1, S2, S6). Lastly, we also continued to find support for the polyphyly of Anisoscelini and Hypselonotini following Forthman et al. (2020), but we recovered an additional lineage of Anisoscelini (all analyses) and one (ML analyses) to two lineages (MSC analyses) of Hypselonotini.

**Figure 3.**
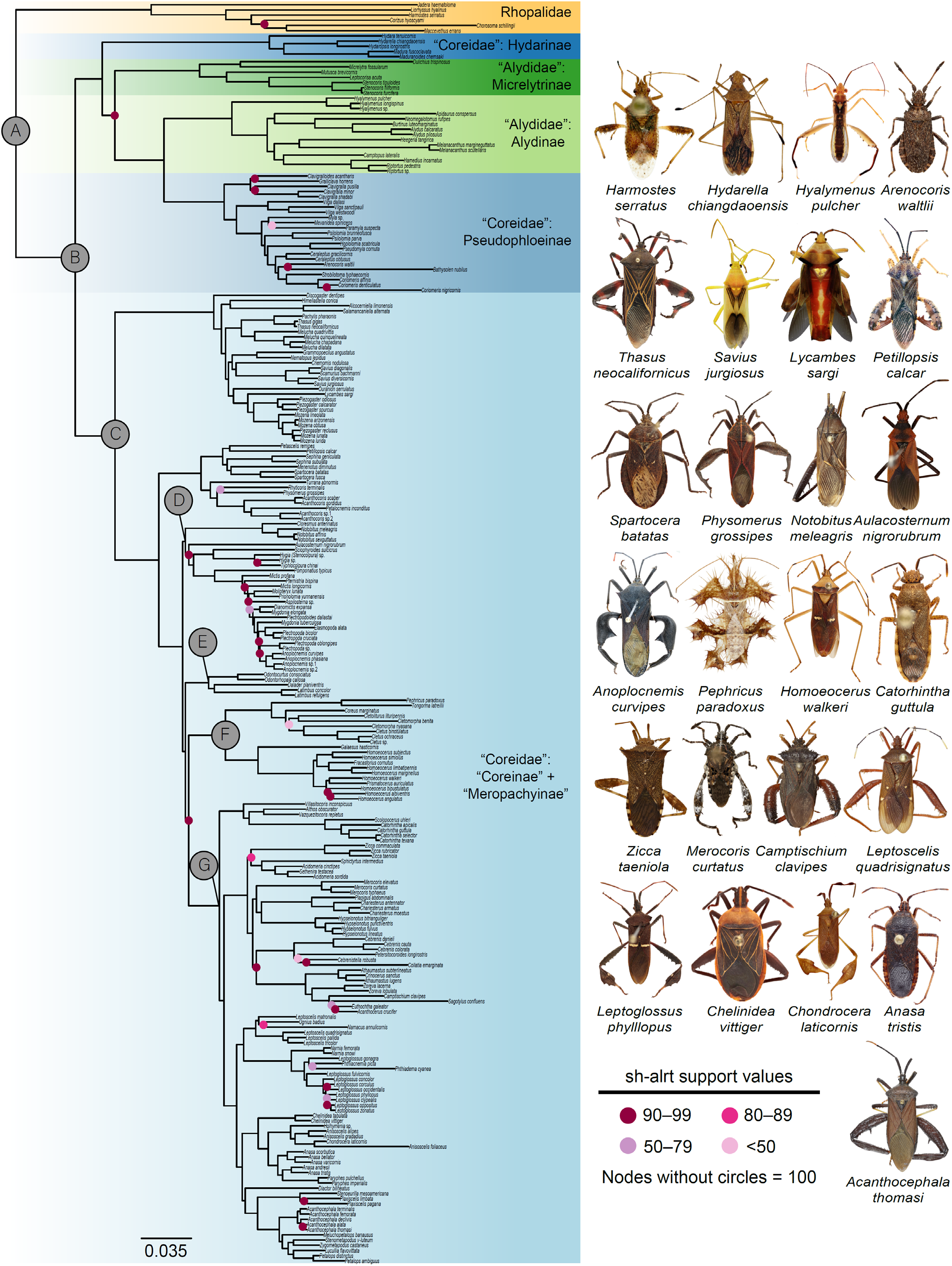
Maximum likelihood (ML) best tree based on the 50p concatenated alignment (outgroups not shown). Nodes labels A–G refer to clades discussed in the text. Colored circles at nodes represent instances when Shimodaira-Hasegawa-like approximate likelihood ratio test (sh-alrt) support is less than 100 (see Data Availability for tree with all terminals and sh-alrt and ultrafast bootstrap values visible). Dorsal habitus images of select species are given to show a range of diversity within the Coreoidea (images not to scale). The families Alydidae and Coreidae, as well as the subfamilies Coreinae and Meropachyinae and several tribes within them are, are not monophyletic.

### Diverse weapon trait combinations in the leaf-footed bugs

Plotting components at the terminals revealed a rich diversity of weapon trait combinations. Several clades include multiple species with a high number of weapon components (Figure 4). Processes off the ventral femora, such as spines and knobs (Component #6), were the most common, followed by the internal tibial processes. The presence of any weapon component was often accompanied by one or more ventral hind leg spines (Component #6), but these spines were not always accompanied by another component (e.g., *Anasa scorbutica* [Fabricius, 1775]) (Figure 4). Species with knobs or spines distributed on more than half of the ventral femoral surface also had more elaborated hind legs compared to species with a more restricted distribution of processes. The laterally expanded metathoracic acetabulum (Component #1), metapleural process (Component #2), and coxal and trochantal processes (Components #4 and #5, respectively) were some of the least common components. All thoracic (Components #1 and #2) and abdominal (Component #15) components were found paired with elaborated hind leg components, but not vice versa.

**Figure 4.**
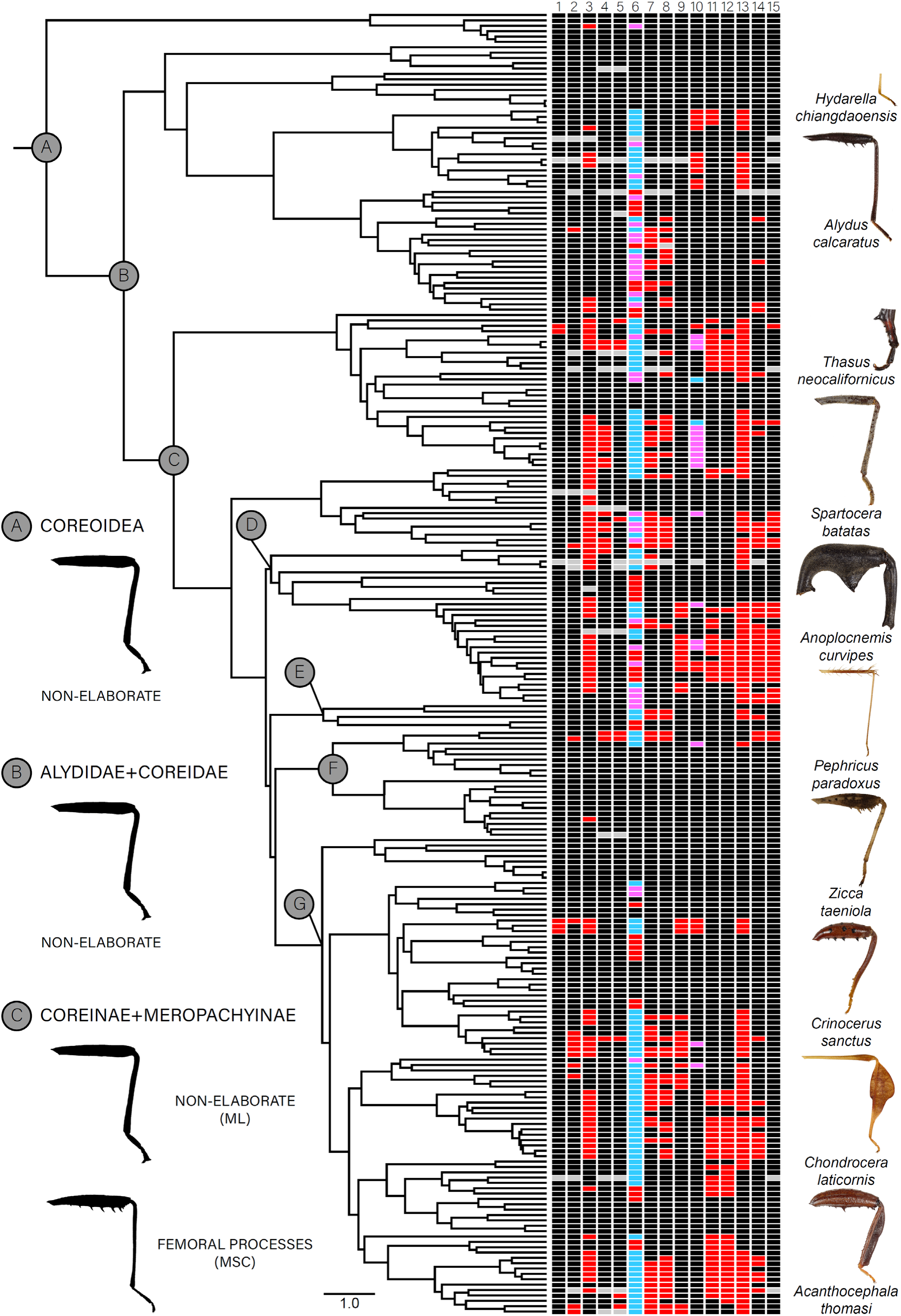
Ultrametric tree based on the 50p ML best tree, with components and component states displayed for terminal taxa on the right (images not to scale; State 0 = black, State 1 = red, State 2 = magenta, State 3 = blue, missing data = gray). Names of terminal taxa are removed for visualization purposes (refer to Figure 2 and Data Availability for terminal names of this particular topology). Nodes labels A–G refer to clades discussed in the text. For select nodes, an illustration representing the general male hind leg morphology based on Ancestral State Estimation (ASE) results is given; non-elaborate hind legs lack processes, flags, and curved femur and tibiae. In all ASE analyses, the last common ancestors of leaf-footed bugs (Nodes A and B) lacked elaborated weapon components on the thorax, hind legs, and abdomen. The last common ancestor of “Coreinae” + “Meropachyinae” (Node C) was similarly estimated to have non-elaborated structures or only with ventral processes on the femur, dependent on the tree topology used (ML or multispecies coalescent [MSC]). Across the phylogeny, there is a rich diversity of weapon trait combinations, with species in several clades expressing a high number of weapon components.

### All male weapon components are convergently evolved from “simple” structures

We found the metathorax, hind legs, and abdomen of the last common ancestors of leaf-footed bugs to consistently lack elaborated components across all ASE analyses (Figures 4, S8–S53). With respect to the metathorax and abdomen, this pattern was also true for the last common ancestor of Coreinae + Meropachyinae. Our ASE results based on the 50p ML ultrametric tree also did not estimate any elaborated components for the hind legs in the last common ancestor (Figures S11–S22). In contrast, ASE analyses based on the 50p and 50p25mi MSC ultrametric trees supported — albeit barely in the former case — the presence of processes on more than half of the ventral femoral surface (Component #6, State 3; Figures S29, S44; for other hind leg components, see Figures S26–S28, S30–S43, S45–S53).

Our ASE analyses found all components to be convergently evolved within leaf-footed bugs, regardless of data filtering and analytical approaches (Tables 2, S4; Figures 5, S8–S53). All weapon component states were reconstructed with at least two or more gains (minimum range = 2–19 gains), with the wider intercoxal distance (Component #3) and presence and distribution of femoral and tibial processes (Components #6, #13, and #14) having the most gains (Tables 2, S4). Losses or reductions were estimated for 12 components (minimum range = 1–26), with the presence and distribution of femoral processes and presence of tibial flags also exhibiting a high number of losses (Components #6 [State 3], #13, and #14) and outnumbering gains by about 2:1. All other components and component states had more gains than losses or slightly more losses than gains reconstructed in ASE analyses. We also evaluated whether branch lengths were associated with the number of evolutionary transitions. While some of the highest numbers of transitions occurred on relatively long branches (Figure 5A), in all other cases, long and short branches were associated with low to moderately high numbers of transitions, suggesting that these did not occur in a “clock-like” fashion.

**Figure 5.**
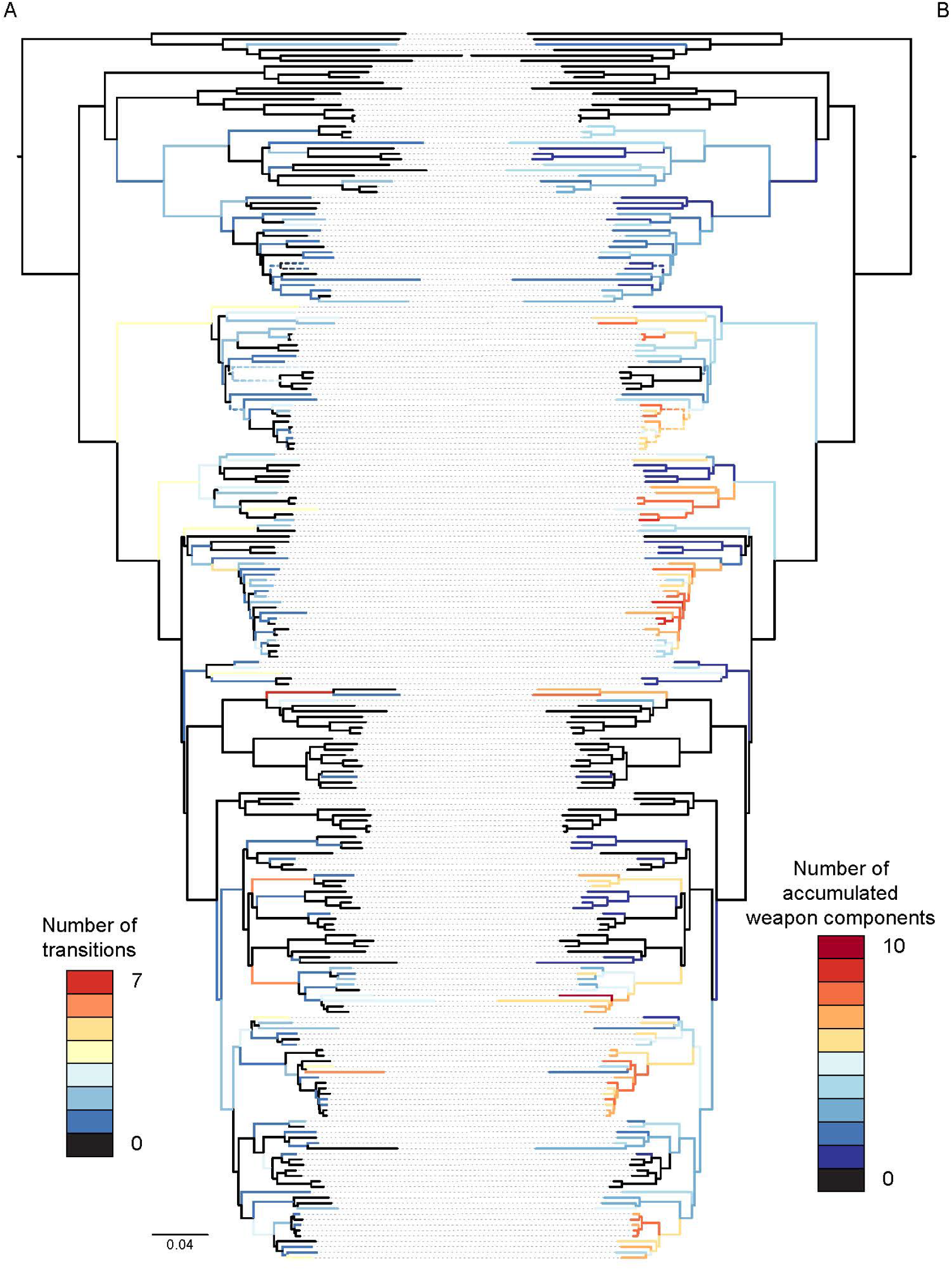
Summary of the total number of (A) inferred transitions (i.e., sum of total gains and losses) and (B) number of weapon states accumulated on branches of the 50p ML phylogram. Names of terminal taxa are removed for visualization purposes (refer to Figure 2 and Data Availability for terminal names of this particular topology). Dashed lines indicate branches affected by at least one component having an ambiguous ancestral state; in this case, a color gradient is given to represent the range of the total number of transitions or weapon states along the branch components. Weapon components generally accumulated along internal branches of several clades, but there were many instances of subsequent reductions weapon complexity over evolutionary time. In some cases, reductions in weapon components were followed by shifts back towards increasing the number of components.

We also tested the hypothesis that weapons evolve greater complexity over evolutionary time. Our results showed a general accumulation of weapon components along internal branches for several clades in leaf-footed bugs (Figure 5B). However, we observed about 50 instances of reduced weapon complexity, mostly at shallow scales of the phylogeny, with two lineages having lost weapon components entirely. In 20 cases, an initially more complex weapon began to exhibit reduced complexity in some clades, but then shifted back towards increasing complexity near the tips of the phylogeny (Figs. 5B, S54).

### A complex network of co-occurrences and correlations among weapon components

All weapon components were significantly associated with at least one other component in our co-occurrence analyses (Figure S55). A total of 57 positive and two negative associations were detected among components, with 35-component pairs not significantly associated with one another. We found that five components had ≥10 positive associations with other components on the thorax, hind legs, and abdomen; these were the intercoxal distance (Component #3), ventral and dorsal femoral processes (Components #6 and #8), and internal and external tibial processes (Components #13 and #14). The metapleural process (Component #2) and dorsal femoral processes (Component #8) were negatively associated with the internal tibial flag (Component #11) and femoral curvature (Component #9), respectively.

When accounting for phylogenetic non-independence among components, we found many of the co-occurring weapon components to be significantly correlated, albeit with some differences in the results of our binary correlated traits tests when using different phylogenetic topologies (Table S5; Figures 6, S56, S57); of the 57 positive *cooccur* component associations, 28–35 were significantly correlated. The metathoracic acetabulum (Component #1) was the only component not correlated with any other components, regardless of tree topology, as well as trochantal processes (Component #5) when using the ML tree. The intercoxal distance (Component #3), internal tibial processes (Component #13), and femoral processes (Components #6–#8), consistently had some of the highest number of correlations with other components, including with each other. In contrast, the metapleural process (Component #2), trochantal processes (Component #5; only MSC topologies), and tibial curvature (Component #10) had the lowest number of significant correlations.

**Figure 6.**
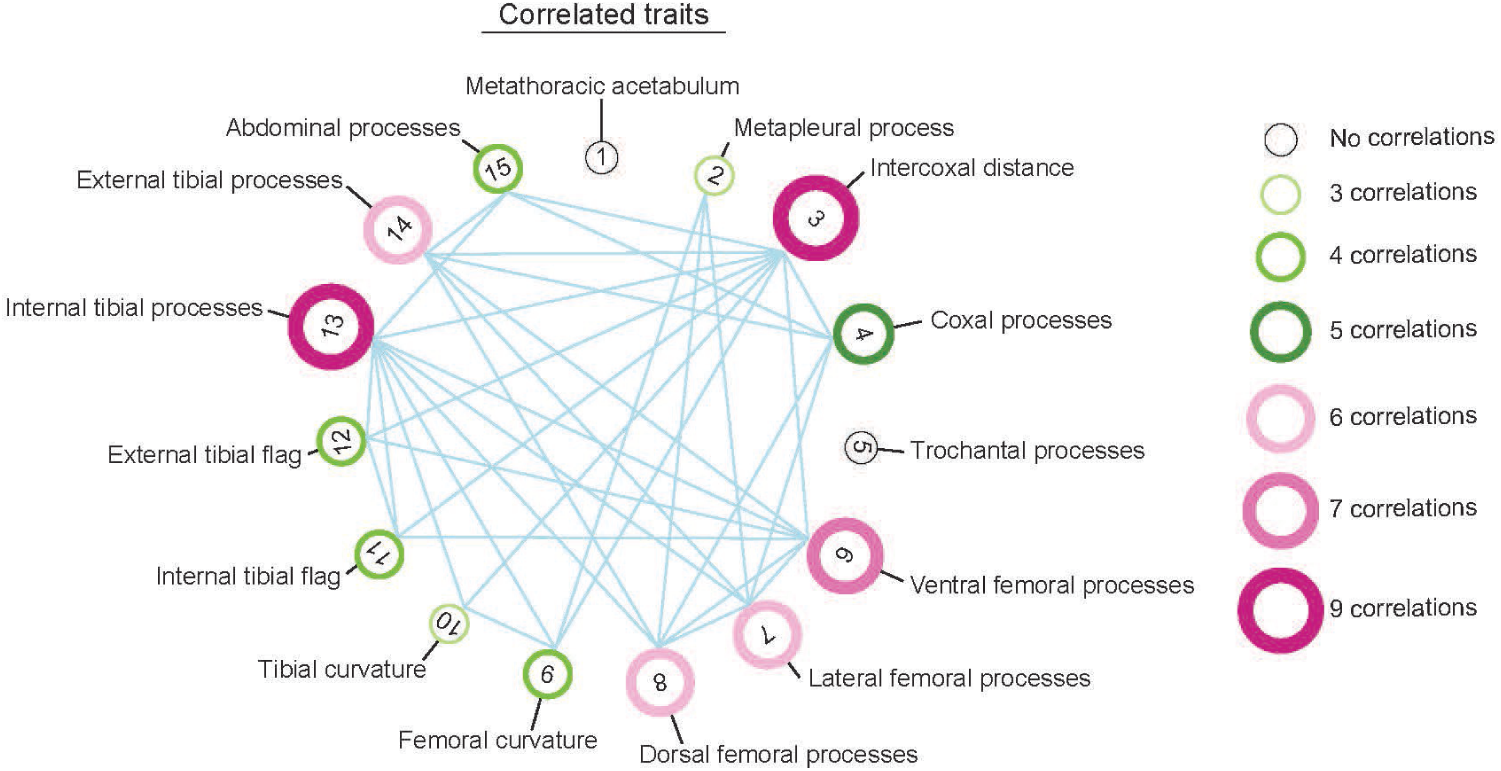
Correlated binary components based on the 50p ML ultrametric tree and trait co-occurrence results. The size and color of circles around component numbers reflects the total number of significant associations. Many of the weapon components examined are significantly correlated, with the intercoxal distance (#3), internal tibial processes (#13), and ventral femoral processes (#6) exhibiting a high number of correlations with other components.

## DISCUSSION

We found that the ancestor of leaf-footed bugs possessed simple hind legs and a streamlined body. Over time, morphological elaborations occurred, including sharp spines, flags, curves, and serrations. The number of these elaborations increased over time in many clades. Yet we also saw examples of reductions, sometimes followed by rapid elaboration again, implying a cyclical nature of weapon complexity. We found numerous instances of correlated evolution of weapon components, which allude to testable hypotheses on coordinated function during battle.

A breathtaking level of diversity and complexity unfurled from the simple, streamlined ancestor of leaf-footed bugs. Multiple discrete weapon components evolved independently a surprising number of times. For example, knobs and spines on the ventral side of the hind femur (ventral processes) arose independently on 21 occasions, becoming the most common weapon element. Curvature of the tibia away from the body evolved nine times independently, while a tibia curved towards the body arose four separate times. The lability in weapon components is remarkable and reminiscent of the extreme evolutionary modifications in the jaws of marine wrasses and freshwater cichlids (Wainwright et al. 2012). Other studies of animal weapons have suggested high lability in location, size, and general type (Emlen et al. 2005; Dalebout et al. 2008; Kim and Farrell 2015). Yet, few studies have examined the separate evolution of multiple weapon components. As an exception, Chow et al. (2021) examined the evolution of five components of the decapod claw across 107 species, finding five independent origins of snapping behavior and showing that snapping appendages can evolve via multiple evolutionary pathways. Similarly, we detected multiple configurations of weapon components associated with leaf-footed bug species known to engage in male-male combat (e.g., *Narnia femorata, Hyalymenus subinermis* Van Duzee, 1923*, Leptoglossus gonagra* [Fabricius, 1775]*, Mictis longicornis*; Figures 2, 4).

Our results support the hypothesis that greater weapon complexity evolved over time. Patterns of increasing weaponization have been previous hypothesized (e.g., Emlen 2008; Moore et al. 2022), but they have very rarely been tested using phylogenetic analyses. Across taxa, early animal weapons were likely small, sharp extensions such as spines, spurs, and fangs (Emlen 2008). In the case of leaf-footed bugs, weapon elaboration started with structures bulging and projecting out of the ventral side of the hind femur (Figure 2). The addition of novel weapon components or the elaboration of existing components may provide an advantage in signaling a male’s fighting prowess (Clutton-Brock et al. 1979; Searcy and Nowicki 2010), or it may directly yield a fighting advantage, for example, allowing a male to better grasp another male’s body part more effectively during battle (e.g., tubercles used in beetles that clamp, Eberhard 1979; and fiddler crabs that grip, Dennenmoser and Christy 2013).

The leaf-footed bugs experienced a proliferation of weapon components over time, but losses and reductions of these weapon components were also abundant. For example, *Petillopsis calcar* (Dallas, 1852) shows seven weaponized components, while its close relative, *Sephina geniculata* Distant, 1881, possesses only one (Figures 4, 5, S8–S53). *Camptischium clavipes* (Fabricius, 1803) exhibits ten weaponized components, while its close relative, *Zoreva lacerna* Brailovsky & Barrera, 1982, has two (Figures 4, 5, S8–S53). Looking across the phylogeny, the ventral side of the hind femur showed 21 independent gains of weaponized components followed by 26 losses or reductions in descendent taxa. Curvature of the tibia away from the body was lost once, and curvature of the tibia towards the body was lost twice. The fact that some components were lost more often than others suggests that they may have been less functionally integrated with other components. Similarly, they may have provided fewer fitness benefits or greater costs, perhaps due to biomechanical compromises or energetic demands.

Weapons and other morphological elaborations cannot continue to become larger and more complex indefinitely. Even a modest weapon may be associated with costs that outweigh the benefits in some contexts (Miller and Svensson 2014). For example, weapons may have high physiological demands during development, maintenance, or use (Basolo and Alcaraz 2003; Somjee et al. 2018; O’Brien et al. 2019) but, see (Kotiaho 2001; McCullough et al. 2012; McCullough and Emlen 2013). Thus, times of resource scarcity (Boggs 2009) or heightened parasite loads (Hamilton and Zuk 1982) may increase the costs and trade-offs associated with investing in a complex morphological structure. In those cases, males with fewer or smaller components may achieve higher fitness than those investing in large size and complex structures (Brockmann 2001; Emlen 2014). Additionally, changes in predator abundance may raise the risks of predation for males that invest in certain bulky or conspicuous traits, thus selecting against some forms of elaboration (Møller 1996; but, see Lane 2018; Metz et al. 2018). Further, given that these structures are likely often important in sexual selection, changes in resource distribution across the landscape and over time may make females more dispersed and less defensible (Emlen and Oring 1977), reducing the benefits of weapon investment (Lüpold et al. 2014). Considering the many scenarios that should favor reduced weaponization, it is perhaps not surprising that weapon losses are so common. Indeed, the loss of sexually selected components have been documented in stalk-eyed flies, dung beetles, birds, artiodactyls, and more (Baker and Wilkinson 2001; Wiens 2001; Caro et al. 2003; Kimball et al. 2011; Kim and Farrell 2015; Chow et al. 2021; Menezes and Palaoro 2022). Interestingly, in leaf-footed bugs, a reduction in weapon complexity was, in some cases, followed by increasing complexity near the tips of the phylogeny, suggesting a cyclical nature to weapon elaboration in clades that are primed for it (Emlen 2014; Emlen, *personal communication*). In those cases, the return of favorable conditions after a weapon loss or reduction may quickly select on a small knob, spine, and flag for expansion, leading to the regain of weapon components in a lineage.

Weapon forms can be associated with specific fighting styles (Geist 1966; Eberhard 1980; Lundrigan 1996; Caro et al. 2003). The high plasticity and lability of behavior (West-Eberhard 2003) suggests that changes in fighting style may take the lead in evolution, with morphology to follow (Emlen 2008). The questions of why male fighting styles initially change and how such changes are retained are largely unaddressed. For many species, the structural context in which fights occur may be central. For example, competitions that occur in flat open spaces should take a different form than competitions that occur in tight burrows or dense vegetation (Eberhard 1980; Emlen 2008; Cabrera and Stankowich 2020). Clades of animal species where males fight in a variety of structural contexts provide outstanding opportunities to investigate the role of the arena in the alteration of fighting behaviors. A wide range of host plant species is used by the ∼3300 species of leaf-footed bugs (Schaefer and Mitchell 1983; Mitchell 2000). Thus, males fight upon many different surfaces, such as the smooth shafts of bamboo (Miyatake 1995), spiny cacti (Procter et al. 2012), or leafy, flexible legumes (Tatarnic and Spence 2013). In some cases, a single leaf-footed bug species can use a wealth of host plants. For example, the well-armed Florida leaf-footed bug, *Acanthocephala femorata* (Fabricius, 1775), competes over territories on plants as strikingly different in structure as sunflower (*Helianthus annus*), white goosefoot (*Chenopodium album*), and yellow thistle (*Cirsium horridulum*) (Baranowski and Slater 1986; Author, personal observation). Fighting surfaces should influence surface grip, and the structure of the host plant will affect the space available for combat maneuvers. It would be fascinating to study selection on weapon components in *A. femorata* or another species that uses multiple host plants.

All biological motion is subject to the laws of physics. As a result, mechanics and evolution are inescapably linked. Mechanical function shapes evolution; certain components of a weapon should thus be expected to correlate with other components, and together they should function in an integrated manner (Chow et al. 2021; Nogueira et al. 2022). Palaoro and Peixoto (2022) recently called for studies to move away from simplified measures of weapons and focus on better understanding of weapon functionality in less-studied taxa. Here, we detected 86 binary (presence/absence) trait combinations in 248 leaf-footed bugs. A network of evolutionary associations emerged from our analyses (Figure 6). Correlations among components may indicate pleiotropy or linkage disequilibrium shaping the pathways of weapon evolution. Further, the correlations suggest testable hypotheses of biomechanical function and integration. For example, when tibia curve (Component #10), it is typically away from the body, often with one or more prominent internal spines (Component #13; e.g., see tibia in Figure 2F). The curved, spined tibia are associated with a curved femur (Component #9), which may act as a catching arch to help hold the opponent in place while the tibial spines pierce into its body. We also found that the metapleural process (Component #2), a spine that emerges laterally from the thorax, is found in species with curved femurs with dorsal or lateral projections (Figures 2, 6), though how these structures would function together will remain unclear until behavioral analysis is pursued.

Evolutionary associations also highlight individual components that may be at the heart of a functional weapon in this superfamily. The two components with the greatest number of correlations with other components were an increased intercoxal distance (Component #3)— which is somewhat akin to broad shoulders in humans—, and bumps, knobs, or spines on the inside of the tibia (Component #13), which may be used for grip and/or as a concentrated force point during squeezing (Figure 6). Increased intercoxal distance was one of several traits that we included in our analyses without prior direct evidence that it used in or contributes to success in aggressive interactions. The substantial number of correlations with other components suggest that it may be part of the crucial morphological machinery of leaf-footed bug weapons (see also Okada et al. 2012). Our hope is that the patterns revealed in this work spark behavioral studies and biomechanical analyses for many years to come.

## Future work

Animal weapons provide a wealth of opportunities to understand the evolution of complexity, assembly, and integration. By examining 15 components of a multi-component weapon system, we were able to reconstruct the remarkable and dynamic evolution of weapon form in the leaf-footed bug superfamily. An exciting next step is to test the extent to which fighting behavior shapes the evolution of weapon morphologies (Lundrigan 1996). Phylogenetic comparative studies will be helpful for testing evolutionary correlations between fighting techniques and specific weapon forms (Caro et al. 2003; Chow et al. 2021). Microevolutionary work should be especially powerful in this line of inquiry. By manipulating the micro-habitat where fighting takes place, researchers may alter fighting style and examine changes in selection on components of the sexually selected weapon. Studies using experimental evolution could subsequently determine if such changes in behavior and selection lead to evolutionary changes in the morphology of the weapon. Future work on animal weapons should also consider that weapon diversity encompasses more than just external form. For example, weapons have been selected to function in physical contests, thus studies of weapon evolution should examine the structure and material properties of weaponized components. Across (Swanson et al. 2013), and even within species (Woodman et al. 2021), weapons can vary in their ability to resist the rigors of combat. Weapon structural integrity should evolve alongside fighting style (McCullough et al. 2014), skill (Briffa and Lane 2017), and the forces applied during combat (Lailvaux and Irschick 2006). A large or complex weapon is of little use if it is easily destroyed during battle.

## Acknowledgements

We thank E.V. Ginny Greenway, David Labonte, and Douglas Emlen for providing helpful feedback and for invigorating discussions. Christiane Weirauch, Dimitri Forero, Paul Masonick, Joe Eger, Petr Kment, Marcos Roca-Cusachs, Vasily Grebennikov, Takahisa Miyatake, Nik Tatarnic, Jason Cryan, Mark Deyrup, Wei Song Hwang, Li You, Oliver Keller, John Leavengood, Wanessa da Silva Costa, Sam Noble Oklahoma Museum of Natural History, Field Museum of Natural History, Florida State Collection of Arthropods, California Academy of Sciences, and National Museums of Kenya contributed specimens. We thank Bob McCleery, Cebisile N. Magagula, and the Savannah Research Center for facilitating specimen collection in Eswatini. South African specimens were collected under Ezemvelo KZN Wildlife permit #OP 172/2018 and the iSimangaliso Wetland Park Authority (World Heritage Site). Additional samples were collected with funding by University of Florida Research Abroad for Doctoral Students, Lee Kong Chian Natural History Museum fellowship, and National Science Foundation (NSF) OISE-1614015 to Zach Emberts (Australia permit #01-000204-1; Singapore permit #NP/RP17-012); the Society of the Study of Evolution Rosemary Grant; Smithsonian Tropical Research Institute Short Term Fellowship; and Systematics, Evolution, and Biodiversity Endowment Award (Entomological Society of America) to Ummat Somjee; and NSF DEB-0542864 to Michael Sharkey and Brian Brown. Nathan Friedman, Caroline Miller, and Min Zhao assisted with molecular benchwork.

---

**Table S1.** Taxon sampling summary.

**Table S2.** Sequence data summary. Abbreviations: bp, base pairs; EtOH, ethanol; Max., maximum; Min., minimum; UCE, ultraconserved element.

**Table S3.** Summary of informative sites and number of UCE loci in datasets generated in this study.

**Table S4.** Summary of component state gains and losses across different ultrametric trees (component state 0 not reported). Abbreviations: ARD, all rates different model; ER, equal rates model; G–L, gains minus losses; Min, minimum; ML, maximum likelihood; MSC, multispecies coalescent; N, sample size; SYM, symmetric model; 50p, alignments comprised of locus alignments with at least 50% of the total taxa sampled in this study; 50p25mi, 25% most informative loci subsampled from the 50p dataset.

**Table S5.** Results of Pagel’s (1994) binary correlated traits test for each pairwise comparison of components found to have significant co-occurrence (from co-occur results) based on the 50p maximum likelihood (ML), 50p multispecies coalescent (MSC), and 50p25mi MSC ultrametric trees. Significant p-values are bolded. Abbreviations: AIC(D), Akaike Information Criterion for dependent model; AIC(I), Akaike Information Criterion for independent model; ARD, all rates different model; BH, Benjamini-Hochberg correction for multiple comparisons (Benjamini & Hochberg, 1995); ER, equal rates model; LR, likelihood ratio; NT, not tested (due to potential unreplicated effects based on ancestral state estimation of components; see main text for more details).

**Figure S1.** Maximum likelihood (ML) best tree based on the 50p concatenated alignment. Circles at nodes represent instances when Shimodaira-Hasegawa-like approximate likelihood ratio test (sh-alrt) and/or bootstrap support are less than 100.

**Figure S2.** 70p ML best tree. Circles at nodes represent instances when sh-alrt and/or bootstrap support are less than 100.

**Figure S3.** Multispecies coalescent (MSC) species tree based on 50p gene trees (displayed as a cladogram). Circles at nodes represent local posterior probabilities (LPPs) less than 1.

**Figure S4.** 70p MSC species tree (displayed as a cladogram). Circles at nodes represent LPPs less than 1.

**Figure S5**. 50p25mi MSC species tree (displayed as a cladogram). Circles at nodes represent LPPs less than 1.

**Figure S6.** 70p25mi MSC species tree (displayed as a cladogram). Circles at nodes represent LPPs less than 1.

**Figure S7.** Evolutionary scenarios between hypothetical Components A and B on the same phylogeny. State 0 (“absent”) is shown in black lines, and State 1 (“present”) is shown in red. (A) Examples of Components A and B showing replicated evolutionary patterns in clades that are in relatively close proximity to one another on the phylogenetic tree. According to Maddison & FitzJohn (2015), the rate of origins in Component B could potentially be explained by a third, unrelated trait that changed in the larger clade marked by the red asterisk. (B) A scenario in which a correlated traits test of Components A and B finds a significant evolutionary model in which Component A depends on the evolutionary origins of Component B. However, when evaluating the evolutionary origins of Components A and B based on ancestral state estimates, Component A generally evolves first, with Component B evolving later in the same clades; this does not support prediction of the significant model. Thus, the evolutionary patterns might alternatively suggest Component B to be dependent on Component A, but this alternative model is not supported by the correlated traits test. As such, the correlated traits test is considered to have resulted in a false positive for the Component A dependency model, and the two components are not considered correlated in the absence of a statistically significant, alternative dependency model.

**Figure S8.** Summary of gains and losses across all male hind leg components. The 50p ML ultrametric tree is displayed as a cladogram, and sister taxa with the same sets of component states have been collapsed into a single terminal for visualization. For simplicity in summarizing general component transitions, multistate Characters #6 and #10 are shown as binary states.

**Figure S9–S23.** Ancestral state estimates (ASE) based on the 50p ML ultrametric tree for Components #1–#15, with each component’s corresponding best model of evolution (equal rates [ER] or all rates different [ARD]). Taxa with missing data for a given component are pruned from the tree for analysis. Pie charts show the likeliest states for a given node (State 0 = black, State 1 = red, State 2 = magenta, State 3 = blue), with branches similarly colored to represent the most likely state.

**Figure S24–S38.** ASE based on the 50p MSC ultrametric tree for Components #1–#15, with each component’s corresponding best model of evolution (ER or ARD). Taxa with missing data for a given component are pruned from the tree for analysis. Pie charts show the likeliest states for a given node (State 0 = black, State 1 = red, State 2 = magenta, State 3 = blue), with branches similarly colored to represent the most likely state.

**Figure S39–S53.** ASE based on the 50p25mi MSC ultrametric tree for Components #1–#15, with each component’s corresponding best model of evolution (ER or ARD). Taxa with missing data for a given component are pruned from the tree for analysis. Pie charts show the likeliest states for a given node (State 0 = black, State 1 = red, State 2 = magenta, State 3 = blue), with branches similarly colored to represent the most likely state.

**Figure S54.** Summary of the total number of components accumulated on branches of the 50p ML phylogram, with insets at the right highlight areas of the tree where cyclical patterns of increasing and decreasing elaborations were observed (each instance indicated by a number and thickened branches). Dashed lines indicate branches affected by at least one component having an ambiguous ancestral state; in this case, a color gradient is given to represent the range of the total number of components along the branch. Branches denoted with an asterisk in insets at the right have been modified (i.e., arbitrarily lengthened and not to scale) for visualization.

**Figure S55.** Trait co-occurrences across 15 weapon components. Components with significant positive associations are shown in solid blue lines, while those with negative associations are shown in dashed yellow lines. The size of circles around component numbers reflects the total number of significant associations.

**Figure S56.** Correlated components based on the 50p MSC ultrametric tree and trait co-occurrence results. The size of circles around component numbers reflects the total number of significant associations.

**Figure S57.** Correlated components based on the 50p25mi MSC ultrametric tree and trait co-occurrence results. The size of circles around component numbers reflects the total number of significant associations.

