## Supplementary Information for "The evolution of multi-component weapons in the superfamily of leaf-footed bugs"

Supplementary methods

*Molecular data collection*

For 216 taxa, we retrieved UCE sequence capture data from Kieran et al. (2019); Forthman, Miller et al. (2019, 2020, 2022); Emberts et al. (2020); and Miller et al. (2022) (Table S1). We also downloaded genome sequences of *Halyomorpha halys* (Pentatomidae) and *Oncopeltus fasciatus* (Lygaeidae) from NCBI to extract UCE sequences from scaffolds.

We generated new sequence data for 30 taxa, 12 of which were preserved in fresh ethanol (EtOH) while the remaining 18 taxa were dried, pinned museum samples (Table S1). Genomic DNA was extracted from the entire body or any part of the body (e.g., thorax, abdomen, and/or legs) to sample similar amounts of tissue across specimens. For the EtOH preserved specimens, DNA was extracted using a Qiagen DNeasy Blood and Tissue kit ("DNeasy"), but the manufacturer's protocol was modified so that tissue was incubated for 24 hr in a solution of 180 µL Buffer ATL and 20 µL proteinase K, and DNA was eluted twice with 50 µL Buffer AE. For the dried specimens, DNA was extracted with a DNeasy kit coupled with a Qiagen QIAquick PCR purification kit ("DNQIA"; Knyshov et al., 2019; Forthman, Miller et al., 2020, 2022). The DNQIA protocol followed the DNeasy protocol described above but with the following modifications: a QIAquick spin column was used, the AW washes were replaced with Buffer PE, and the DNA was eluted twice with 50 µL Buffer EB. DNA quality and quantity was evaluated using 1% agarose gel electrophoresis and a Qubit 2.0 fluorometer, respectively. Where possible, samples were normalized (10–20 ng/µL), and high molecular weight samples were fragmented into 200–1000 bp using a Covaris M220 Focused-ultrasonicator for 60 s. For DNA extracted from dried specimens, a PreCR Repair Mix kit with a 3X SPRI clean-up was used to repair DNA.

Libraries were made following the KAPA Hyper Prep Kit protocol with modifications. Half volume reactions were used for all steps. DNA samples were ligated with iTru universal adapter stubs and 8 bp dual indexes (Glenn et al., 2019). Library amplification was performed with initial denaturation at 98°C for 3 min; followed by 14–16 cycles of 98°C for 30 s, 60°C for 30 s, and 72°C for 30 s; and final extension at 72°C for 5 min. The quality and quantity of amplified libraries were evaluated by gel electrophoresis and Qubit, respectively, before combining into 1000 ng pools in equimolar amounts, dried at 60°C, and resuspended in 14 µL IDTE.

Target enrichment was done using baits designed from two pentatomomorphan taxa (Faircloth, 2017; Forthman et al., 2019). A hybridization mixture with 1/2 volume of baits for EtOH preserved samples and 1/4 volume of baits for dried samples was used. Baits were hybridized with libraries following Forthman, Miller et al.'s (2022) touchdown capture protocol: 65°C for 12 hours, 62°C for 12 hours, and 60°C for 12 hours. Bait-target hybrids were bound to Dynabeads M-280 Streptavidin beads, washed four times at 60°C, and resuspended in 30 µL IDTE. Post-capture amplification was performed using a PCR amplification mix containing 2.5 µL each of 5 µM iTru P5/P7 primers (Glenn et al., 2019) and 14–18 amplification cycles following manufacturer's protocol, except an annealing temperature of 60°C and an extension period of 45 s were used. Post-amplified enrichments were cleaned with Hydrophobic Sera-Mag SpeedBeads Carboxyl Magnetic Beads and two washes in 70% EtOH. Enriched library pools were resuspended in 22 µL IDTE, quantified with Qubit, and pooled into a single pool in equimolar amounts prior to sequencing on a single Illumina HiSeq3000 lane (2x100) at the University of Florida's Interdisciplinary Center for Biotechnology Research.

Default settings were used for all data processing steps described below unless otherwise stated. Illumiprocessor v2.0 (Faircloth, 2013) was used to trim adapters from demultiplexed, raw sequence reads. Duplicate reads were discarded using PRINSEQ-lite v0.20.4 (Schmieder & Edwards, 2011). QuorUM v1.1.0 (Marçais et al., 2015) was then used to error-correct the remaining reads. Filtered reads were *de novo* assembled using SPAdes v3.13.0 (single-cell and auto coverage cutoff options; Nurk et al., 2013). We used PHYLUCE v1.7.0 (Faircloth, 2016) to extract UCE loci from assembled contigs following Forthman et al. (2019, 2020, 2022b). We also used PHYLUCE to align UCE baits to two genome sequences (*H. halys* and *O. fasciatus*) and extract UCE loci with 500 bp of flanking nucleotides. A summary regarding newly generated read, contig, and UCE data are given in Table S2.

Loci were aligned individually with PHYLUCE using the following settings: mafft alignment (--mafft; Katoh et al., 2002; Katoh & Standley, 2013), generate incomplete matrices (--incomplete-matrix), no alignment trimming (--no-trim), and allow nucleotide uncertainty (--ambiguous). Locus alignments were trimmed using trimAl v1.2 (heuristic automated1 method; Capella-Gutiérrez et al., 2009). Locus alignments with at least 50% and 70% of the total taxa were selected for analysis (referred to as "50p" and "70p" datasets, respectively). We also subsampled each of these datasets for the 25% most parsimony-informative loci (referred to as "25mi"), resulting in four datasets: 50p, 50p25mi, 70p, 70p25mi (see Table S3 for a summary of informative sites and number of UCE loci in each dataset).

*Phylogenetic inferences*

For the 50p and 70p datasets, we concatenated locus alignments for maximum likelihood (ML) phylogenetic analysis. We used IQ-Tree v2.1.2 (Minh et al., 2020) to determine the best model of sequence evolution and partitioning scheme of loci with the following settings: -m MF+MERGE (Kalyaanamoorthy et al., 2017), -rcluster 10, and -mrate E,I,G,R; we excluded I+G from model selection since these parameters are not independent of each other (Sullivan et al., 1999; Yang, 2006). For each of these datasets, ten separate partitioned ML analyses (Chernomor et al., 2016) were performed in IQ-Tree, with support measured by 1000 ultrafast bootstrap replicates optimized by nearest neighbor interchange based on bootstrap alignments (-B 1000 -bnni; Hoang et al., 2018) and 1000 Shimodaira-Hasegawa-like approximate likelihood ratio test (sh-alrt) replicates (-alrt 1000; Guindon et al., 2010). The tree with the best log-likelihood was selected for each dataset.

To account for gene tree discordance due to incomplete lineage sorting, we also inferred species trees for all four datasets under the multispecies coalescent (MSC) model (Kubatko & Degnan, 2007; Degnan & Rosenberg, 2006, 2009; Roch & Steel, 2015). We included the

50p25mi and 70p25mi datasets for species tree inference given that filtering for more informative loci has been shown to improve topological and branch length (in coalescent units) estimates in summary coalescent analyses (Mirarab, Bayzid et al., 2014; Hosner et al., 2016; Meiklejohn et al., 2016; Sayyari & Mirarab, 2016; Sayyari et al., 2017; Forthman, Braun et al., 2022). We estimated the best-fit model of sequence evolution (-mrate E,I,G,R) for each locus alignment using IQ-Tree. Gene trees were then estimated (-m MFP) with near-zero branch lengths collapsed (-czb) as this has also been shown to improve species tree inferences (Zhang et al., 2017; Forthman, Braun et al., 2022). Species trees were inferred from optimal gene trees using ASTRAL-III v5.7.7 (Mirarab, Reaz et al., 2014; Sayyari & Mirarab, 2016; Zhang et al., 2018). We assessed clade support using local posterior probabilities (Sayyari & Mirarab, 2016).

For ASE analyses, we transformed our 50p ML and 50p and 50p25mi MSC trees into ultrametric trees. First, we used IQ-Tree to estimate branch lengths as units of substitutions on the 50p and 50p25mi MSC topologies. We pruned outgroup taxa and used the *chronos* function in the *ape* package v5.6.1 (Paradis & Schliep, 2019) with R v4.1.2 (R Core Team, 2021) to generate ultrametric trees for the ML and MSC phylograms. We tested four models (correlated, discrete, relaxed, clock) and four values of lambda (0, 0.1, 1, 10) with the root node calibrated to a relative age of 1. While the ultrametric tree with the highest likelihood was generated under a correlated model when lambda equaled 0, the relative branch length distributions did not accurately reflect the distributions of branch lengths in their corresponding phylograms. After further inspection, we found the correlated model with lambda set to 0.1 produced ultrametric trees that better represented relative branch length distributions with little difference in the penalized log-likelihood values; thus, we selected these trees for ancestral state estimation analyses.

### Table S1. Taxon sampling summary.

| Family | Subfamily | Tribe | Genus | Species | Reference | NCBI accession |
|---|---|---|---|---|---|---|
| Alydidae | Alydinae | | *Alydus* | *calcaratus* | Forthman et al. (2022) | SRR16596614 |
| Alydidae | Alydinae | | *Alydus* | *pilosulus* | Emberts et al. (2020) | SRR11213492 |
| Alydidae | Alydinae | | *Apidaurus* | *conspersus* | Forthman et al. (2022) | SRR16596602 |
| Alydidae | Alydinae | | *Burtinus* | *luteomarginatus* | Forthman et al. (2022) | SRR16596569 |
| Alydidae | Alydinae | | *Camptopus* | *lateralis* | Forthman et al. (2022) | SRR16596566 |
| Alydidae | Alydinae | | *Hamedius* | *incarnatus* | Forthman et al. (2019) | SRR8903492 |
| Alydidae | Alydinae | | *Heegeria* | *tangirica* | Forthman et al. (2022) | SRR16596599 |
| Alydidae | Alydinae | | *Hyalymenus* | *longispinus* | Forthman et al. (2019) | SRR8903490 |
| Alydidae | Alydinae | | *Hyalymenus* | *pulcher* | Forthman et al. (2022) | SRR16596597 |
| Alydidae | Alydinae | | *Hyalymenus* | sp. | Forthman et al. (2022) | SRR16596595 |
| Alydidae | Alydinae | | *Melanacanthus* | *margineguttatus* | Emberts et al. (2020) | SRR11213508 |
| Alydidae | Alydinae | | *Melanacanthus* | *scutellaris* | Emberts et al. (2020) | SRR11213507 |
| Alydidae | Alydinae | | *Neomegalotomus* | *rufipes* | Forthman et al. (2019) | SRR8903491 |
| Alydidae | Alydinae | | *Riptortus* | *pedestris* | Emberts et al. (2020) | SRR11213489 |
| Alydidae | Alydinae | | *Riptortus* | sp. | Forthman et al. (2022) | SRR16596574 |
| Alydidae | Micrelytrinae | Leptocorisini | *Leptocorisa* | *acuta* | Emberts et al. (2020) | SRR11213505 |
| Alydidae | Micrelytrinae | Leptocorisini | *Mutusca* | *brevicornis* | Forthman et al. (2019) | SRR8903496 |
| Alydidae | Micrelytrinae | Leptocorisini | *Stenocoris* | *filiformis* | Emberts et al. (2020) | SRR11213500 |
| Alydidae | Micrelytrinae | Leptocorisini | *Stenocoris* | *furcifera* | Emberts et al. (2020) | SRR11213488 |
| Alydidae | Micrelytrinae | Leptocorisini | *Stenocoris* | *tipuloides* | Forthman et al. (2019) | SRR8903489 |
| Alydidae | Micrelytrinae | Micrelytrini | *Dulichius* | *trispinosus* | Forthman et al. (2022) | SRR16596603 |
| Alydidae | Micrelytrinae | Micrelytrini | *Micrelytra* | *fossularum* | Forthman et al. (2022) | SRR16596583 |
| Coreidae | Coreinae | Acanthocephalini | *Acanthocephala* | *alata* | Forthman et al. (2020) | SRR9224639 |
| Coreidae | Coreinae | Acanthocephalini | *Acanthocephala* | *declivis* | Emberts et al. (2020) | SRR11213518 |
| Coreidae | Coreinae | Acanthocephalini | *Acanthocephala* | *femorata* | Kieran et al. (2019) | SRR7819299 |
| Coreidae | Coreinae | Acanthocephalini | *Acanthocephala* | *terminalis* | Emberts et al. (2020) | SRR11213487 |
| Coreidae | Coreinae | Acanthocephalini | *Acanthocephala* | *thomasi* | Kieran et al. (2019) | SRR7819302 |
| Coreidae | Coreinae | Acanthocephalini | *Lucullia* | *flavovittata* | Miller et al. (2022) | SRR23782870 |
| Coreidae | Coreinae | Acanthocephalini | *Meluchopetalops* | *banasus* | Miller et al. (2022) | SRR23782887 |
| Coreidae | Coreinae | Acanthocephalini | *Petalops* | *ambiguus* | Miller et al. (2022) | SRR23782862 |
| Coreidae | Coreinae | Acanthocephalini | *Petalops* | *distinctus* | Forthman et al. (2020) | SRR9224645 |
| Coreidae | Coreinae | Acanthocephalini | *Stenometapodus* | *v-luteum* | Forthman et al. (2020) | SRR9224643 |
| Coreidae | Coreinae | Acanthocephalini | *Zygometapodus* | *castaneus* | Forthman et al. (2020) | SRR9224642 |
| Coreidae | Coreinae | Acanthocerini | *Athaumastus* | *lugens* | | SRR23782915 |
| Coreidae | Coreinae | Acanthocerini | *Athaumastus* | *subterlineatus* | Forthman et al. (2020) | SRR9224636 |
| Coreidae | Coreinae | Acanthocerini | *Camptischium* | *clavipes* | Forthman et al. (2020) | SRR9224650 |
| Coreidae | Coreinae | Acanthocerini | *Crinocerus* | *sanctus* | Forthman et al. (2020) | SRR9224651 |
| Coreidae | Coreinae | Acanthocerini | *Euthochtha* | *galeator* | Forthman et al. (2020) | SRR9224652 |
| Coreidae | Coreinae | Acanthocerini | *Sagotylus* | *confluens* | Forthman et al. (2020) | SRR23782912 |
| Coreidae | Coreinae | Acanthocerini | *Zoreva* | *lacerna* | Forthman et al. (2020) | SRR9224653 |
| Coreidae | Coreinae | Acanthocerini | *Zoreva* | *lobulata* | Forthman et al. (2020) | SRR9224646 |
| Coreidae | Coreinae | Acanthocorini | *Acanthocerus* | *crucifer* | Forthman et al. (2020) | SRR9224637 |
| Coreidae | Coreinae | Acanthocorini | *Acanthocoris* | *scaber* | Emberts et al. (2020) | SRR11213517 |
| Coreidae | Coreinae | Acanthocorini | *Acanthocoris* | *sordidus* | Forthman et al. (2020) | SRR9224690 |
| Coreidae | Coreinae | Acanthocorini | *Acanthocoris* | sp.1 | Emberts et al. (2020) | SRR11213495 |
| Coreidae | Coreinae | Acanthocorini | *Acanthocoris* | sp.2 | Forthman et al. (2020) | SRR9224693 |
| Coreidae | Coreinae | Acanthocorini | *Petalocnemis* | *inconditus* | Miller et al. (2022) | SRR23782863 |
| Coreidae | Coreinae | Acanthocorini | *Physomerus* | *grossipes* | Emberts et al. (2020) | SRR11213509 |
| Coreidae | Coreinae | Acanthocorini | *Pomponatius* | *typicus* | | SRR23782913 |
| Coreidae | Coreinae | Acanthocorini | *Rhyticoris* | *terminalis* | Forthman et al. (2020) | SRR9224688 |
| Coreidae | Coreinae | Acanthocorini | *Turrana* | *abnormis* | | SRR23782878 |
| Coreidae | Coreinae | Anisoscelini | *Anisoscelis* | *alipes* | Kieran et al. (2019) | SRR7819303 |
| Coreidae | Coreinae | Anisoscelini | *Anisoscelis* | *foliaceus* | | SRR23782867 |
| Coreidae | Coreinae | Anisoscelini | *Anisoscelis* | *gradadius* | Forthman et al. (2020) | SRR9224647 |
| Coreidae | Coreinae | Anisoscelini | *Chondrocera* | *laticornis* | Forthman et al. (2020) | SRR9224648 |
| Coreidae | Coreinae | Anisoscelini | *Diactor* | *bilineatus* | | SRR23782905 |
| Coreidae | Coreinae | Anisoscelini | *Holhymenia* | sp. | Forthman et al. (2020) | SRR9224649 |
| Coreidae | Coreinae | Anisoscelini | *Leptoglossus* | *clypealis* | Forthman et al. (2020) | SRR9224655 |
| Coreidae | Coreinae | Anisoscelini | *Leptoglossus* | *concolor* | Forthman et al. (2020) | SRR9224657 |
| Coreidae | Coreinae | Anisoscelini | *Leptoglossus* | *corculus* | Emberts et al. (2020) | SRR11213490 |
| Coreidae | Coreinae | Anisoscelini | *Leptoglossus* | *fulvicornis* | Emberts et al. (2020) | SRR11213497 |
| Coreidae | Coreinae | Anisoscelini | *Leptoglossus* | *gonagra* | Forthman et al. (2020) | SRR9224656 |
| Coreidae | Coreinae | Anisoscelini | *Leptoglossus* | *occidentalis* | | SRR23782873 |
| Coreidae | Coreinae | Anisoscelini | *Leptoglossus* | *oppositus* | Emberts et al. (2020) | SRR11213506 |
| Coreidae | Coreinae | Anisoscelini | *Leptoglossus* | *phyllopus* | Forthman et al. (2020) | SRR9224659 |
| Coreidae | Coreinae | Anisoscelini | *Leptoglossus* | *zonatus* | Emberts et al. (2020) | SRR11213485 |
| Coreidae | Coreinae | Anisoscelini | *Leptoscelis* | *matronalis* | Miller et al. (2022) | SRR23782872 |
| Coreidae | Coreinae | Anisoscelini | *Leptoscelis* | *pallida* | Miller et al. (2022) | SRR23782871 |
| Coreidae | Coreinae | Anisoscelini | *Leptoscelis* | *quadrisignatus* | Emberts et al. (2020) | SRR11213504 |
| Coreidae | Coreinae | Anisoscelini | *Leptoscelis* | *tricolor* | Forthman et al. (2020) | SRR9224658 |
| Coreidae | Coreinae | Anisoscelini | *Narnia* | *femorata* | Forthman et al. (2020) | SRR9224661 |
| Coreidae | Coreinae | Anisoscelini | *Narnia* | *snowi* | Forthman et al. (2020) | SRR9224660 |
| Coreidae | Coreinae | Anisoscelini | *Phthiacnemia* | *picta* | Forthman et al. (2020) | SRR9224663 |
| Coreidae | Coreinae | Anisoscelini | *Phthiadema* | *cyanea* | | SRR23782859 |
| Coreidae | Coreinae | Anisoscelini | *Ugnius* | *badius* | Miller et al. (2022) | SRR23782877 |
| Coreidae | Coreinae | Chariesterini | *Chariesterus* | *antennator* | Forthman et al. (2020) | SRR9224662 |
| Coreidae | Coreinae | Chariesterini | *Chariesterus* | *armatus* | Forthman et al. (2020) | SRR9224665 |
| Coreidae | Coreinae | Chariesterini | *Chariesterus* | *moestus* | Miller et al. (2022) | SRR23782908 |
| Coreidae | Coreinae | Chariesterini | *Plapigus* | *abdominalis* | Forthman et al. (2020) | SRR9224664 |
| Coreidae | Coreinae | Chelinideini | *Chelinidea* | *tabulata* | Forthman et al. (2020) | SRR9224678 |
| Coreidae | Coreinae | Chelinideini | *Chelinidea* | *vittiger* | Forthman et al. (2020) | SRR9224679 |
| Coreidae | Coreinae | Cloresmini | *Cloresmus* | *antennatus* | Forthman et al. (2020) | SRR9224632 |
| Coreidae | Coreinae | Cloresmini | *Notobitus* | *affinis* | Forthman et al. (2020) | SRR9224630 |
| Coreidae | Coreinae | Cloresmini | *Notobitus* | *meleagris* | Forthman et al. (2020) | SRR9224633 |
| Coreidae | Coreinae | Cloresmini | *Notobitus* | *sexguttatus* | Forthman et al. (2020) | SRR9224631 |

| | | | | | | |
|---|---|---|---|---|---|---|
| Coreidae | Coreinae | Colpurini | *Hygia* | sp. | Forthman et al. (2020) | SRR9224629 |
| Coreidae | Coreinae | Colpurini | *Hygia (Stenocolpura)* | sp. | Miller et al. (2022) | SRR23782888 |
| Coreidae | Coreinae | Colpurini | *Sciophyroides* | *sulcicrus* | | SRR23782899 |
| Coreidae | Coreinae | Colpurini | *Typhlocolpura* | *chinai* | Forthman et al. (2020) | SRR9224626 |
| Coreidae | Coreinae | Coreini | *Coreus* | *marginatus* | Forthman et al. (2020) | SRR9224619 |
| Coreidae | Coreinae | Daladerini | *Dalader* | *planiventris* | Forthman et al. (2020) | SRR9224618 |
| Coreidae | Coreinae | Daladerini | *Odontocurtus* | *consociatus* | Forthman et al. (2020) | SRR9224621 |
| Coreidae | Coreinae | Daladerini | *Odontorhopala* | *callosa* | Forthman et al. (2020) | SRR9224622 |
| Coreidae | Coreinae | Dasynini | *Aulacosternum* | *nigrorubrum* | | SRR23782896 |
| Coreidae | Coreinae | Dasynini | *Galaesus* | *hasticornis* | Forthman et al. (2020) | SRR9224694 |
| Coreidae | Coreinae | Discogastrini | *Cnemomis* | *nodulosa* | Miller et al. (2022) | SRR23782907 |
| Coreidae | Coreinae | Discogastrini | *Discogaster* | *dentipes* | Miller et al. (2022) | SRR23782904 |
| Coreidae | Coreinae | Discogastrini | *Savius* | *diagonalis* | Miller et al. (2022) | SRR23782902 |
| Coreidae | Coreinae | Discogastrini | *Savius* | *diversicornis* | Miller et al. (2022) | SRR23782901 |
| Coreidae | Coreinae | Discogastrini | *Savius* | *jurgiosus* | Forthman et al. (2020) | SRR9224712 |
| Coreidae | Coreinae | Discogastrini | *Scamurius* | *bachmanni* | Miller et al. (2022) | SRR23782900 |
| Coreidae | Coreinae | Gonocerini | *Cletoliturus* | *lituripennis* | Forthman et al. (2020) | SRR9224692 |
| Coreidae | Coreinae | Gonocerini | *Cletomorpha* | *benita* | Forthman et al. (2020) | SRR9224625 |
| Coreidae | Coreinae | Gonocerini | *Cletomorpha* | *nyasana* | Emberts et al. (2020) | SRR11213501 |
| Coreidae | Coreinae | Gonocerini | *Cletus* | *binotulatus* | Emberts et al. (2020) | SRR11213494 |
| Coreidae | Coreinae | Gonocerini | *Cletus* | *ochraceus* | Forthman et al. (2020) | SRR9224620 |
| Coreidae | Coreinae | Gonocerini | *Cletus* | sp. | Emberts et al. (2020) | SRR11213496 |
| Coreidae | Coreinae | Homoeocerini | *Fracastorius* | *cornutus* | Forthman et al. (2020) | SRR9224695 |
| Coreidae | Coreinae | Homoeocerini | *Homoeocerus* | *albiventris* | | SRR23782894 |
| Coreidae | Coreinae | Homoeocerini | *Homoeocerus* | *angulatus* | Emberts et al. (2020) | SRR11213499 |
| Coreidae | Coreinae | Homoeocerini | *Homoeocerus* | *bipustulatus* | Forthman et al. (2020) | SRR9224696 |
| Coreidae | Coreinae | Homoeocerini | *Homoeocerus* | *limbatipennis* | | SRR23782893 |
| Coreidae | Coreinae | Homoeocerini | *Homoeocerus* | *marginellus* | Emberts et al. (2020) | SRR11213514 |
| Coreidae | Coreinae | Homoeocerini | *Homoeocerus* | *simiolus* | | SRR23782891 |
| Coreidae | Coreinae | Homoeocerini | *Homoeocerus* | *subjectus* | | SRR23782890 |
| Coreidae | Coreinae | Homoeocerini | *Homoeocerus* | *walkeri* | | SRR23782889 |
| Coreidae | Coreinae | Homoeocerini | *Prismatocerus* | *auriculatus* | Emberts et al. (2020) | SRR11213502 |
| Coreidae | Coreinae | Hypselonotini | *Acidomeria* | *cinctipes* | Miller et al. (2022) | SRR23782920 |
| Coreidae | Coreinae | Hypselonotini | *Acidomeria* | *sordida* | Miller et al. (2022) | SRR23782911 |
| Coreidae | Coreinae | Hypselonotini | *Althos* | *obscurator* | Forthman et al. (2020) | SRR9224676 |
| Coreidae | Coreinae | Hypselonotini | *Anasa* | *andresii* | Emberts et al. (2020) | SRR11213511 |
| Coreidae | Coreinae | Hypselonotini | *Anasa* | *bellator* | Forthman et al. (2020) | SRR9224677 |
| Coreidae | Coreinae | Hypselonotini | *Anasa* | *scorbutica* | Emberts et al. (2020) | SRR11213510 |
| Coreidae | Coreinae | Hypselonotini | *Anasa* | *tristis* | Forthman et al. (2020) | SRR9224682 |
| Coreidae | Coreinae | Hypselonotini | *Anasa* | *varicornis* | Forthman et al. (2020) | SRR9224683 |
| Coreidae | Coreinae | Hypselonotini | *Catorhintha* | *apicalis* | | SRR23782876 |
| Coreidae | Coreinae | Hypselonotini | *Catorhintha* | *guttula* | Forthman et al. (2020) | SRR9224680 |
| Coreidae | Coreinae | Hypselonotini | *Catorhintha* | *selector* | | SRR23782875 |
| Coreidae | Coreinae | Hypselonotini | *Catorhintha* | *texana* | Forthman et al. (2020) | SRR9224681 |
| Coreidae | Coreinae | Hypselonotini | *Cebrenis* | *cauta* | Miller et al. (2022) | SRR23782874 |
| Coreidae | Coreinae | Hypselonotini | *Cebrenis* | *colorata* | Miller et al. (2022) | SRR23782910 |
| Coreidae | Coreinae | Hypselonotini | *Cebrenis* | *danieli* | Forthman et al. (2020) | SRR9224684 |
| Coreidae | Coreinae | Hypselonotini | *Cebrenistella* | *robusta* | Miller et al. (2022) | SRR23782909 |
| Coreidae | Coreinae | Hypselonotini | *Collatia* | *emarginata* | | SRR23782906 |
| Coreidae | Coreinae | Hypselonotini | *Hypselonotus* | *bitrianguliger* | Forthman et al. (2020) | SRR9224685 |
| Coreidae | Coreinae | Hypselonotini | *Hypselonotus* | *fulvus* | Forthman et al. (2020) | SRR9224675 |
| Coreidae | Coreinae | Hypselonotini | *Hypselonotus* | *lineatus* | Emberts et al. (2020) | SRR11213519 |
| Coreidae | Coreinae | Hypselonotini | *Hypselonotus* | *punctiventris* | Emberts et al. (2020) | SRR11213491 |
| Coreidae | Coreinae | Hypselonotini | *Namacus* | *annulicornis* | Miller et al. (2022) | SRR23782881 |
| Coreidae | Coreinae | Hypselonotini | *Paryphes* | *imperialis* | Miller et al. (2022) | SRR23782865 |
| Coreidae | Coreinae | Hypselonotini | *Paryphes* | *pulchellus* | Forthman et al. (2020) | SRR9224674 |
| Coreidae | Coreinae | Hypselonotini | *Petersitocoroides* | *longirostris* | Miller et al. (2022) | SRR23782861 |
| Coreidae | Coreinae | Hypselonotini | *Scolopocerus* | *secundarius* | Forthman et al. (2020) | SRR9224673 |
| Coreidae | Coreinae | Hypselonotini | *Sethenira* | *testacea* | Miller et al. (2022) | SRR23782898 |
| Coreidae | Coreinae | Hypselonotini | *Sphictyrtus* | *intermedius* | | SRR23782897 |
| Coreidae | Coreinae | Hypselonotini | *Vazquezitocoris* | *repletus* | Forthman et al. (2020) | SRR9224672 |
| Coreidae | Coreinae | Hypselonotini | *Villasitocoris* | *inconspicuus* | Forthman et al. (2020) | SRR9224671 |
| Coreidae | Coreinae | Hypselonotini | *Zicca* | *commaculata* | Forthman et al. (2020) | SRR9224670 |
| Coreidae | Coreinae | Hypselonotini | *Zicca* | *rubricator* | Forthman et al. (2020) | SRR9224669 |
| Coreidae | Coreinae | Hypselonotini | *Zicca* | *taeniola* | Forthman et al. (2020) | SRR9224668 |
| Coreidae | Coreinae | Latimbini | *Latimbus* | *concolor* | Forthman et al. (2020) | SRR9224617 |
| Coreidae | Coreinae | Latimbini | *Latimbus* | *refulgens* | Forthman et al. (2020) | SRR9224634 |
| Coreidae | Coreinae | Mictini | *Anoplocnemis* | *curvipes* | Emberts et al. (2020) | SRR11213513 |
| Coreidae | Coreinae | Mictini | *Anoplocnemis* | *phasiana* | Emberts et al. (2020) | SRR11213498 |
| Coreidae | Coreinae | Mictini | *Anoplocnemis* | sp.1 | Kieran et al. (2019) | SRR7819304 |
| Coreidae | Coreinae | Mictini | *Anoplocnemis* | sp.2 | Forthman et al. (2020) | SRR9224667 |
| Coreidae | Coreinae | Mictini | *Aspilosterna* | sp. | | SRR23782864 |
| Coreidae | Coreinae | Mictini | *Dianomictis* | *expansa* | Forthman et al. (2020) | SRR9224666 |
| Coreidae | Coreinae | Mictini | *Elasmopoda* | *alata* | Emberts et al. (2020) | SRR21508692 |
| Coreidae | Coreinae | Mictini | *Mictis* | *longicornis* | Emberts et al. (2020) | SRR11213516 |
| Coreidae | Coreinae | Mictini | *Mictis* | *profana* | Emberts et al. (2020) | SRR11213512 |
| Coreidae | Coreinae | Mictini | *Molipteryx* | *lunata* | Forthman et al. (2020) | SRR9224641 |
| Coreidae | Coreinae | Mictini | *Mygdonia* | *elongata* | Miller et al. (2022) | SRR23782882 |
| Coreidae | Coreinae | Mictini | *Mygdonia* | *tuberculosa* | Kieran et al. (2019) | SRR7819297 |
| Coreidae | Coreinae | Mictini | *Plectropoda* | *bicolor* | | SRR23782916 |
| Coreidae | Coreinae | Mictini | *Plectropoda* | *cruciata* | Forthman et al. (2020) | SRR9224699 |
| Coreidae | Coreinae | Mictini | *Plectropoda* | *oblongipes* | | SRR23782914 |
| Coreidae | Coreinae | Mictini | *Plectropoda* | sp. | Forthman et al. (2020) | SRR9224700 |
| Coreidae | Coreinae | Mictini | *Plectropodoides* | *dallastai* | Forthman et al. (2020) | SRR9224701 |
| Coreidae | Coreinae | Mictini | *Prionolomia* | *yunnanensis* | Forthman et al. (2020) | SRR9224640 |
| Coreidae | Coreinae | Mictini | *Pternistria* | *bispina* | Forthman et al. (2020) | SRR9224627 |
| Coreidae | Coreinae | Nematopodini | *Grammopoecilus* | *angustatus* | Forthman et al. (2020) | SRR9224706 |
| Coreidae | Coreinae | Nematopodini | *Melucha* | *chapadana* | Miller et al. (2022) | SRR23782869 |
| Coreidae | Coreinae | Nematopodini | *Melucha* | *dilatata* | | SRR23782868 |
| Coreidae | Coreinae | Nematopodini | *Melucha* | *quadrivittis* | Forthman et al. (2020) | SRR9224710 |

| | | | | | | |
|---|---|---|---|---|---|---|
| Coreidae | Coreinae | Nematopodini | *Melucha* | *quinquelineata* | | SRR23782866 |
| Coreidae | Coreinae | Nematopodini | *Mozena* | *arizonensis* | Miller et al. (2022) | SRR23782885 |
| Coreidae | Coreinae | Nematopodini | *Mozena* | *lineolata* | Kieran et al. (2019) | SRR7819301 |
| Coreidae | Coreinae | Nematopodini | *Mozena* | *lunata* | Miller et al. (2022) | SRR23782884 |
| Coreidae | Coreinae | Nematopodini | *Mozena* | *lurida* | Forthman et al. (2020) | SRR9224709 |
| Coreidae | Coreinae | Nematopodini | *Mozena* | *obtusa* | Miller et al. (2022) | SRR23782883 |
| Coreidae | Coreinae | Nematopodini | *Nematopus* | *lepidus* | Forthman et al. (2020) | SRR9224614 |
| Coreidae | Coreinae | Nematopodini | *Ouranion* | *serrulatus* | Forthman et al. (2020) | SRR9224707 |
| Coreidae | Coreinae | Nematopodini | *Pachylis* | *pharaonis* | | SRR23782880 |
| Coreidae | Coreinae | Nematopodini | *Piezogaster* | *calcarator* | Forthman et al. (2020) | SRR9224708 |
| Coreidae | Coreinae | Nematopodini | *Piezogaster* | *odiosus* | | SRR23782858 |
| Coreidae | Coreinae | Nematopodini | *Piezogaster* | *reclusus* | Miller et al. (2022) | SRR23782919 |
| Coreidae | Coreinae | Nematopodini | *Piezogaster* | *spurcus* | Miller et al. (2022) | SRR23782918 |
| Coreidae | Coreinae | Nematopodini | *Thasus* | *gigas* | | SRR23782879 |
| Coreidae | Coreinae | Nematopodini | *Thasus* | *neocalifornicus* | Kieran et al. (2019) | SRR7819295 |
| Coreidae | Coreinae | Petascelini | *Petascelis* | *remipes* | Forthman et al. (2020) | SRR9224615 |
| Coreidae | Coreinae | Petascelini | *Petillopsis* | *calcar* | | SRR23782860 |
| Coreidae | Coreinae | Phyllomorphini | *Pephricus* | *paradoxus* | Emberts et al. (2020) | SRR11213503 |
| Coreidae | Coreinae | Phyllomorphini | *Tongorma* | *latreillii* | Forthman et al. (2020) | SRR9224623 |
| Coreidae | Coreinae | Placoscelini | *Plaxiscelis* | *limbata* | Forthman et al. (2020) | SRR9224702 |
| Coreidae | Coreinae | Placoscelini | *Plaxiscelis* | *pagana* | Miller et al. (2022) | SRR23782917 |
| Coreidae | Coreinae | Placoscelini | *Stenoeurilla* | *mesoamericana* | Forthman et al. (2020) | SRR9224703 |
| Coreidae | Coreinae | Spartocerini | *Menenotus* | *diminutus* | | SRR23782886 |
| Coreidae | Coreinae | Spartocerini | *Sephina* | *geniculata* | Forthman et al. (2020) | SRR9224686 |
| Coreidae | Coreinae | Spartocerini | *Sephina* | *subulata* | Forthman et al. (2020) | SRR9224687 |
| Coreidae | Coreinae | Spartocerini | *Spartocera* | *batatas* | Emberts et al. (2020) | SRR11213493 |
| Coreidae | Coreinae | Spartocerini | *Spartocera* | *fusca* | Forthman et al. (2020) | SRR9224616 |
| Coreidae | Hydarinae | | *Hydara* | *tenuicornis* | Forthman et al. (2019) | SRR8903500 |
| Coreidae | Hydarinae | | *Hydarella* | *chiangdaoensis* | Forthman et al. (2022) | SRR16596594 |
| Coreidae | Hydarinae | | *Hydaropsis* | *longirostris* | Forthman et al. (2022) | SRR16596593 |
| Coreidae | Hydarinae | | *Madura* | *fuscoclavata* | Forthman et al. (2022) | SRR16596588 |
| Coreidae | Hydarinae | | *Maduranoides* | *chemsaki* | Forthman et al. (2022) | SRR16596587 |
| Coreidae | Meropachyinae | Merocorini | *Merocoris* | *curtatus* | Forthman et al. (2020) | SRR9224704 |
| Coreidae | Meropachyinae | Merocorini | *Merocoris* | *elevatus* | Forthman et al. (2020) | SRR9224705 |
| Coreidae | Meropachyinae | Merocorini | *Merocoris* | *typhaeus* | Forthman et al. (2020) | SRR9224697 |
| Coreidae | Meropachyinae | Meropachyini | *Alcocerniella* | *limonesis* | Miller et al. (2022) | SRR23782892 |
| Coreidae | Meropachyinae | Meropachyini | *Salamancaniella* | *alternata* | | SRR23782903 |
| Coreidae | Meropachyinae | Spathophorini | *Himellastella* | *conica* | | SRR23782895 |
| Coreidae | Meropachyinae | Spathophorini | *Lycambes* | *sargi* | Kieran et al. (2019) | SRR7819300 |
| Coreidae | Pseudophloeinae | Clavigrallini | *Clavigralla* | *minor* | Forthman et al. (2022) | SRR16596612 |
| Coreidae | Pseudophloeinae | Clavigrallini | *Clavigralla* | *pusilla* | Forthman et al. (2022) | SRR16596610 |
| Coreidae | Pseudophloeinae | Clavigrallini | *Clavigralla* | *shadabi* | Forthman et al. (2022) | SRR16596609 |
| Coreidae | Pseudophloeinae | Clavigrallini | *Clavigralloides* | *acantharis* | Forthman et al. (2022) | SRR16596608 |
| Coreidae | Pseudophloeinae | Clavigrallini | *Gralliclava* | *horrens* | Forthman et al. (2022) | SRR16596601 |
| Coreidae | Pseudophloeinae | Pseudophloeini | *Arenocoris* | *waltlii* | Forthman et al. (2022) | SRR16596591 |
| Coreidae | Pseudophloeinae | Pseudophloeini | *Bathysolen* | *nubilus* | Forthman et al. (2022) | SRR16596580 |
| Coreidae | Pseudophloeinae | Pseudophloeini | *Ceraleptus* | *gracilicornis* | Forthman et al. (2022) | SRR16596565 |
| Coreidae | Pseudophloeinae | Pseudophloeini | *Ceraleptus* | *obtusus* | Forthman et al. (2022) | SRR16596564 |
| Coreidae | Pseudophloeinae | Pseudophloeini | *Coriomeris* | *affinis* | Forthman et al. (2022) | SRR16596607 |
| Coreidae | Pseudophloeinae | Pseudophloeini | *Coriomeris* | *denticulatus* | Forthman et al. (2022) | SRR16596606 |
| Coreidae | Pseudophloeinae | Pseudophloeini | *Coriomeris* | *nigricornis* | Forthman et al. (2022) | SRR16596605 |
| Coreidae | Pseudophloeinae | Pseudophloeini | *Hoplolomia* | *scabricula* | Forthman et al. (2022) | SRR16596598 |
| Coreidae | Pseudophloeinae | Pseudophloeini | *Mevanidea* | *spiniceps* | Forthman et al. (2022) | SRR16596584 |
| Coreidae | Pseudophloeinae | Pseudophloeini | *Myla* | sp. | Forthman et al. (2019) | SRR8903488 |
| Coreidae | Pseudophloeinae | Pseudophloeini | *Paramyla* | *suspecta* | Forthman et al. (2022) | SRR16596578 |
| Coreidae | Pseudophloeinae | Pseudophloeini | *Pseudomyla* | *cornuta* | Forthman et al. (2022) | SRR16596577 |
| Coreidae | Pseudophloeinae | Pseudophloeini | *Psilolomia* | *brunneofusca* | Forthman et al. (2022) | SRR16596576 |
| Coreidae | Pseudophloeinae | Pseudophloeini | *Psilolomia* | *parva* | Forthman et al. (2022) | SRR16596575 |
| Coreidae | Pseudophloeinae | Pseudophloeini | *Strobilotoma* | *typhaecornis* | Forthman et al. (2022) | SRR16596572 |
| Coreidae | Pseudophloeinae | Pseudophloeini | *Vilga* | *dallasi* | Forthman et al. (2022) | SRR16596570 |
| Coreidae | Pseudophloeinae | Pseudophloeini | *Vilga* | *sanctipauli* | Forthman et al. (2022) | SRR16596568 |
| Coreidae | Pseudophloeinae | Pseudophloeini | *Vilga* | *westwoodi* | Forthman et al. (2022) | SRR16596567 |
| Largidae | Larginae | Largini | *Largus* | sp. | Forthman et al. (2019) | SRR8903493 |
| Lygaeidae | Lygaeinae | Lygaeini | *Oncopeltus* | *fasciatus* | | GCA_000696205.2 |
| Lygaeidae | Lygaeinae | Lygaeini | *Oncopeltus* | *cingulifer* | Emberts et al. (2020) | SRR11213515 |
| Pentatomidae | Pentatominae | Cappaeini | *Halyomorpha* | *halys* | | GCA_000696795.3 |
| Pyrrhocoridae | | | *Dysdercus* | *suturellus* | Forthman et al. (2019) | SRR8903495 |
| Rhopalidae | Rhopalinae | Chorosomatini | *Chorosoma* | *schillingii* | Forthman et al. (2022) | SRR16596563 |
| Rhopalidae | Rhopalinae | Harmostini | *Harmostes* | *serratus* | Forthman et al. (2019) | SRR8903499 |
| Rhopalidae | Rhopalinae | Rhopalini | *Corizus* | *hyoscyami* | Forthman et al. (2022) | SRR16596604 |
| Rhopalidae | Rhopalinae | Rhopalini | *Liorhyssus* | *hyalinus* | Forthman et al. (2022) | SRR16596590 |
| Rhopalidae | Rhopalinae | Rhopalini | *Maccevethus* | *errans* | Forthman et al. (2022) | SRR16596589 |
| Rhopalidae | Serinethinae | | *Jadera* | *haematoloma* | Forthman et al. (2019) | SRR8903498 |

Table S2. Sequence data summary. Abbreviations: bp, base pairs; EtOH, ethanol; Max., maximum; Min., minimum; UCE, ultraconserved element.

| **Species** | **Preservation** | **Total raw reads** | **Total filtered reads** | **Total contigs** | **Total bp** | **Mean contig length** | **Min. contig length** | **Max. contig length** | **Total UCEs** | **% UCEs recovered** | **Mean UCE length** | **Min. UCE length** | **Max. UCE length** |
|---|---|---|---|---|---|---|---|---|---|---|---|---|---|
| *Anisoscelis foliaceus* | Dried | 535232 | 250845 | 3917 | 325962 | 83.22 | 56 | 712 | 24 | 0.90 | 299.21 | 208 | 642 |
| *Aspilosterna* sp. | Dried | 7600142 | 2112140 | 13780 | 1977299 | 143.49 | 56 | 2625 | 635 | 23.76 | 313.26 | 206 | 2625 |
| *Athaumastus lugens* | Dried | 1833266 | 745325 | 12858 | 1207796 | 93.93 | 56 | 1468 | 178 | 6.66 | 297.36 | 206 | 1426 |
| *Aulacosternum nigrorubrum* | Dried | 2031522 | 1120685 | 22799 | 2469311 | 108.31 | 56 | 6250 | 522 | 19.53 | 318.99 | 206 | 2042 |
| *Catorhintha apicalis* | EtOH | 8629218 | 5042288 | 21803 | 8458100 | 387.93 | 56 | 4073 | 1419 | 53.09 | 820.51 | 206 | 3684 |
| *Catorhintha selector* | EtOH | 5983458 | 3389011 | 16115 | 6818647 | 423.12 | 56 | 3718 | 1392 | 52.08 | 834.28 | 207 | 3107 |
| *Collatia emarginata* | Dried | 654436 | 246703 | 4765 | 432322 | 90.73 | 56 | 1015 | 39 | 1.46 | 269.92 | 207 | 648 |
| *Diactor bilineatus* | Dried | 5762776 | 2138737 | 21454 | 2426674 | 113.11 | 56 | 2122 | 795 | 29.74 | 314.47 | 206 | 2122 |
| *Himellastella concia* | Dried | 3021462 | 1274663 | 8821 | 2175568 | 246.64 | 56 | 2206 | 1111 | 41.56 | 356.85 | 206 | 2206 |
| *Homoeocerus albiventris* | EtOH | 1909402 | 1098949 | 16840 | 5145291 | 305.54 | 56 | 6787 | 1114 | 41.68 | 586.75 | 206 | 2598 |
| *Homoeocerus limbatipennis* | EtOH | 4858488 | 2732985 | 18076 | 6751613 | 373.51 | 56 | 6307 | 1444 | 54.02 | 750.02 | 206 | 4553 |
| *Homoeocerus simiolus* | EtOH | 724614 | 354944 | 6242 | 1676213 | 268.54 | 56 | 2949 | 744 | 27.83 | 319.58 | 206 | 2177 |
| *Homoeocerus subjectus* | EtOH | 3089966 | 1712247 | 13086 | 5074318 | 387.77 | 56 | 3432 | 1350 | 50.51 | 701.51 | 208 | 3015 |
| *Homoeocerus walkeri* | EtOH | 3973580 | 2190185 | 23573 | 5180062 | 219.75 | 56 | 2655 | 1485 | 55.56 | 451.89 | 206 | 2655 |
| *Leptoglossus occidentalis* | Dried | 3347072 | 1580042 | 20861 | 2960846 | 141.93 | 56 | 4120 | 868 | 32.47 | 352.15 | 206 | 2332 |
| *Melucha dilatata* | EtOH | 2207132 | 1284810 | 13678 | 5509845 | 402.83 | 56 | 2992 | 1269 | 47.47 | 712.68 | 206 | 2962 |
| *Melucha quinquelineata* | Dried | 8557872 | 3604416 | 22006 | 7729381 | 351.24 | 56 | 5071 | 1388 | 51.93 | 672.23 | 195 | 3452 |
| *Menenotus diminutus* | EtOH | 15095080 | 8762469 | 62506 | 22188283 | 354.98 | 56 | 16425 | 1527 | 57.13 | 1104.63 | 207 | 5985 |
| *Pachylis pharaonis* | Dried | 7223490 | 1933083 | 10183 | 1959646 | 192.44 | 56 | 2400 | 1019 | 38.12 | 340.52 | 182 | 2170 |
| *Petillopsis calcar* | Dried | 3110750 | 1391027 | 13109 | 2467684 | 188.24 | 56 | 3456 | 1009 | 37.75 | 351.18 | 206 | 2165 |
| *Phthiadema cyanea* | Dried | 530972 | 239257 | 4620 | 380651 | 82.39 | 56 | 779 | 13 | 0.49 | 293.15 | 210 | 517 |
| *Piezogaster odiosus* | EtOH | 6083788 | 3991886 | 38793 | 12413312 | 319.99 | 56 | 7369 | 1441 | 53.91 | 862.82 | 207 | 3281 |
| *Plectropoda bicolor* | Dried | 3141520 | 766628 | 4207 | 750937 | 178.50 | 56 | 2061 | 427 | 15.97 | 315.10 | 165 | 2061 |
| *Plectropoda oblongipes* | Dried | 2300414 | 826443 | 12082 | 1327763 | 109.90 | 56 | 1863 | 295 | 11.04 | 320.27 | 148 | 1863 |
| *Pomponatius typicus* | Dried | 5480332 | 2166713 | 14072 | 2927243 | 208.02 | 56 | 3634 | 1264 | 47.29 | 425.96 | 92 | 2677 |
| *Salamancaniella alternata* | EtOH | 5251544 | 3632223 | 49967 | 10506520 | 210.27 | 56 | 5103 | 1350 | 50.51 | 749.58 | 206 | 3756 |
| *Sciophyroides sulcicrus* | Dried | 2302026 | 976827 | 17157 | 1452044 | 84.63 | 56 | 1931 | 185 | 6.92 | 329.84 | 206 | 1582 |
| *Sphictyrtus intermedius* | Dried | 1847518 | 578118 | 8967 | 892803 | 99.57 | 56 | 2636 | 969 | 36.25 | 352.47 | 193 | 2202 |
| *Thasus gigas* | Dried | 7449708 | 3113434 | 41566 | 4296876 | 103.37 | 56 | 2202 | 1122 | 41.98 | 650.91 | 208 | 4349 |
| *Turrana abnormis* | EtOH | 1691596 | 1098330 | 20161 | 4858306 | 240.98 | 56 | 4349 | 1444 | 0.90 | 299.21 | 206 | 4553 |

Table S3. Summary of informative sites and number of UCE loci in datasets generated in this study.

| Dataset | Total UCEs | Total partitions (concatenated alignments) | Total sites | % Parsimony-informative sites | % Uninformative sites | % Invariant sites |
|---|---|---|---|---|---|---|
| 50p | 1093 | 147 | 284013 | 46.61 | 7.78 | 45.61 |
| 50p25mi | 273 | NA | 136097 | 54.77 | 7.78 | 37.45 |
| 70p | 647 | 114 | 180597 | 47.21 | 7.93 | 44.86 |
| 70p25mi | 162 | NA | 90659 | 55.25 | 7.93 | 36.82 |

Table S4. Summary of component state gains and losses across different ultrametric trees (component state 0 not reported). Abbreviations: ARD, all rates different model; ER, equal rates model; G–L, gains minus losses; Min, minimum; ML, maximum likelihood; MSC, multispecies coalescent; N, sample size; SYM, symmetric model; 50p, alignments comprised of locus alignments with at least 50% of the total taxa sampled in this study; 50p25mi, 25% most informative loci subsampled from the 50p dataset.

| **Component number** | **Model** | **Component state** | **N taxa** | **50p ML** | | **50p MSC** | | **50p25mi MSC** | | **Min gains** | **Min losses** | **G–L** |
|---|---|---|---|---|---|---|---|---|---|---|---|---|
| | | | | **Gains** | **Losses** | **Gains** | **Losses** | **Gains** | **Losses** | | | |
| **1** | ER | 1 | 5 | 2 | 0 | 2 | 0 | 2 | 0 | **2** | **0** | **2** |
| **2** | ARD | 1 | 13 | 8 | 2 | 8 | 2 | 8 | 2 | **8** | **2** | **6** |
| **3** | ER | 1 | 89 | 19 | 3 | 20 | 3 | 21 | 3 | **19** | **3** | **16** |
| **4** | ARD | 1 | 18 | 6 | 4–5 | 6 | 5 | 6 | 5 | **6** | **4** | **2** |
| **5** | ER | 1 | 9 | 7 | 0 | 7 | 0 | 7 | 0 | **7** | **0** | **7** |
| **6** | SYM | 1 | 36 | 18 | 4 | 21 | 2–3 | 21 | 2 | **18** | **2** | **16** |
| | | 2 | 31 | 16 | 11 | 14 | 15 | 13 | 13 | **13** | **11** | **2** |
| | | 3 | 103 | 21 | 26 | 13–14 | 32 | 13 | 34 | **13** | **26** | **-13** |
| **7** | ARD | 1 | 59 | 15 | 17 | 16 | 12–13 | 16 | 14 | **15** | **12** | **3** |
| **8** | ARD | 1 | 55 | 14 | 28–30 | 17 | 18 | 16 | 19–20 | **14** | **18** | **-4** |
| **9** | ARD | 1 | 27 | 5 | 9 | 5 | 8 | 6 | 7 | **5** | **7** | **-2** |
| **10** | ER | 1 | 16 | 4 | 2 | 4 | 2 | 4 | 2 | **4** | **2** | **2** |
| | | 2 | 18 | 9 | 1 | 9 | 1 | 9 | 1 | **9** | **1** | **8** |
| | | 3 | 2 | 2 | 0 | 2 | 0 | 2 | 0 | **2** | **0** | **2** |
| **11** | ARD | 1 | 53 | 5 | 15 | 6 | 11 | 6 | 12 | **5** | **11** | **-6** |
| **12** | ARD | 1 | 47 | 6 | 9 | 5 | 13 | 5 | 13 | **5** | **9** | **-4** |
| **13** | ER | 1 | 111 | 15 | 7 | 15 | 7 | 15 | 7 | **15** | **7** | **8** |
| **14** | ARD | 1 | 45 | 13 | 7 | 13 | 7 | 14 | 7 | **13** | **7** | **6** |
| **15** | ARD | 1 | 26 | 5 | 3 | 5 | 3 | 5 | 3 | **5** | **3** | **2** |

Table S5. Results of Pagel's (1994) binary correlated traits test for each pairwise comparison of components found to have significant co-occurrence (from co-occur results) based on the 50p maximum likelihood (ML), 50p multispecies coalescent (MSC), and 50p25mi MSC ultrametric trees. Significant p-values are bolded. Abbreviations: AIC(D), Akaike Information Criterion for dependent model; AIC(I), Akaike Information Criterion for independent model; ARD, all rates different model; BH, Benjamini-Hochberg correction for multiple comparisons (Benjamini & Hochberg, 1995); ER, equal rates model; LR, likelihood ratio; NT, not tested (due to potential unreplicated effects based on ancestral state estimation of components; see main text for more details).

| Component | | | 50p ML | | | | | | | 50p MSC | | | | | | | 50p25mi MSC | | | | | | |
|---|---|---|---|---|---|---|---|---|---|---|---|---|---|---|---|---|---|---|---|---|---|---|---|
| **Comparison** | **Model** | **Dependent** | **L(D)** | **L(I)** | **AIC(D)** | **AIC(I)** | **LR** | **p-value** | **BH p-value** | **L(D)** | **L(I)** | **AIC(D)** | **AIC(I)** | **LR** | **p-value** | **BH p-value** | **L(D)** | **L(I)** | **AIC(D)** | **AIC(I)** | **LR** | **p-value** | **BH p-value** |
| 1 vs. 3 | ER | Component 1 | -91.6230 | -93.1845 | 189.2459 | 190.3691 | 3.1232 | 0.0772 | 0.1647 | -93.1037 | -94.4150 | 192.2074 | 192.8299 | 2.6226 | 0.1054 | 0.1976 | -94.33206 | -95.6298 | 194.6641 | 195.2596 | 2.5955 | 0.1072 | 0.1998 |
| | ER | Component 3 | -92.3466 | -93.1845 | 190.6931 | 190.3691 | 1.6760 | 0.1955 | 0.3293 | -93.1167 | -94.4150 | 192.2334 | 192.8299 | 2.5966 | 0.1071 | 0.1996 | -94.4378 | -95.6298 | 194.8756 | 195.2596 | 2.3839 | 0.1226 | 0.2232 |
| | ER | Interdependent | -91.2609 | -93.1845 | 190.5219 | 190.3691 | 3.8472 | 0.1461 | 0.2645 | -92.7290 | -94.4150 | 193.4579 | 192.8299 | 3.3720 | 0.1853 | 0.2963 | -93.9235 | -95.6298 | 195.8469 | 195.2596 | 3.4126 | 0.1815 | 0.2873 |
| 1 vs. 13 | ER | Component 1 | -85.3718 | -86.5891 | 176.7435 | 177.1782 | 2.4347 | 0.1187 | 0.2294 | -88.2292 | -89.4757 | 182.4583 | 182.9515 | 2.4931 | 0.1143 | 0.2057 | -87.8658 | -89.0589 | 181.7316 | 182.1178 | 2.3862 | 0.1224 | 0.2232 |
| | ER | Component 13 | -86.5254 | -86.5891 | 179.0509 | 177.1782 | 0.1274 | 0.7212 | 0.8103 | -89.3857 | -89.4757 | 184.7713 | 182.9515 | 0.1801 | 0.6713 | 0.7861 | -88.9680 | -89.0589 | 183.9360 | 182.1178 | 0.1818 | 0.6698 | 0.7814 |
| | ER | Interdependent | -84.9975 | -86.5891 | 177.9949 | 177.1782 | 3.1833 | 0.2036 | 0.3393 | -87.8302 | -89.4757 | 183.6604 | 182.9515 | 3.2911 | 0.1929 | 0.3053 | -87.4548 | -89.0589 | 182.9095 | 182.1178 | 3.2083 | 0.2011 | 0.3152 |
| 2 vs. 6 | ARD | Component 2 | -108.2978 | -111.5385 | 228.5956 | 231.0771 | 6.4815 | **0.0391** | 0.0988 | NT | NT | NT | NT | NT | NT | NT | NT | NT | NT | NT | NT | NT | NT |
| | ARD | Component 6 | -109.5643 | -111.5385 | 231.1285 | 231.0771 | 3.9485 | 0.1389 | 0.2589 | NT | NT | NT | NT | NT | NT | NT | NT | NT | NT | NT | NT | NT | NT |
| | ARD | Interdependent | -108.0952 | -111.5385 | 232.1904 | 231.0771 | 6.8866 | 0.1420 | 0.2601 | NT | NT | NT | NT | NT | NT | NT | NT | NT | NT | NT | NT | NT | NT |
| 2 vs. 7 | ARD | Component 2 | -129.6391 | -137.0200 | 271.2781 | 282.0401 | 14.7619 | **0.0006** | **0.0042** | -130.8203 | -137.6796 | 273.6406 | 283.3591 | 13.7185 | **0.0010** | **0.0053** | -131.9501 | -139.0492 | 275.9002 | 286.0984 | 14.1982 | **0.0008** | **0.0048** |
| | ARD | Component 7 | -128.5627 | -137.0200 | 269.1254 | 282.0401 | 16.9147 | **0.0002** | **0.0018** | -130.1221 | -137.6796 | 272.2441 | 283.3591 | 15.1150 | **0.0005** | **0.0031** | -130.8777 | -139.0492 | 273.7554 | 286.0984 | 16.3431 | **0.0003** | **0.0021** |
| | ARD | Interdependent | -126.7169 | -137.0200 | 269.4339 | 282.0401 | 20.6062 | **0.0004** | **0.0032** | -128.2391 | -137.6796 | 272.4782 | 283.3591 | 18.8809 | **0.0008** | **0.0043** | -128.6060 | -139.0492 | 273.2120 | 286.0984 | 20.8864 | **0.0003** | **0.0021** |
| 2 vs. 8 | ARD | Component 2 | -130.2383 | -135.6892 | 272.4765 | 279.3783 | 10.9018 | **0.0043** | **0.0217** | -131.6868 | -137.1896 | 275.3736 | 282.3792 | 11.0056 | **0.0041** | **0.0166** | -128.6578 | -134.8401 | 269.3155 | 277.6802 | 12.3646 | **0.0021** | **0.0105** |
| | ARD | Component 8 | -130.2703 | -135.6892 | 272.5406 | 279.3783 | 10.8378 | **0.0044** | **0.0217** | -132.8596 | -137.1896 | 277.7193 | 282.3792 | 8.6599 | **0.0132** | **0.0415** | -130.1629 | -134.8401 | 272.3259 | 277.6802 | 9.3543 | **0.0093** | **0.0332** |
| | ARD | Interdependent | -129.0000 | -135.6892 | 274.2573 | 279.3783 | 13.1211 | **0.0107** | **0.0396** | -131.6619 | -137.1896 | 279.3238 | 282.3792 | 11.0553 | **0.0259** | 0.0703 | -128.8115 | -134.8401 | 273.6230 | 277.6802 | 12.0572 | **0.0169** | 0.0507 |
| 2 vs. 9 | ARD | Component 2 | -76.0012 | -86.2761 | 164.0024 | 180.5522 | 20.5498 | **0.0000** | **0.0000** | -77.5627 | -88.4221 | 167.1253 | 184.8442 | 21.7189 | **0.0000** | **0.0000** | -76.6878 | -87.8197 | 165.3756 | 183.6395 | 22.2638 | **0.0000** | **0.0000** |
| | ARD | Component 9 | -72.7236 | -86.2761 | 157.4473 | 180.5522 | 27.1050 | **0.0000** | **0.0000** | -73.9968 | -88.4221 | 159.9935 | 184.8442 | 28.8507 | **0.0000** | **0.0000** | -73.5395 | -87.8197 | 159.0791 | 183.6395 | 28.5604 | **0.0000** | **0.0000** |
| | ARD | Interdependent | -72.1799 | -86.2761 | 160.3597 | 180.5522 | 28.1925 | **0.0000** | **0.0000** | -73.3571 | -88.4221 | 162.7141 | 184.8442 | 30.1301 | **0.0000** | **0.0000** | -72.7358 | -87.8197 | 161.4715 | 183.6395 | 30.1679 | **0.0000** | **0.0000** |
| 2 vs. 13 | ARD | Component 2 | -112.0250 | -114.4532 | 236.0499 | 236.9064 | 4.8565 | 0.0882 | 0.1785 | -115.7058 | -118.3888 | 243.4117 | 244.7776 | 5.3659 | 0.0684 | 0.1446 | -115.2690 | -118.1084 | 242.5381 | 244.2168 | 5.6788 | 0.0585 | 0.1316 |
| | ARD | Component 13 | -109.1833 | -114.4532 | 230.3665 | 236.9064 | 10.5398 | **0.0051** | **0.0233** | -111.7488 | -118.3888 | 235.4975 | 244.7776 | 13.2801 | **0.0013** | **0.0066** | -111.4588 | -118.1084 | 234.9176 | 244.2168 | 13.2992 | **0.0013** | **0.0069** |
| | ARD | Interdependent | -111.8110 | -114.4532 | 239.6221 | 236.9064 | 5.2843 | 0.2594 | 0.4005 | -111.6201 | -118.3888 | 239.2402 | 244.7776 | 13.5374 | **0.0089** | **0.0301** | -111.3937 | -118.1084 | 238.7873 | 244.2168 | 13.4295 | **0.0094** | **0.0332** |
| 3 vs. 4 | ARD | Component 3 | -117.2166 | -119.6422 | 246.4332 | 247.2845 | 4.8513 | 0.0884 | 0.1785 | -119.0646 | -122.3738 | 250.1292 | 252.7477 | 6.6185 | **0.0365** | 0.0923 | -119.6551 | -122.8300 | 251.3103 | 253.6601 | 6.3498 | **0.0418** | 0.1005 |
| | ARD | Component 4 | -115.3736 | -119.6422 | 242.7472 | 247.2845 | 8.5373 | **0.0140** | **0.0478** | -118.0520 | -122.3738 | 248.1039 | 252.7477 | 8.6438 | **0.0133** | **0.0415** | -118.6367 | -122.8300 | 249.2735 | 253.6601 | 8.3866 | **0.0151** | **0.0479** |
| | ARD | Interdependent | -114.4982 | -119.6422 | 244.9964 | 247.2845 | 10.2881 | **0.0358** | 0.0948 | -117.1943 | -122.3738 | 250.3886 | 252.7477 | 10.3591 | **0.0348** | 0.0906 | -117.7462 | -122.8300 | 251.4924 | 253.6601 | 10.1677 | **0.0377** | 0.0928 |
| 3 vs. 6 | ER | Component 3 | -148.4390 | -152.5317 | 302.8780 | 309.0635 | 8.1855 | **0.0042** | **0.0217** | NT | NT | NT | NT | NT | NT | NT | NT | NT | NT | NT | NT | NT | NT |
| | ER | Component 6 | -151.2026 | -152.5317 | 308.4052 | 309.0635 | 2.6583 | 0.1030 | 0.2003 | NT | NT | NT | NT | NT | NT | NT | NT | NT | NT | NT | NT | NT | NT |
| | ER | Interdependent | -146.7309 | -152.5317 | 301.4617 | 309.0635 | 11.6018 | **0.0030** | **0.0160** | NT | NT | NT | NT | NT | NT | NT | NT | NT | NT | NT | NT | NT | NT |
| 3 vs. 7 | ARD | Component 3 | -176.6846 | -177.3328 | 365.3691 | 362.6655 | 1.2964 | 0.5230 | 0.6590 | -178.3758 | -178.3983 | 368.7571 | 364.7966 | 0.0395 | 0.9805 | 1.0000 | -180.2770 | -180.8233 | 372.5540 | 369.6465 | 1.0925 | 0.5791 | 0.7070 |
| | ARD | Component 7 | -173.0705 | -177.3328 | 358.1409 | 362.6655 | 8.5246 | **0.0141** | **0.0478** | -173.5405 | -178.3983 | 359.0810 | 364.7966 | 9.7156 | **0.0078** | **0.0270** | -176.1646 | -180.8233 | 364.3292 | 369.6465 | 9.3173 | **0.0095** | **0.0332** |
| | ARD | Interdependent | -172.6328 | -177.3328 | 361.2655 | 362.6655 | 9.4000 | 0.0518 | 0.1267 | -172.8977 | -178.3983 | 361.7954 | 364.7966 | 11.0012 | **0.0266** | 0.0716 | -175.6755 | -180.8233 | 367.3509 | 369.6465 | 10.2956 | **0.0357** | 0.0893 |
| 3 vs. 8 | ARD | Component 3 | -173.3181 | -176.0019 | 358.6363 | 360.0038 | 5.3676 | 0.0683 | 0.1524 | -175.6397 | -177.9083 | 363.2794 | 363.8166 | 4.5372 | 0.1035 | 0.1976 | -174.0630 | -176.6141 | 360.1261 | 361.2283 | 5.1022 | 0.0780 | 0.1595 |
| | ARD | Component 8 | -172.5478 | -176.0019 | 357.0956 | 360.0038 | 6.9082 | **0.0316** | 0.0858 | -174.6045 | -177.9083 | 361.2091 | 363.8166 | 6.6076 | **0.0367** | 0.0923 | -174.2681 | -176.6141 | 360.5361 | 361.2283 | 4.6922 | 0.0957 | 0.1896 |
| | ARD | Interdependent | -171.3979 | -176.0019 | 358.7958 | 360.0038 | 9.2080 | 0.0561 | 0.1329 | -173.2813 | -177.9083 | 362.5624 | 363.8166 | 9.2541 | 0.0551 | 0.1258 | -173.0399 | -176.6141 | 362.0797 | 361.2283 | 7.1486 | 0.1282 | 0.2294 |
| 3 vs. 9 | ARD | Component 3 | -122.4568 | -126.5888 | 256.9135 | 261.1777 | 8.2642 | **0.0160** | 0.0520 | -124.2236 | -129.1408 | 260.4471 | 266.2817 | 9.8345 | **0.0073** | **0.0261** | -125.2101 | -129.5938 | 262.4202 | 267.1876 | 8.7673 | **0.0125** | **0.0414** |
| | ARD | Component 9 | -121.1713 | -126.5888 | 254.3426 | 261.1777 | 10.8351 | **0.0044** | **0.0217** | -122.0574 | -129.1408 | 256.1147 | 266.2817 | 14.1669 | **0.0008** | **0.0043** | -122.3199 | -129.5938 | 256.6398 | 267.1876 | 14.5478 | **0.0007** | **0.0042** |
| | ARD | Interdependent | -119.9852 | -126.5888 | 255.9649 | 261.1777 | 13.2128 | **0.0103** | **0.0386** | -118.4747 | -129.1408 | 252.9494 | 266.2817 | 21.3323 | **0.0003** | **0.0021** | -119.1129 | -129.5938 | 254.2257 | 267.1876 | 20.9618 | **0.0003** | **0.0021** |
| 3 vs. 10 | ER | Component 3 | -144.9175 | -145.4763 | 295.8350 | 294.9527 | 1.1177 | 0.2904 | 0.4295 | -146.0764 | -146.5368 | 298.1527 | 297.0735 | 0.9208 | 0.3373 | 0.4722 | -146.7230 | -147.0873 | 299.4460 | 298.1746 | 0.7286 | 0.3933 | 0.5250 |
| | ER | Component 10 | -141.6375 | -145.4763 | 289.2750 | 294.9527 | 7.6777 | **0.0056** | **0.0252** | -142.5897 | -146.5368 | 291.1794 | 297.0735 | 7.8942 | **0.0050** | **0.0197** | -143.1464 | -147.0873 | 292.2927 | 298.1746 | 7.8818 | **0.0050** | **0.0207** |
| | ER | Interdependent | -141.5430 | -145.4763 | 291.0860 | 294.9527 | 7.8667 | **0.0196** | 0.0605 | -142.5348 | -146.5368 | 293.0696 | 297.0735 | 8.0040 | **0.0183** | 0.0519 | -143.1142 | -147.0873 | 294.2283 | 298.1746 | 7.9463 | **0.0188** | 0.0553 |
| 3 vs. 11 | ARD | Component 3 | -136.2416 | -141.0934 | 284.4832 | 290.1869 | 9.7036 | **0.0078** | **0.0323** | -137.5696 | -145.4851 | 287.1392 | 298.9702 | 15.8311 | **0.0004** | **0.0026** | -139.6938 | -147.4055 | 291.3876 | 302.8109 | 15.4233 | **0.0004** | **0.0027** |
| | ARD | Component 11 | -140.4234 | -141.0934 | 292.8467 | 290.1869 | 1.3401 | 0.5117 | 0.6499 | -143.1619 | -145.4851 | 298.3238 | 298.9702 | 4.6464 | 0.0980 | 0.1900 | -145.1380 | -147.4055 | 302.2761 | 302.8109 | 4.5349 | 0.1036 | 0.1961 |
| | ARD | Interdependent | -135.7796 | -141.0934 | 287.5593 | 290.1869 | 10.6276 | **0.0311** | 0.0852 | -137.1743 | -145.4851 | 290.3485 | 298.9702 | 16.6217 | **0.0023** | **0.0104** | -139.3777 | -147.4055 | 294.7553 | 302.8109 | 16.0556 | **0.0029** | **0.0134** |
| 3 vs. 12 | ARD | Component 3 | -131.2480 | -137.4021 | 274.4960 | 282.8042 | 12.3082 | **0.0021** | **0.0123** | -133.5108 | -141.1689 | 279.0216 | 290.3378 | 15.3162 | **0.0005** | **0.0031** | -138.1538 | -141.7726 | 288.3075 | 291.5451 | 7.2376 | **0.0268** | 0.0728 |
| | ARD | Component 12 | -134.8455 | -137.4021 | 281.6910 | 282.8042 | 5.1132 | 0.0776 | 0.1647 | -138.4273 | -141.1689 | 288.8545 | 290.3378 | 5.4833 | 0.0645 | 0.1411 | -139.1305 | -141.7726 | 290.2611 | 291.5451 | 5.2841 | 0.0712 | 0.1518 |
| | ARD | Interdependent | -131.7951 | -137.4021 | 279.5902 | 282.8042 | 11.2140 | **0.0253** | 0.0731 | -135.2791 | -141.1689 | 286.5582 | 290.3378 | 11.7796 | **0.0191** | 0.0537 | -136.5554 | -141.7726 | 289.1108 | 291.5451 | 10.4343 | **0.0337** | 0.0849 |
| 3 vs. 13 | ER | Component 3 | -151.4552 | -155.4383 | 308.9104 | 314.8766 | 7.9662 | **0.0048** | **0.0229** | -154.8390 | -159.5527 | 315.6779 | 323.1055 | 9.4275 | **0.0021** | **0.0097** | -156.8898 | -160.4308 | 319.7796 | 324.8616 | 7.0821 | **0.0078** | **0.0286** |
| | ER | Component 13 | -155.2870 | -155.4383 | 316.5740 | 314.8766 | 0.3026 | 0.5823 | 0.7082 | -159.2814 | -159.5527 | 324.5629 | 323.1055 | 0.5426 | 0.4614 | 0.5884 | -160.1897 | -160.4308 | 326.3793 | 324.8616 | 0.4823 | 0.4874 | 0.6241 |
| | ER | Interdependent | -151.3739 | -155.4383 | 310.7478 | 314.8766 | 8.1287 | **0.0172** | 0.0547 | -154.7853 | -159.5527 | 317.5706 | 323.1055 | 9.5349 | **0.0085** | **0.0291** | -156.8598 | -160.4308 | 321.7197 | 324.8616 | 7.1419 | **0.0281** | 0.0757 |
| 3 vs. 14 | ARD | Component 3 | -148.9373 | -150.9776 | 309.8747 | 309.9552 | 4.0805 | 0.1300 | 0.2452 | -151.6527 | -153.7629 | 315.3049 | 315.5258 | 4.2209 | 0.1212 | 0.2166 | -154.3354 | -156.9046 | 320.6708 | 321.8092 | 5.1384 | 0.0766 | 0.1595 |
| | ARD | Component 14 | -145.8543 | -150.9776 | 303.7087 | 309.9552 | 10.2465 | **0.0060** | **0.0259** | -148.5259 | -153.7629 | 309.0518 | 315.5258 | 10.4739 | **0.0053** | **0.0204** | -151.6413 | -156.9046 | 315.2826 | 321.8092 | 10.5266 | **0.0052** | **0.0213** |
| | ARD | Interdependent | -144.2483 | -150.9776 | 304.4966 | 309.9552 | 13.4586 | **0.0092** | **0.0362** | -147.4022 | -153.7629 | 310.8044 | 315.5258 | 12.7214 | **0.0127** | **0.0408** | -150.5828 | -156.9046 | 317.1657 | 321.8092 | 12.6435 | **0.0132** | **0.0429** |
| 3 vs. 15 | ARD | Component 3 | -114.5601 | -117.2901 | 241.1202 | 242.5803 | 5.4601 | 0.0652 | 0.1488 | -117.6960 | -120.3217 | 247.3920 | 248.6435 | 5.2515 | 0.0724 | 0.1505 | -118.5888 | -120.8064 | 249.1776 | 249.6127 | 4.4351 | 0.1089 | 0.2018 |
| | ARD | Component 15 | -112.2261 | -117.2901 | 236.4521 | 242.5803 | 10.1282 | **0.0063** | **0.0268** | -115.1881 | -120.3217 | 242.3763 | 248.6435 | 10.2672 | **0.0059** | **0.0219** | -115.8462 | -120.8064 | 243.6924 | 249.6127 | 9.9203 | **0.0070** | **0.0269** |
| | ARD | Interdependent | -110.9209 | -117.2901 | 237.8419 | 242.5803 | 12.7384 | **0.0126** | **0.0441** | -114.2768 | -120.3217 | 244.5537 | 248.6435 | 12.0898 | **0.0167** | **0.0487** | -114.9360 | -120.8064 | 245.8720 | 249.6127 | 11.7408 | **0.0194** | 0.0561 |
| 4 vs. 6 | ARD | Component 4 | -107.5510 | -110.7921 | 227.1021 | 229.5841 | 6.4820 | **0.0391** | 0.0988 | NT | NT | NT | NT | NT | NT | NT | NT | NT | NT | NT | NT | NT | NT |
| | ARD | Component 6 | -107.5115 | -110.7921 | 227.0231 | 229.5841 | 6.5611 | **0.0376** | 0.0971 | NT | NT | NT | NT | NT | NT | NT | NT | NT | NT | NT | NT | NT | NT |
| | ARD | Interdependent | -106.3934 | -110.7921 | 228.7867 | 229.5841 | 8.7975 | 0.0664 | 0.1505 | NT | NT | NT | NT | NT | NT | NT | NT | NT | NT | NT | NT | NT | NT |
| 4 vs. 7 | ARD | Component 4 | -130.1173 | -136.2736 | 272.2346 | 280.5471 | 12.3125 | **0.0021** | **0.0123** | -130.9866 | -137.1327 | 273.9732 | 282.2655 | 12.2922 | **0.0021** | **0.0097** | -131.0695 | -137.7481 | 274.1389 | 283.4963 | 13.3573 | **0.0013** | **0.0069** |
| | ARD | Component 7 | -127.5696 | -136.2736 | 267.1391 | 280.5471 | 17.4080 | **0.0002** | **0.0018** | -128.2277 | -137.1327 | 268.4554 | 282.2655 | 17.8100 | **0.0001** | **0.0008** | -128.6994 | -137.7481 | 269.3988 | 283.4963 | 18.0975 | **0.0001** | **0.0009** |
| | ARD | Interdependent | -126.7200 | -136.2736 | 269.4401 | 280.5471 | 19.1071 | **0.0007** | **0.0048** | -127.3053 | -137.1327 | 270.6106 | 282.2655 | 19.6549 | **0.0006** | **0.0034** | -127.7684 | -137.7481 | 271.5368 | 283.4963 | 19.9595 | **0.0005** | **0.0032** |
| 4 vs. 8 | ARD | Component 4 | -131.2355 | -134.9427 | 274.4710 | 277.8854 | 7.4144 | **0.0245** | 0.0715 | -132.8454 | -136.6428 | 277.6908 | 281.2855 | 7.5948 | **0.0224** | 0.0624 | -129.6575 | -133.5390 | 271.3150 | 275.0780 | 7.7630 | **0.0206** | 0.0579 |
| | ARD | Component 8 | -128.6870 | -134.9427 | 269.3741 | 277.8854 | 12.5113 | **0.0019** | **0.0115** | -129.2982 | -136.6428 | 270.5964 | 281.2855 | 14.6892 | **0.0006** | **0.0034** | -127.5930 | -133.5390 | 267.1860 | 275.0780 | 11.8921 | **0.0026** | **0.0126** |
| | ARD | Interdependent | -127.0201 | -134.9427 | 270.0402 | 277.8854 | 15.8452 | **0.0032** | **0.0168** | -126.8531 | -136.6428 | 269.7062 | 281.2855 | 19.5793 | **0.0006** | **0.0034** | -125.1968 | -133.5390 | 266.3935 | 275.0780 | 16.6845 | **0.0022** | **0.0108** |
| 4 vs. 10 | ARD | Component 4 | -100.8322 | -101.9333 | 213.6644 | 211.8667 | 2.2023 | 0.3325 | 0.4718 | -103.0709 | -104.1340 | 218.1419 | 216.2680 | 2.1261 | 0.3454 | 0.4778 | -102.0744 | -102.8306 | 216.1488 | 213.6612 | 1.5125 | 0.4694 | 0.6085 |
| | ARD | Component 10 | -98.9983 | -101.9333 | 209.9967 | 211.8667 | 5.8700 | 0.0531 | 0.1277 | -101.8147 | -104.1340 | 215.6294 | 216.2680 | 4.6386 | 0.0983 | 0.1900 | -101.2565 | -102.8306 | 214.5131 | 213.6612 | 3.1482 | 0.2072 | 0.3231 |
| | ARD | Interdependent | -98.0218 | -101.9333 | 212.0436 | 211.8667 | 7.8231 | 0.0983 | 0.1947 | -100.6683 | -104.1340 | 217.3367 | 216.2680 | 6.9313 | 0.1396 | 0.2403 | -100.1866 | -102.8306 | 216.3732 | 213.6612 | 5.2880 | 0.2590 | 0.3830 |
| 4 vs. 13 | ARD | Component 4 | -111.1443 | -113.7067 | 234.2886 | 235.4134 | 5.1248 | 0.0771 | 0.1647 | -115.2300 | -117.8420 | 242.4600 | 243.6839 | 5.2239 | 0.0734 | 0.1505 | -114.5166 | -116.8073 | 241.0332 | 241.6147 | 4.5815 | 0.1012 | 0.1944 |

| | | | | | | | | | | | | | | | | | | | | | | | |
|---|---|---|---|---|---|---|---|---|---|---|---|---|---|---|---|---|---|---|---|---|---|---|---|
| | ARD | Component 13 | -109.5217 | -113.7067 | 231.0433 | 235.4134 | 8.3701 | **0.0152** | 0.0504 | -112.7416 | -117.8420 | 237.4831 | 243.6839 | 10.2008 | **0.0061** | **0.0223** | -111.9025 | -116.8073 | 235.8050 | 241.6147 | 9.8097 | **0.0074** | **0.0278** |
| | ARD | Interdependent | -110.4934 | -113.7067 | 236.9867 | 235.4134 | 6.4267 | 0.1695 | 0.2934 | -114.5069 | -117.8420 | 245.0137 | 243.6839 | 6.6702 | 0.1544 | 0.2615 | -113.7977 | -116.8073 | 243.5953 | 241.6147 | 6.0194 | 0.1977 | 0.3114 |
| 4 vs. 14 | ARD | Component 4 | -107.0500 | -109.9184 | 226.1001 | 227.8368 | 5.7367 | 0.0568 | 0.1335 | -109.8000 | -112.4973 | 231.6000 | 232.9947 | 5.3947 | 0.0674 | 0.1435 | -110.9435 | -113.8295 | 233.8871 | 235.6590 | 5.7719 | 0.0558 | 0.1274 |
| | ARD | Component 14 | -105.4250 | -109.9184 | 222.8500 | 227.8368 | 8.9869 | **0.0112** | **0.0406** | -107.1998 | -112.4973 | 226.3997 | 232.9947 | 10.5950 | **0.0050** | **0.0197** | -109.2300 | -113.8295 | 230.4601 | 235.6590 | 9.1989 | **0.0101** | **0.0346** |
| | ARD | Interdependent | -104.7484 | -109.9184 | 225.4968 | 227.8368 | 10.3400 | **0.0351** | 0.0937 | -106.3950 | -112.4973 | 228.7900 | 232.9947 | 12.2047 | **0.0159** | **0.0473** | -108.4918 | -113.8295 | 232.9837 | 235.6590 | 10.6753 | **0.0305** | 0.0807 |
| 4 vs. 15 | ARD | Component 4 | -72.7088 | -76.2309 | 157.4176 | 160.4619 | 7.0442 | **0.0295** | 0.0818 | -74.8882 | -79.0562 | 161.7763 | 166.1124 | 8.3361 | **0.0155** | **0.0465** | -73.8355 | -77.7312 | 159.6711 | 163.4625 | 7.7914 | **0.0203** | 0.0576 |
| | ARD | Component 15 | -69.9886 | -76.2309 | 151.9772 | 160.4619 | 12.4846 | **0.0019** | **0.0115** | -72.1276 | -79.0562 | 156.2552 | 166.1124 | 13.8572 | **0.0010** | **0.0053** | -70.9585 | -77.7312 | 153.9170 | 163.4625 | 13.5455 | **0.0011** | **0.0062** |
| | ARD | Interdependent | -69.9840 | -76.2309 | 155.9680 | 160.4619 | 12.4939 | **0.0140** | **0.0478** | -72.0689 | -79.0562 | 160.1378 | 166.1124 | 13.9745 | **0.0074** | **0.0262** | -71.2032 | -77.7312 | 158.4064 | 163.4625 | 13.0560 | **0.0110** | **0.0369** |
| 5 vs. 6 | ER | Component 5 | -102.0758 | -104.0799 | 210.1517 | 212.1598 | 4.0081 | **0.0453** | 0.1133 | NT | NT | NT | NT | NT | NT | NT | NT | NT | NT | NT | NT | NT | NT |
| | ER | Component 6 | -103.9109 | -104.0799 | 213.8217 | 212.1598 | 0.3381 | 0.5609 | 0.6956 | NT | NT | NT | NT | NT | NT | NT | NT | NT | NT | NT | NT | NT | NT |
| | ER | Interdependent | -101.5850 | -104.0799 | 211.1700 | 212.1598 | 4.9898 | 0.0825 | 0.1699 | NT | NT | NT | NT | NT | NT | NT | NT | NT | NT | NT | NT | NT | NT |
| 5 vs. 7 | ARD | Component 5 | -125.1476 | -128.3860 | 262.2952 | 264.7720 | 6.4767 | **0.0392** | 0.0988 | -124.7492 | -128.4869 | 261.4984 | 264.9738 | 7.4754 | **0.0238** | 0.0652 | -125.6562 | -129.7355 | 263.3125 | 267.4709 | 8.1584 | 0.1692 | 0.2805 |
| | ARD | Component 7 | -124.1469 | -128.3860 | 260.2937 | 264.7720 | 8.4782 | **0.0144** | **0.0483** | -123.9226 | -128.4869 | 259.8452 | 264.9738 | 9.1286 | **0.0104** | **0.0345** | -124.7634 | -129.7355 | 261.5269 | 267.4709 | 9.9440 | **0.0069** | **0.0268** |
| | ARD | Interdependent | -123.9410 | -128.3860 | 263.8820 | 264.7720 | 8.8900 | 0.0639 | 0.1469 | -123.6584 | -128.4869 | 263.3169 | 264.9738 | 9.6569 | **0.0466** | 0.1121 | -124.4312 | -129.7355 | 264.8624 | 267.4709 | 10.6085 | 0.3133 | 0.4487 |
| 5 vs. 8 | ARD | Component 5 | -122.9338 | -127.0551 | 257.8676 | 262.1103 | 8.2426 | **0.0162** | 0.0521 | -123.5519 | -127.9969 | 259.1038 | 263.9938 | 8.8900 | **0.0117** | **0.0383** | -120.5014 | -125.5263 | 253.0028 | 259.0527 | 10.0499 | **0.0066** | **0.0260** |
| | ARD | Component 8 | -121.9107 | -127.0551 | 255.8213 | 262.1103 | 10.2889 | **0.0058** | **0.0254** | -122.4348 | -127.9969 | 256.8696 | 263.9938 | 11.1242 | **0.0038** | **0.0158** | -119.8964 | -125.5263 | 251.7928 | 259.0527 | 11.2599 | **0.0036** | **0.0162** |
| | ARD | Interdependent | -121.5607 | -127.0551 | 259.1215 | 262.1103 | 10.9888 | **0.0267** | 0.0765 | -123.5121 | -127.9969 | 263.0241 | 263.9938 | 8.9697 | 0.0619 | 0.1383 | -121.1662 | -125.5263 | 258.3324 | 259.0527 | 8.7203 | 0.0685 | 0.1498 |
| 6 vs. 7 | ARD | Component 6 | -158.7816 | -168.4826 | 329.5632 | 344.9652 | 19.4019 | **0.0001** | **0.0010** | NT | NT | NT | NT | NT | NT | NT | NT | NT | NT | NT | NT | NT | NT |
| | ARD | Component 7 | -159.3086 | -168.4826 | 330.6172 | 344.9652 | 18.3479 | **0.0001** | **0.0010** | NT | NT | NT | NT | NT | NT | NT | NT | NT | NT | NT | NT | NT | NT |
| | ARD | Interdependent | -154.7614 | -168.4826 | 325.5229 | 344.9652 | 27.4423 | **0.0000** | **0.0000** | NT | NT | NT | NT | NT | NT | NT | NT | NT | NT | NT | NT | NT | NT |
| 6 vs. 8 | ARD | Component 6 | -157.8613 | -167.1517 | 327.7226 | 342.3035 | 18.5808 | **0.0001** | **0.0010** | NT | NT | NT | NT | NT | NT | NT | NT | NT | NT | NT | NT | NT | NT |
| | ARD | Component 8 | -156.9342 | -167.1517 | 325.8684 | 342.3035 | 20.4351 | **0.0000** | **0.0000** | NT | NT | NT | NT | NT | NT | NT | NT | NT | NT | NT | NT | NT | NT |
| | ARD | Interdependent | -154.3025 | -167.1517 | 324.6050 | 342.3035 | 25.6985 | **0.0000** | **0.0000** | NT | NT | NT | NT | NT | NT | NT | NT | NT | NT | NT | NT | NT | NT |
| 6 vs. 9 | ARD | Component 6 | -116.9187 | -117.7387 | 245.8373 | 243.4773 | 1.6400 | 0.4404 | 0.5804 | NT | NT | NT | NT | NT | NT | NT | NT | NT | NT | NT | NT | NT | NT |
| | ARD | Component 9 | -114.8489 | -117.7387 | 241.6978 | 243.4773 | 5.7795 | 0.0556 | 0.1327 | NT | NT | NT | NT | NT | NT | NT | NT | NT | NT | NT | NT | NT | NT |
| | ARD | Interdependent | -114.8870 | -117.7387 | 245.7740 | 243.4773 | 5.7034 | 0.2224 | 0.3611 | NT | NT | NT | NT | NT | NT | NT | NT | NT | NT | NT | NT | NT | NT |
| 6 vs. 11 | ARD | Component 6 | -128.2394 | -132.2432 | 268.4787 | 272.4865 | 8.0078 | **0.0182** | 0.0573 | NT | NT | NT | NT | NT | NT | NT | NT | NT | NT | NT | NT | NT | NT |
| | ARD | Component 11 | -125.6025 | -132.2432 | 263.2051 | 272.4865 | 13.2814 | **0.0013** | **0.0084** | NT | NT | NT | NT | NT | NT | NT | NT | NT | NT | NT | NT | NT | NT |
| | ARD | Interdependent | -123.9097 | -132.2432 | 263.8194 | 272.4865 | 16.6671 | **0.0022** | **0.0126** | NT | NT | NT | NT | NT | NT | NT | NT | NT | NT | NT | NT | NT | NT |
| 6 vs. 12 | ARD | Component 6 | -125.7549 | -128.5519 | 263.5099 | 265.1038 | 5.5940 | 0.0610 | 0.1413 | NT | NT | NT | NT | NT | NT | NT | NT | NT | NT | NT | NT | NT | NT |
| | ARD | Component 12 | -123.9572 | -128.5519 | 259.9145 | 265.1038 | 9.1894 | **0.0101** | **0.0383** | NT | NT | NT | NT | NT | NT | NT | NT | NT | NT | NT | NT | NT | NT |
| | ARD | Interdependent | -123.8529 | -128.5519 | 263.7059 | 265.1038 | 9.3979 | 0.0519 | 0.1267 | NT | NT | NT | NT | NT | NT | NT | NT | NT | NT | NT | NT | NT | NT |
| 6 vs. 13 | ER | Component 6 | -140.6797 | -145.9363 | 287.3595 | 295.8727 | 10.5132 | **0.0012** | **0.0079** | NT | NT | NT | NT | NT | NT | NT | NT | NT | NT | NT | NT | NT | NT |
| | ER | Component 13 | -140.0523 | -145.9363 | 286.1047 | 295.8727 | 11.7680 | **0.0006** | **0.0042** | NT | NT | NT | NT | NT | NT | NT | NT | NT | NT | NT | NT | NT | NT |
| | ER | Interdependent | -133.0696 | -145.9363 | 274.1393 | 295.8727 | 25.7334 | **0.0000** | **0.0000** | NT | NT | NT | NT | NT | NT | NT | NT | NT | NT | NT | NT | NT | NT |
| 6 vs. 14 | ARD | Component 6 | -138.5261 | -142.1274 | 289.0521 | 292.2548 | 7.2027 | **0.0273** | 0.0775 | NT | NT | NT | NT | NT | NT | NT | NT | NT | NT | NT | NT | NT | NT |
| | ARD | Component 14 | -136.8429 | -142.1274 | 285.6858 | 292.2548 | 10.5691 | **0.0051** | **0.0233** | NT | NT | NT | NT | NT | NT | NT | NT | NT | NT | NT | NT | NT | NT |
| | ARD | Interdependent | -136.0325 | -142.1274 | 288.0649 | 292.2548 | 12.1899 | **0.0160** | 0.0520 | NT | NT | NT | NT | NT | NT | NT | NT | NT | NT | NT | NT | NT | NT |
| 6 vs. 15 | ARD | Component 6 | -105.4926 | 108.4400 | 222.9852 | 224.8799 | 5.8947 | 0.0525 | 0.1272 | NT | NT | NT | NT | NT | NT | NT | NT | NT | NT | NT | NT | NT | NT |
| | ARD | Component 15 | -106.4857 | 108.4400 | 224.9714 | 224.8799 | 3.9086 | 0.1417 | 0.2601 | NT | NT | NT | NT | NT | NT | NT | NT | NT | NT | NT | NT | NT | NT |
| | ARD | Interdependent | -105.2350 | 108.4400 | 226.4700 | 224.8799 | 6.4099 | 0.1706 | 0.2937 | NT | NT | NT | NT | NT | NT | NT | NT | NT | NT | NT | NT | NT | NT |
| 7 vs. 8 | ARD | Component 7 | -156.6934 | -192.6332 | 325.3867 | 393.2665 | 71.8797 | **0.0000** | **0.0000** | -156.6834 | -192.6672 | 325.3668 | 393.3344 | 71.9676 | **0.0000** | **0.0000** | -154.9149 | -191.5323 | 321.8299 | 391.0645 | 73.2346 | **0.0000** | **0.0000** |
| | ARD | Component 8 | -154.4443 | -192.6332 | 320.8885 | 393.2665 | 76.3779 | **0.0000** | **0.0000** | -154.1116 | -192.6672 | 320.2233 | 393.3344 | 77.1111 | **0.0000** | **0.0000** | -154.0003 | -191.5323 | 320.0006 | 391.0645 | 75.0639 | **0.0000** | **0.0000** |
| | ARD | Interdependent | -153.2075 | -192.6332 | 322.4150 | 393.2665 | 78.8515 | **0.0000** | **0.0000** | -152.9186 | -192.6672 | 321.8373 | 393.3344 | 79.4971 | **0.0000** | **0.0000** | -152.5472 | -191.5323 | 321.0944 | 391.0645 | 77.9701 | **0.0000** | **0.0000** |
| 7 vs. 13 | ARD | Component 7 | -159.9020 | -171.3972 | 331.8041 | 350.7945 | 22.9904 | **0.0000** | **0.0000** | -162.8760 | -173.8664 | 337.7520 | 355.7328 | 21.9809 | **0.0000** | **0.0000** | -163.2087 | -174.8006 | 338.4174 | 357.6011 | 23.1838 | **0.0000** | **0.0000** |
| | ARD | Component 13 | -162.2937 | -171.3972 | 336.5873 | 350.7945 | 18.2071 | **0.0001** | **0.0010** | -164.3966 | -173.8664 | 340.7931 | 355.7328 | 18.9397 | **0.0001** | **0.0008** | -165.8587 | -174.8006 | 343.7173 | 357.6011 | 17.8838 | **0.0001** | **0.0009** |
| | ARD | Interdependent | -158.1532 | -171.3972 | 332.3065 | 350.7945 | 26.4880 | **0.0000** | **0.0000** | -160.6084 | -173.8664 | 337.2168 | 355.7328 | 26.5160 | **0.0000** | **0.0000** | -161.5275 | -174.8006 | 339.0549 | 357.6011 | 26.5462 | **0.0000** | **0.0000** |
| 7 vs. 14 | ARD | Component 7 | -162.7974 | -167.6089 | 337.5949 | 343.2178 | 9.6230 | **0.0081** | **0.0331** | -163.9165 | -168.5218 | 339.8330 | 345.0435 | 9.2106 | **0.0100** | **0.0335** | -166.6433 | -171.8227 | 345.2866 | 351.6454 | 10.3589 | **0.0056** | **0.0223** |
| | ARD | Component 14 | -159.4243 | -167.6089 | 330.8485 | 343.2178 | 16.3693 | **0.0003** | **0.0025** | -159.8256 | -168.5218 | 331.6512 | 345.0435 | 17.3923 | **0.0002** | **0.0015** | -162.6237 | -171.8227 | 337.2474 | 351.6454 | 18.3980 | **0.0001** | **0.0009** |
| | ARD | Interdependent | -157.5467 | -167.6089 | 331.0934 | 343.2178 | 20.1244 | **0.0005** | **0.0038** | -157.3201 | -168.5218 | 330.6403 | 345.0435 | 22.4032 | **0.0002** | **0.0015** | -162.0648 | -171.8227 | 340.1295 | 351.6454 | 19.5159 | **0.0006** | **0.0038** |
| 8 vs. 11 | ARD | Component 8 | -154.5562 | -156.3939 | 321.1123 | 320.7878 | 3.6755 | 0.1592 | 0.2802 | -156.7575 | -159.7540 | 325.5151 | 327.5081 | 5.9930 | **0.0500** | 0.1167 | -153.9591 | -158.1145 | 319.9182 | 324.2289 | 8.3107 | **0.0157** | **0.0486** |
| | ARD | Component 11 | -155.2410 | -156.3939 | 322.4820 | 320.7878 | 2.3058 | 0.3157 | 0.4569 | -156.7117 | -159.7540 | 325.4233 | 327.5081 | 6.0848 | **0.0477** | 0.1138 | -155.5588 | -158.1145 | 323.1177 | 324.2289 | 5.1113 | 0.0776 | 0.1595 |
| | ARD | Interdependent | -154.4880 | -156.3939 | 324.9760 | 320.7878 | 3.8118 | 0.4321 | 0.5719 | -158.1318 | -159.7540 | 332.2635 | 327.5081 | 3.2446 | 0.5178 | 0.6473 | -154.6277 | -158.1145 | 325.2554 | 324.2289 | 6.9736 | 0.1373 | 0.2430 |
| 8 vs. 12 | ARD | Component 8 | -152.2099 | -152.7026 | 316.4198 | 313.4051 | 0.9853 | 0.6110 | 0.7290 | -155.2704 | -155.4378 | 322.5409 | 318.8756 | 0.3348 | 0.8459 | 0.9032 | -152.1427 | -152.4816 | 316.2854 | 312.9631 | 0.6777 | 0.7126 | 0.8074 |
| | ARD | Component 12 | -151.3640 | -152.7026 | 314.7281 | 313.4051 | 2.6770 | 0.2622 | 0.4007 | -153.8214 | -155.4378 | 319.6427 | 318.8756 | 3.2329 | 0.1986 | 0.3112 | -151.9043 | -152.4816 | 315.8086 | 312.9631 | 1.1545 | 0.5614 | 0.6956 |
| | ARD | Interdependent | -151.2703 | -152.7026 | 318.5405 | 313.4051 | 2.8646 | 0.5807 | 0.7082 | -153.7889 | -155.4378 | 323.5778 | 318.8756 | 3.2979 | 0.5093 | 0.6417 | -151.7824 | -152.4816 | 319.5648 | 312.9631 | 1.3983 | 0.8445 | 0.8954 |
| 8 vs. 13 | ARD | Component 8 | -160.4011 | -170.0664 | 332.8022 | 348.1328 | 19.3306 | **0.0001** | **0.0010** | -164.1321 | -173.3764 | 340.2643 | 354.7529 | 18.4886 | **0.0001** | **0.0008** | -162.1245 | -170.5914 | 336.2490 | 349.1829 | 16.9339 | **0.0002** | **0.0015** |
| | ARD | Component 13 | -162.7020 | -170.0664 | 337.4040 | 348.1328 | 14.7288 | **0.0006** | **0.0042** | -166.0131 | -173.3764 | 344.0263 | 354.7529 | 14.7266 | **0.0006** | **0.0034** | 164.1298 | -170.5914 | 340.2595 | 349.1829 | 12.9234 | **0.0016** | **0.0083** |
| | ARD | Interdependent | -156.9631 | -170.0664 | 329.9262 | 348.1328 | 26.2066 | **0.0000** | **0.0000** | -161.7627 | -173.3764 | 339.5255 | 354.7529 | 23.2274 | **0.0001** | **0.0008** | -160.2001 | -170.5914 | 336.4003 | 349.1829 | 20.7826 | **0.0003** | **0.0021** |
| 8 vs. 14 | ARD | Component 8 | -153.3732 | -166.2781 | 318.7464 | 340.5561 | 25.8098 | **0.0000** | **0.0000** | -154.8654 | -168.0318 | 321.7308 | 344.0636 | 26.3328 | **0.0000** | **0.0000** | -153.9335 | -167.6136 | 319.8670 | 343.2272 | 27.3602 | **0.0000** | **0.0000** |
| | ARD | Component 14 | -150.1657 | -166.2781 | 312.3313 | 340.5561 | 32.2248 | **0.0000** | **0.0000** | -151.4168 | -168.0318 | 314.8337 | 344.0636 | 33.2299 | **0.0000** | **0.0000** | -151.7985 | -167.6136 | 315.5969 | 343.2272 | 31.6303 | **0.0000** | **0.0000** |
| | ARD | Interdependent | -146.6523 | -166.2781 | 309.3047 | 340.5561 | 39.2515 | **0.0000** | **0.0000** | -147.2156 | -168.0318 | 310.4312 | 344.0636 | 41.6324 | **0.0000** | **0.0000** | -146.6628 | -167.6136 | 309.3256 | 343.2272 | 41.9017 | **0.0000** | **0.0000** |
| 9 vs. 10 | ARD | Component 9 | -103.9890 | -107.8717 | 219.9781 | 223.7434 | 7.7653 | **0.0206** | 0.0618 | -104.0834 | -109.7234 | 220.1668 | 227.4469 | 11.2801 | **0.0036** | **0.0151** | -102.9543 | -108.6083 | 217.9085 | 225.2165 | 11.3080 | **0.0035** | **0.0160** |
| | ARD | Component 10 | -98.6741 | -107.8717 | 209.3481 | 223.7434 | 18.3953 | **0.0001** | **0.0010** | -100.7844 | -109.7234 | 213.5687 | 227.4469 | 17.8782 | **0.0001** | **0.0008** | -99.5116 | -108.6083 | 211.0231 | 225.2165 | 18.1934 | **0.0001** | **0.0009** |
| | ARD | Interdependent | -96.2781 | -107.8717 | 208.5561 | 223.7434 | 23.1872 | **0.0001** | **0.0010** | -98.1795 | -109.7234 | 212.3589 | 227.4469 | 23.0879 | **0.0001** | **0.0008** | -96.8018 | -108.6083 | 209.6036 | 225.2165 | 23.6129 | **0.0001** | **0.0009** |
| 9 vs. 13 | ARD | Component 9 | -114.5795 | -120.6533 | 241.1590 | 249.3066 | 12.1476 | **0.0023** | **0.0129** | -117.7792 | -124.6090 | 247.5583 | 257.2179 | 13.6596 | **0.0011** | **0.0057** | -116.3635 | -123.5711 | 244.7271 | 255.1422 | 14.4151 | **0.0007** | **0.0042** |
| | ARD | Component 13 | -106.9999 | -120.6533 | 225.9998 | 249.3066 | 27.3068 | **0.0000** | **0.0000** | -109.3689 | -124.6090 | 230.7378 | 257.2179 | 30.4802 | **0.0000** | **0.0000** | -108.7144 | -123.5711 | 229.4289 | 255.1422 | 29.7133 | **0.0000** | **0.0000** |
| | ARD | Interdependent | -106.9446 | -120.6533 | 229.8892 | 249.3066 | 27.4174 | **0.0000** | **0.0000** | -109.2205 | -124.6090 | 234.4410 | 257.2179 | 30.7769 | **0.0000** | **0.0000** | -108.5791 | -123.5711 | 233.1582 | 255.1422 | 29.9839 | **0.0000** | **0.0000** |
| 9 vs. 14 | ARD | Component 9 | -114.3149 | -115.8568 | 240.6298 | 239.7135 | 3.0838 | 0.2140 | 0.3529 | -116.3580 | -118.1802 | 244.7160 | 244.3605 | 3.6445 | 0.1617 | 0.2661 | -120.5226 | -119.6071 | 253.0451 | 247.2142 | -1.8309 | 1.0000 | 1.0000 |
| | ARD | Component 14 | -115.4383 | -115.8568 | 242.8766 | 239.7135 | 0.8370 | 0.6580 | 0.7705 | -117.0994 | -118.1802 | 246.1987 | 244.3605 | 2.1618 | 0.3393 | 0.4729 | -118.4468 | -119.6071 | 248.8936 | 247.2142 | 2.3206 | 0.3134 | 0.4487 |
| | ARD | Interdependent | -115.4367 | -115.8568 | 246.8734 | 239.7135 | 0.8401 | 0.9330 | 0.9604 | -117.0885 | -118.1802 | 250.1770 | 244.3605 | 2.1835 | 0.7021 | 0.8083 | -118.4005 | -119.6071 | 252.8011 | 247.2142 | 2.4132 | 0.6602 | 0.7789 |
| 9 vs. 15 | ARD | Component 9 | -79.3123 | -82.1693 | 170.6245 | 172.3386 | 5.7141 | 0.0574 | 0.1339 | -81.4609 | -84.7391 | 174.9218 | 177.4782 | 6.5564 | **0.0377** | 0.0928 | -80.3047 | -83.5089 | 172.6095 | 175.0177 | 6.4083 | **0.0406** | 0.0991 |
| | ARD | Component 15 | -81.3709 | -82.1693 | 174.7417 | 172.3386 | 1.5969 | 0.4500 | 0.5906 | -84.4001 | -84.7391 | 180.8001 | 177.4782 | 0.6781 | 0.7125 | 0.8124 | -83.2535 | -83.5089 | 178.5071 | 175.0177 | 0.5107 | 0.7747 | 0.8562 |
| | ARD | Interdependent | -80.7979 | -82.1693 | 177.5957 | 172.3386 | 2.7429 | 0.6017 | 0.7219 | -83.1842 | -84.7391 | 182.3683 | 177.4782 | 3.1099 | 0.5396 | 0.6658 | -82.5671 | -83.5089 | 181.1343 | 175.0177 | 1.8835 | 0.7572 | 0.8428 |
| 10 vs. 13 | ER | Component 10 | -124.3809 | -138.8809 | 254.7617 | 281.7618 | 29.0001 | **0.0000** | **0.0000** | -126.4449 | -141.5975 | 258.8897 | 287.1950 | 30.3053 | **0.0000** | **0.0000** | -125.8026 | -140.5164 | 257.6053 | 285.0328 | 29.4276 | **0.0000** | **0.0000** |
| | ER | Component 13 | -138.8627 | -138.8809 | 283.7253 | 281.7618 | 0.0365 | 0.8485 | 0.8919 | -141.5593 | -141.5975 | 289.1187 | 287.1950 | 0.0764 | 0.7823 | 0.8675 | -140.4445 | -140.5164 | 286.8890 | 285.0328 | 0.1438 | 0.7045 | 0.8057 |
| | ER | Interdependent | -121.9435 | -138.8809 | 251.8869 | 281.7618 | 33.8749 | **0.0000** | **0.0000** | -123.9686 | -141.5975 | 255.9372 | 287.1950 | 35.2578 | **0.0000** | **0.0000** | -123.2649 | -140.5164 | 254.5298 | 285.0328 | 34.5030 | **0.0000** | **0.0000** |
| 11 vs. 12 | ARD | Component 11 | -88.7474 | -117.6732 | 189.4948 | 243.3465 | 57.8517 | **0.0000** | **0.0000** | -92.0053 | -121.2265 | 196.0106 | 250.4529 | 58.4423 | **0.0000** | **0.0000** | -91.7683 | -121.4358 | 195.5365 | 250.8715 | 59.3350 | **0.0000** | **0.0000** |
| | ARD | Component 12 | -86.2068 | -117.6732 | 184.4135 | 243.3465 | 62.9330 | **0.0000** | **0.0000** | -89.3382 | -121.2265 | 190.6765 | 250.4529 | 63.7764 | **0.0000** | **0.0000** | -88.6782 | -121.4358 | 189.3563 | 250.8715 | 65.5152 | **0.0000** | **0.0000** |
| | ARD | Interdependent | -86.1244 | -117.6732 | 188.2489 | 243.3465 | 63.0976 | **0.0000** | **0.0000** | -89.2450 | -121.2265 | 194.4899 | 250.4529 | 63.9630 | **0.0000** | **0.0000** | -88.4842 | -121.4358 | 192.9684 | 250.8715 | 65.9032 | **0.0000** | **0.0000** |
| 11 vs. 13 | ARD | Component 11 | -128.8295 | -135.1579 | 269.6591 | 278.3158 | 12.6567 | **0.0018** | **0.0113** | -130.9359 | -140.9532 | 273.8718 | 289.9065 | 20.0347 | **0.0000** | **0.0000** | -132.2924 | 141.3828 | 276.5848 | 290.7655 | 18.1807 | **0.0001** | **0.0009** |
| | ARD | Component 13 | -124.4855 | -135.1579 | 260.9709 | 278.3158 | 21.3449 | **0.0000** | **0.0000** | -127.4903 | -140.9532 | 266.9807 | 289.9065 | 26.9258 | **0.0000** | **0.0000** | -127.7728 | 141.3828 | 267.5456 | 290.7655 | 27.2199 | **0.0000** | **0.0000** |
| | ARD | Interdependent | -124.4662 | -135.1579 | 264.9325 | 278.3158 | 21.3833 | **0.0003** | **0.0025** | -129.5115 | -140.9532 | 275.0230 | 289.9065 | 22.8835 | **0.0001** | **0.0008** | -130.5666 | 141.3828 | 277.1333 | 290.7655 | 21.6323 | **0.0002** | **0.0015** |
| 11 vs. 14 | ARD | Component 11 | -130.6101 | -131.3696 | 273.2201 | 270.7392 | 1.5191 | 0.4679 | 0.6090 | -132.7772 | -133.9735 | 277.5543 | 275.9471 | 2.3927 | 0.3023 | 0.4368 | -135.2010 | -136.3740 | 282.4021 | 280.7480 | 2.3459 | 0.3094 | 0.4471 |
| | ARD | Component 14 | -131.0389 | -131.3696 | 274.0778 | 270.7392 | 0.6614 | 0.7184 | 0.8103 | -134.6720 | -133.9735 | 281.3439 | 275.9471 | -1.3969 | 1.0000 | 1.0000 | -135.6776 | -136.3740 | 283.3552 | 280.7480 | 1.3928 | 0.4984 | 0.6330 |
| | ARD | Interdependent | -130.5252 | -131.3696 | 277.0503 | 270.7392 | 1.6889 | 0.7927 | 0.8543 | -134.6648 | -133.9735 | 285.3296 | 275.9471 | -1.3825 | 1.0000 | 1.0000 | -137.0965 | -136.3740 | 290.1929 | 280.7480 | -1.4449 | 1.0000 | 1.0000 |
| 12 vs. 13 | ARD | Component 12 | -125.4262 | -131.4666 | 262.8525 | 270.9331 | 12.0807 | **0.0024** | **0.0133** | -128.8298 | -136.6370 | 269.6596 | 281.2740 | 15.6144 | **0.0004** | **0.0026** | -128.7751 | -135.7499 | 269.5502 | 279.4998 | 13.9496 | **0.0009** | **0.0053** |
| | ARD | Component 13 | -123.0559 | -131.4666 | 258.1118 | 270.9331 | 16.8214 | **0.0002** | **0.0018** | -127.2669 | -136.6370 | 266.5338 | 281.2740 | 18.7402 | **0.0001** | **0.0008** | -127.0555 | -135.7499 | 266.1110 | 279.4998 | 17.3887 | **0.0002** | **0.0015** |
| | ARD | Interdependent | -122.1827 | -131.4666 | 260.3655 | 270.9331 | 18.5677 | **0.0010** | **0.0067** | -126.4029 | -136.6370 | 268.8057 | 281.2740 | 20.4683 | **0.0004** | **0.0026** | -126.5743 | -135.7499 | 269.1485 | 279.4998 | 18.3512 | **0.0011** | **0.0062** |
| 12 vs. 14 | ARD | Component 12 | -127.0510 | -127.5574 | 266.1021 | 263.1148 | 1.0128 | 0.6027 | 0.7219 | -130.2301 | -131.1393 | 272.4603 | 270.2786 | 1.8183 | 0.4029 | 0.5333 | -132.3421 | -132.9658 | 276.6842 | 273.9317 | 1.2474 | 0.5359 | 0.6725 |
| | ARD | Component 14 | -126.3060 | -127.5574 | 264.6119 | 263.1148 | 2.5029 | 0.2861 | 0.4251 | -129.6211 | -131.1393 | 271.2421 | 270.2786 | 3.0365 | 0.2191 | 0.3334 | -130.6567 | -132.9658 | 273.3135 | 273.9317 | 4.6182 | 0.0994 | 0.1933 |

| | | | | | | | | | | | | | | | | | | | | | | |
|---|---|---|---|---|---|---|---|---|---|---|---|---|---|---|---|---|---|---|---|---|---|---|
| | ARD | Interdependent | -126.1076 | -127.5574 | 268.2153 | 263.1148 | 2.8996 | 0.5748 | 0.7045 | -128.8406 | -131.1393 | 273.6811 | 270.2786 | 4.5975 | 0.3311 | 0.4688 | -130.3211 | -132.9658 | 276.6421 | 273.9317 | 5.2896 | 0.2589 | 0.3830 |
| 12 vs. 15 | ARD | Component 12 | -92.4331 | -93.8699 | 196.8661 | 195.7399 | 2.8737 | 0.2377 | 0.3763 | -95.9586 | -97.6982 | 203.9173 | 203.3963 | 3.4790 | 0.1756 | 0.2837 | -95.3750 | -96.8676 | 202.7499 | 201.7352 | 2.9852 | 0.2248 | 0.3454 |
| | ARD | Component 15 | -92.0912 | -93.8699 | 196.1824 | 195.7399 | 3.5574 | 0.1689 | 0.2934 | -94.9911 | -97.6982 | 201.9822 | 203.3963 | 5.4142 | 0.0667 | 0.1429 | -94.3538 | -96.8676 | 200.7076 | 201.7352 | 5.0276 | 0.0810 | 0.1636 |
| | ARD | Interdependent | -90.4235 | -93.8699 | 196.8470 | 195.7399 | 6.8929 | 0.1417 | 0.2601 | -93.5983 | -97.6982 | 203.1965 | 203.3963 | 8.1998 | 0.0845 | 0.1695 | -92.9536 | -96.8676 | 201.9073 | 201.7352 | 7.8279 | 0.0981 | 0.1919 |
| 13 vs. 14 | ARD | Component 13 | -140.5112 | -145.0421 | 293.0225 | 298.0842 | 9.0617 | **0.0108** | **0.0396** | -143.1444 | -149.2310 | 298.2888 | 306.4620 | 12.1732 | **0.0023** | **0.0104** | -145.4648 | -150.8819 | 302.9296 | 309.7638 | 10.8342 | **0.0044** | **0.0193** |
| | ARD | Component 14 | -141.7347 | -145.0421 | 295.4695 | 298.0842 | 6.6147 | **0.0366** | 0.0953 | -145.8210 | -149.2310 | 303.6419 | 306.4620 | 6.8201 | **0.0330** | 0.0866 | -147.2452 | -150.8819 | 306.4905 | 309.7638 | 7.2734 | **0.0263** | 0.0720 |
| | ARD | Interdependent | -140.9858 | -145.0421 | 297.9715 | 298.0842 | 8.1126 | 0.0875 | 0.1785 | -141.5421 | -149.2310 | 299.0842 | 306.4620 | 15.3778 | **0.0040** | **0.0164** | -146.5158 | -150.8819 | 309.0316 | 309.7638 | 8.7322 | 0.0682 | 0.1498 |
| 13 vs. 15 | ARD | Component 13 | -107.4720 | -111.3546 | 226.9440 | 230.7092 | 7.7652 | **0.0206** | 0.0618 | -108.7113 | -115.7899 | 229.4225 | 239.5797 | 14.1572 | **0.0008** | **0.0043** | -110.6434 | -114.7837 | 233.2868 | 237.5673 | 8.2805 | **0.0159** | **0.0486** |
| | ARD | Component 15 | -107.7755 | -111.3546 | 227.5511 | 230.7092 | 7.1582 | **0.0279** | 0.0785 | -111.7291 | -115.7899 | 235.4582 | 239.5797 | 8.1215 | **0.0172** | **0.0493** | -110.8251 | -114.7837 | 233.6501 | 237.5673 | 7.9172 | **0.0191** | 0.0557 |
| | ARD | Interdependent | -104.5175 | -111.3546 | 225.0349 | 230.7092 | 13.6743 | **0.0084** | **0.0339** | -107.0816 | -115.7899 | 230.1631 | 239.5797 | 17.4166 | **0.0016** | **0.0079** | -106.7467 | -114.7837 | 229.4935 | 237.5673 | 16.0738 | **0.0029** | **0.0134** |
| 14 vs. 15 | ARD | Component 14 | -98.0675 | -107.5663 | 208.1350 | 223.1326 | 18.9976 | **0.0001** | **0.0010** | -99.6835 | -110.4452 | 211.3669 | 228.8905 | 21.5236 | **0.0000** | **0.0000** | -100.6380 | -111.8058 | 213.2760 | 231.6116 | 22.3357 | **0.0000** | **0.0000** |
| | ARD | Component 15 | -99.7018 | -107.5663 | 211.4037 | 223.1326 | 15.7289 | **0.0004** | **0.0032** | -101.5432 | -110.4452 | 215.0865 | 228.8905 | 17.8040 | **0.0001** | **0.0008** | -103.1916 | -111.8058 | 218.3831 | 231.6116 | 17.2285 | **0.0002** | **0.0015** |
| | ARD | Interdependent | -97.6350 | -107.5663 | 211.2700 | 223.1326 | 19.8626 | **0.0005** | **0.0038** | -99.0841 | -110.4452 | 214.1682 | 228.8905 | 22.7223 | **0.0001** | **0.0008** | -99.5634 | -111.8058 | 215.1267 | 231.6116 | 24.4849 | **0.0001** | **0.0009** |

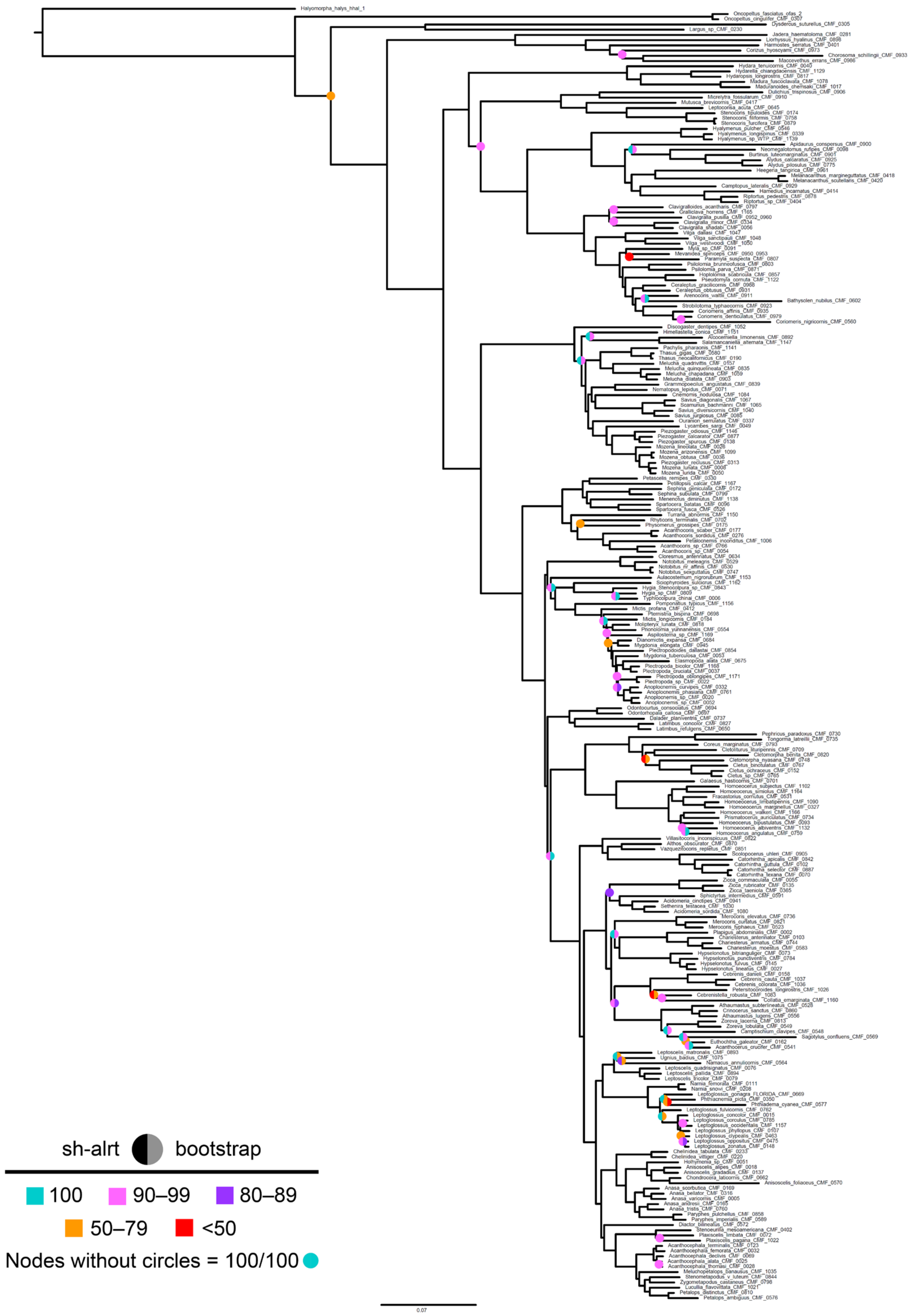


Figure S1. Maximum likelihood (ML) best tree based on the 50p concatenated alignment. Circles at nodes represent instances when Shimodaira-Hasegawa-like approximate likelihood ratio test (sh-alrt) and/or bootstrap support are less than 100.

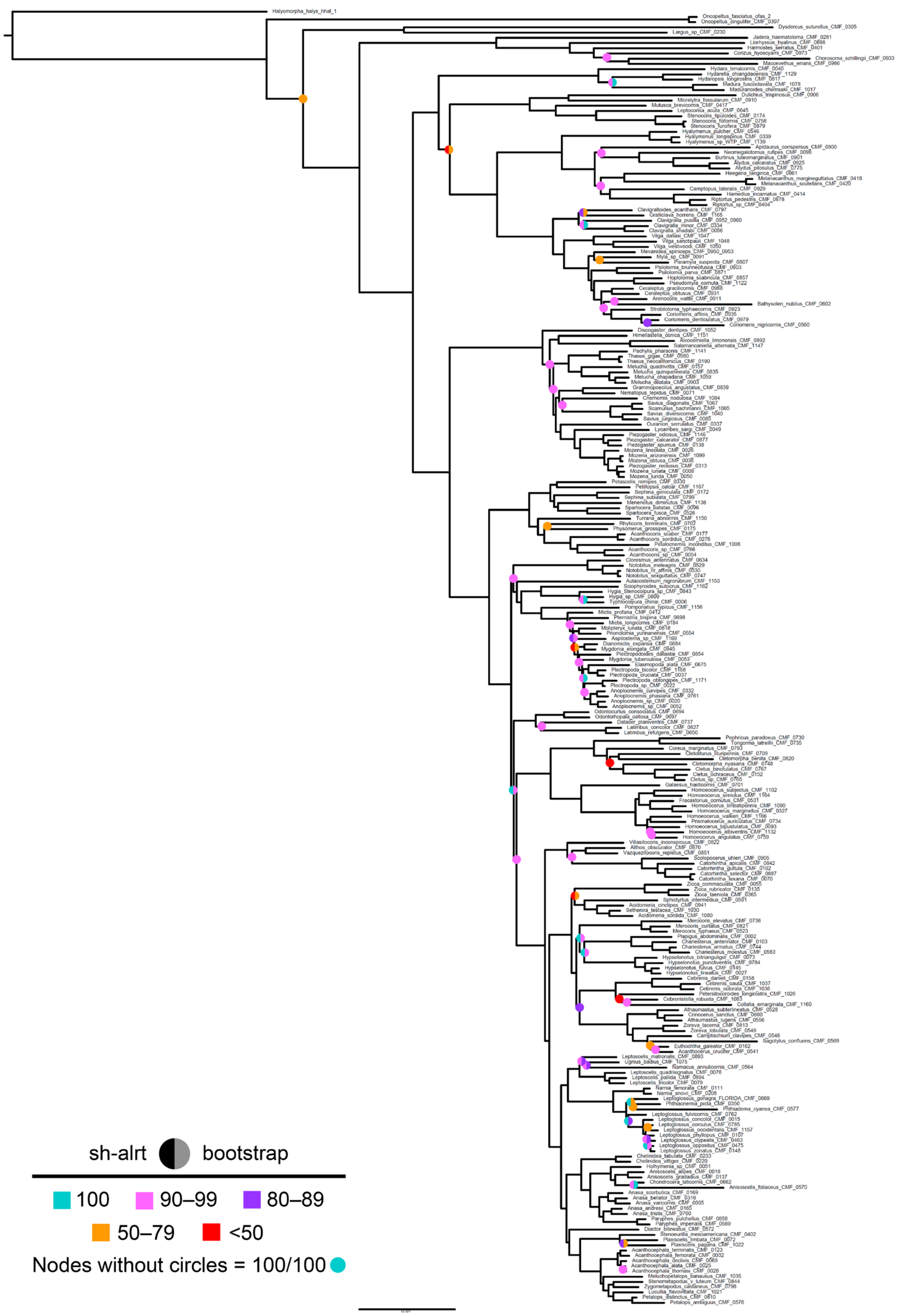


Figure S2. 70p ML best tree. Circles at nodes represent instances when sh-alrt and/or bootstrap support are less than 100.

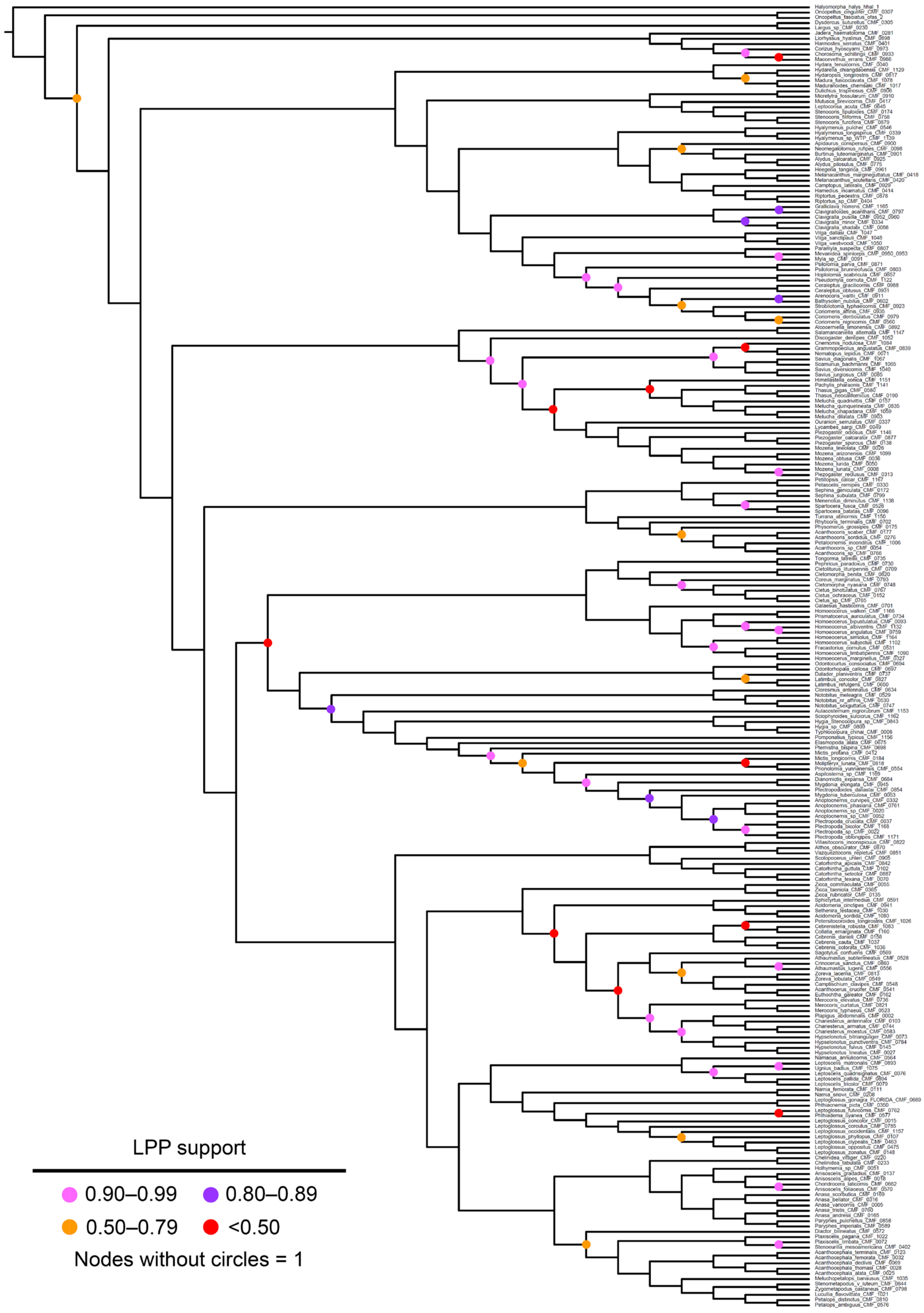


Figure S3. Multispecies coalescent (MSC) species tree based on 50p gene trees (displayed as a cladogram). Circles at nodes represent local posterior probabilities (LPPs) less than 1.

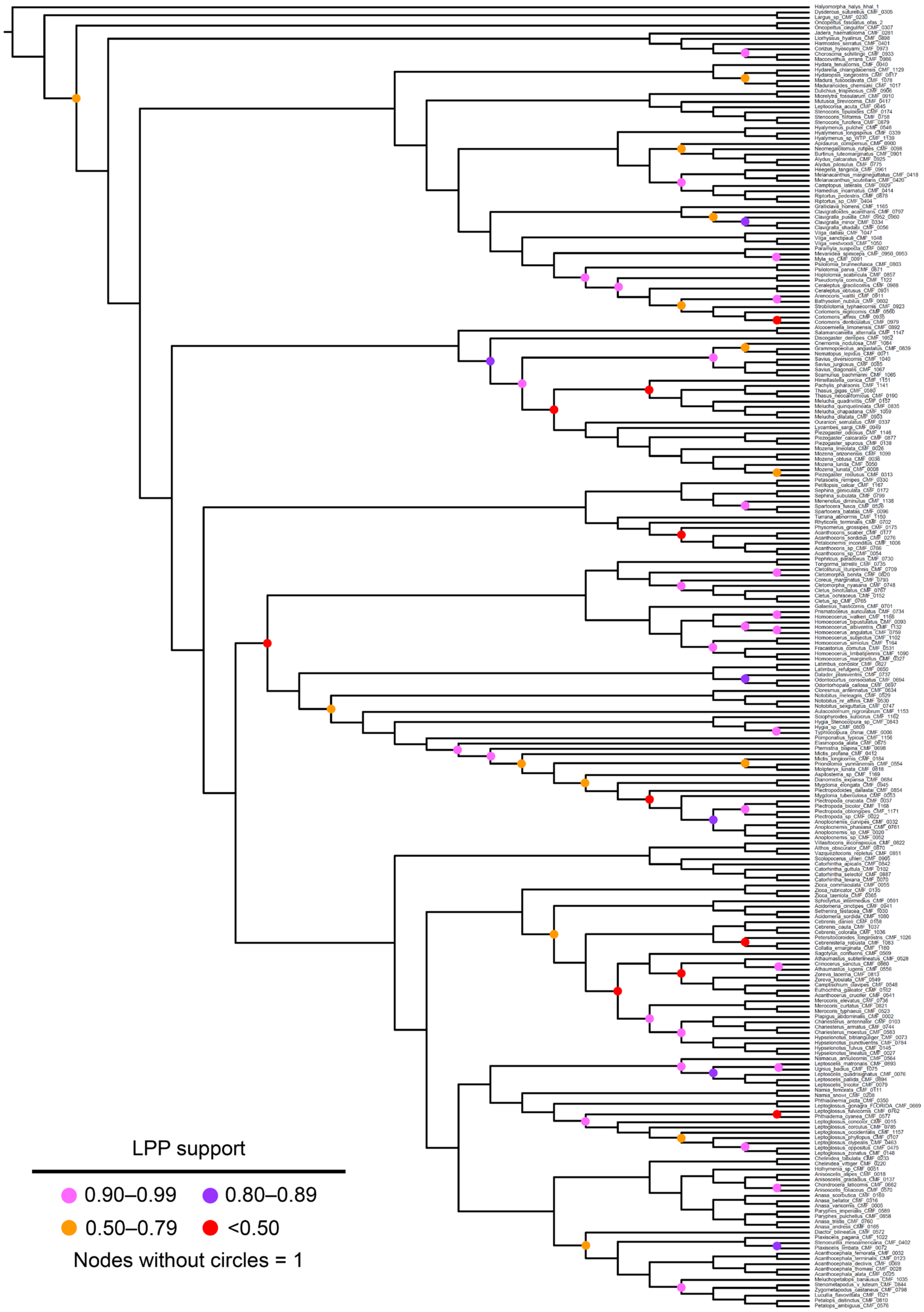


Figure S4. 70p MSC species tree (displayed as a cladogram). Circles at nodes represent LPPs less than 1.

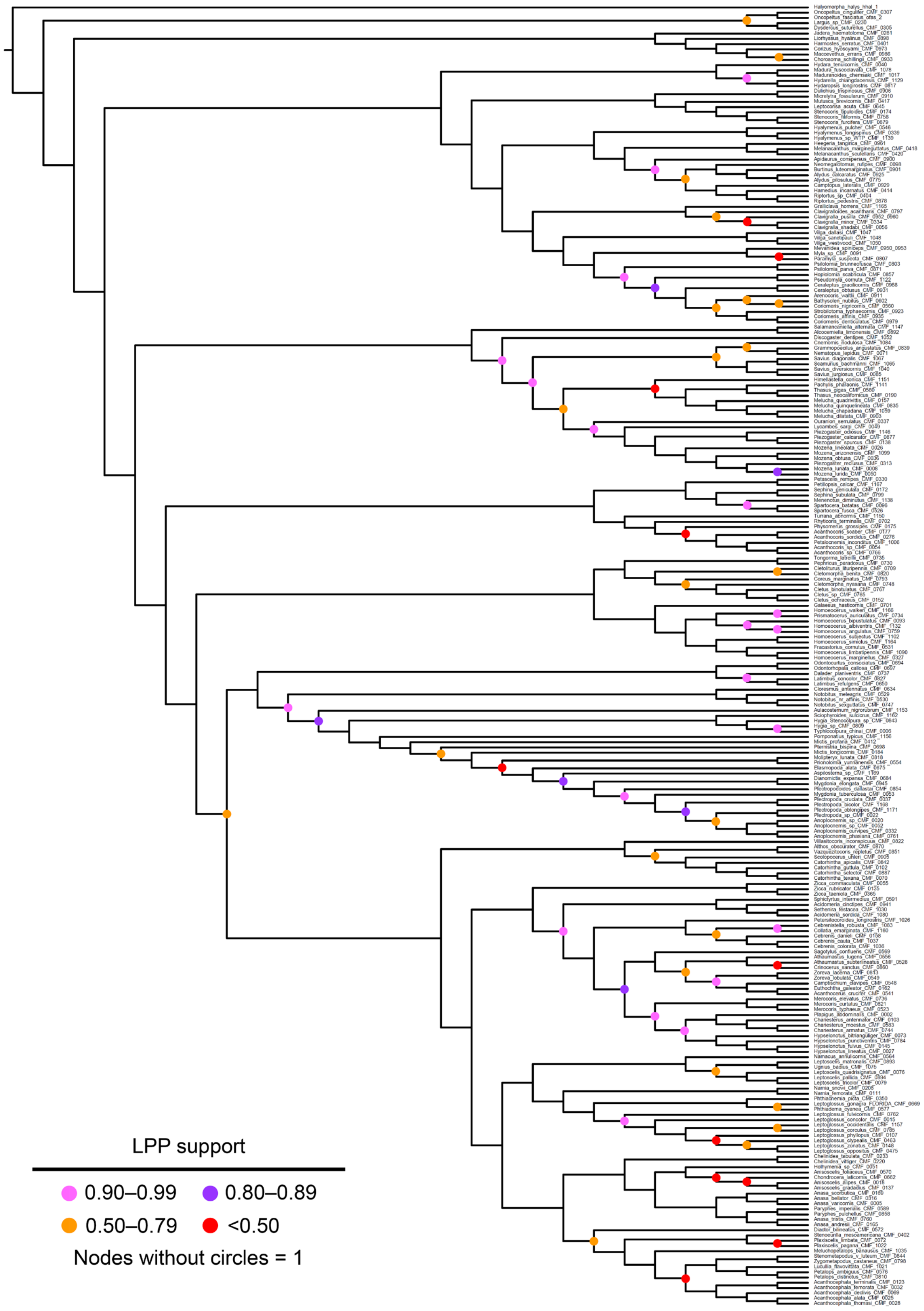


Figure S5. 50p25mi MSC species tree (displayed as a cladogram). Circles at nodes represent LPPs less than 1.

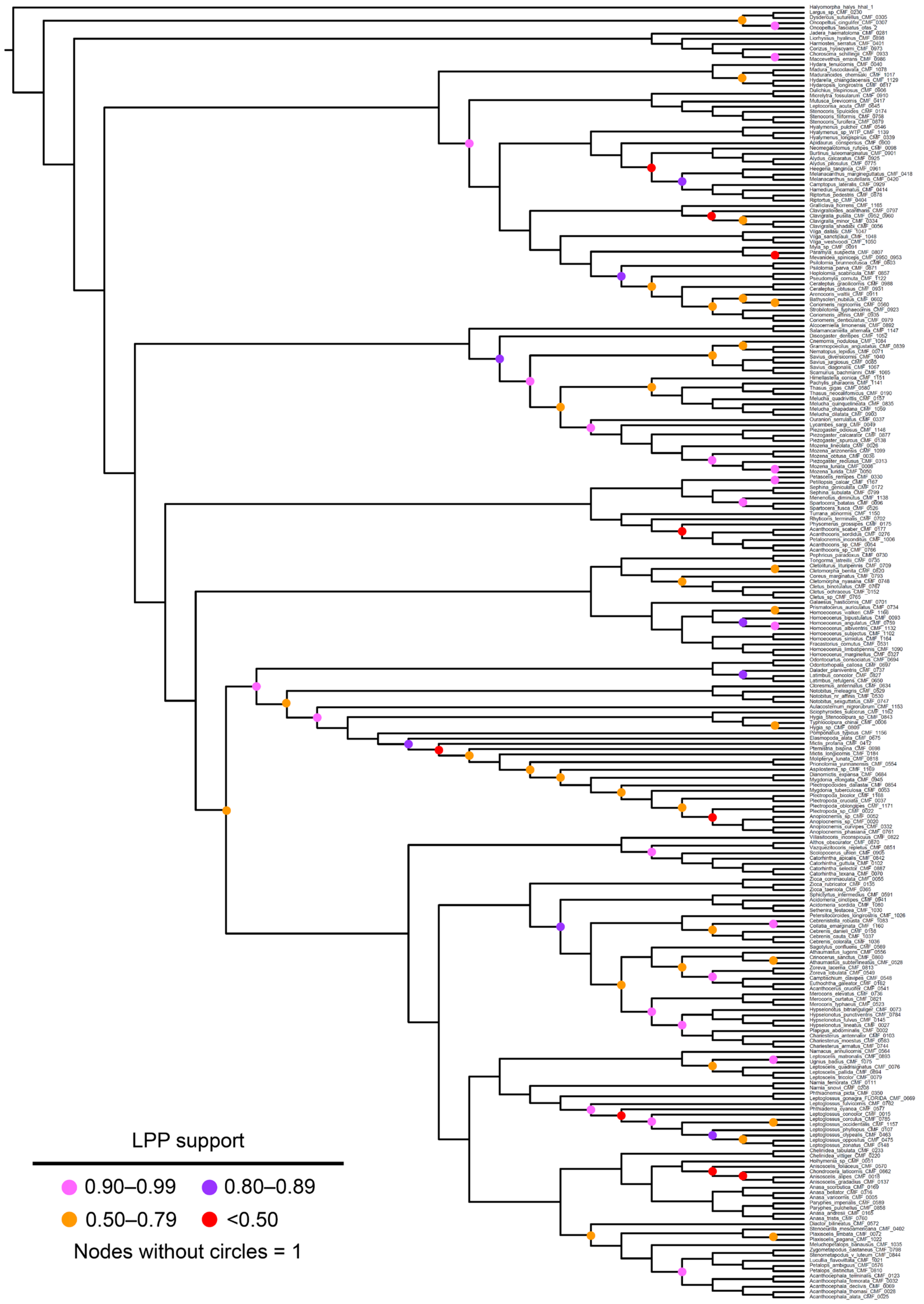


Figure S6. 70p25mi MSC species tree (displayed as a cladogram). Circles at nodes represent LPPs less than 1.

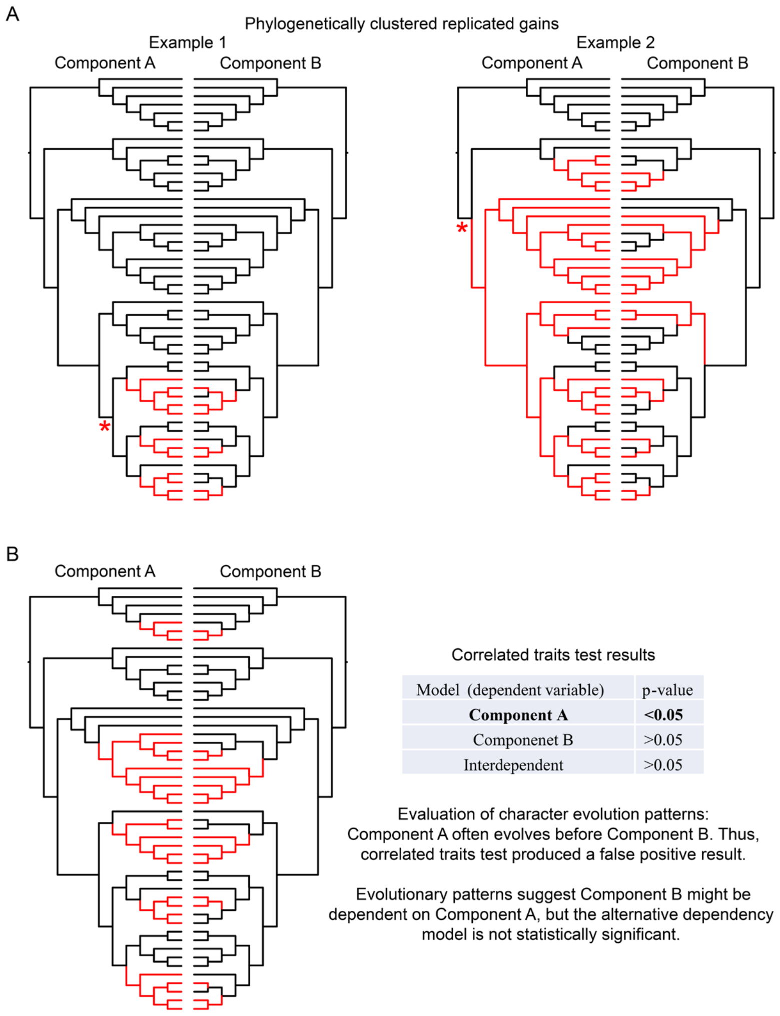


Figure S7. Evolutionary scenarios between hypothetical Components A and B on the same phylogeny. State 0 (“absent”) is shown in black lines, and State 1 (“present”) is shown in red. (A) Examples of Components A and B showing replicated evolutionary patterns in clades that are in relatively close proximity to one another on the phylogenetic tree. According to Maddison & FitzJohn (2015), the rate of origins in Component B could potentially be explained by a third, unrelated trait that changed in the larger clade marked by the red asterisk. (B) A scenario in which a correlated traits test of Components A and B finds a significant evolutionary model in which Component A depends on the evolutionary origins of Component B. However, when evaluating the evolutionary origins of Components A and B based on ancestral state estimates, Component A generally evolves first, with Component B evolving later in the same clades; this does not support prediction of the significant model. Thus, the evolutionary patterns might alternatively suggest Component B to be dependent on Component A, but this alternative model is not supported by the correlated traits test. As such, the correlated traits test is considered to have resulted in a false positive for the Component A dependency model, and the two components are not considered correlated in the absence of a statistically significant, alternative dependency model.

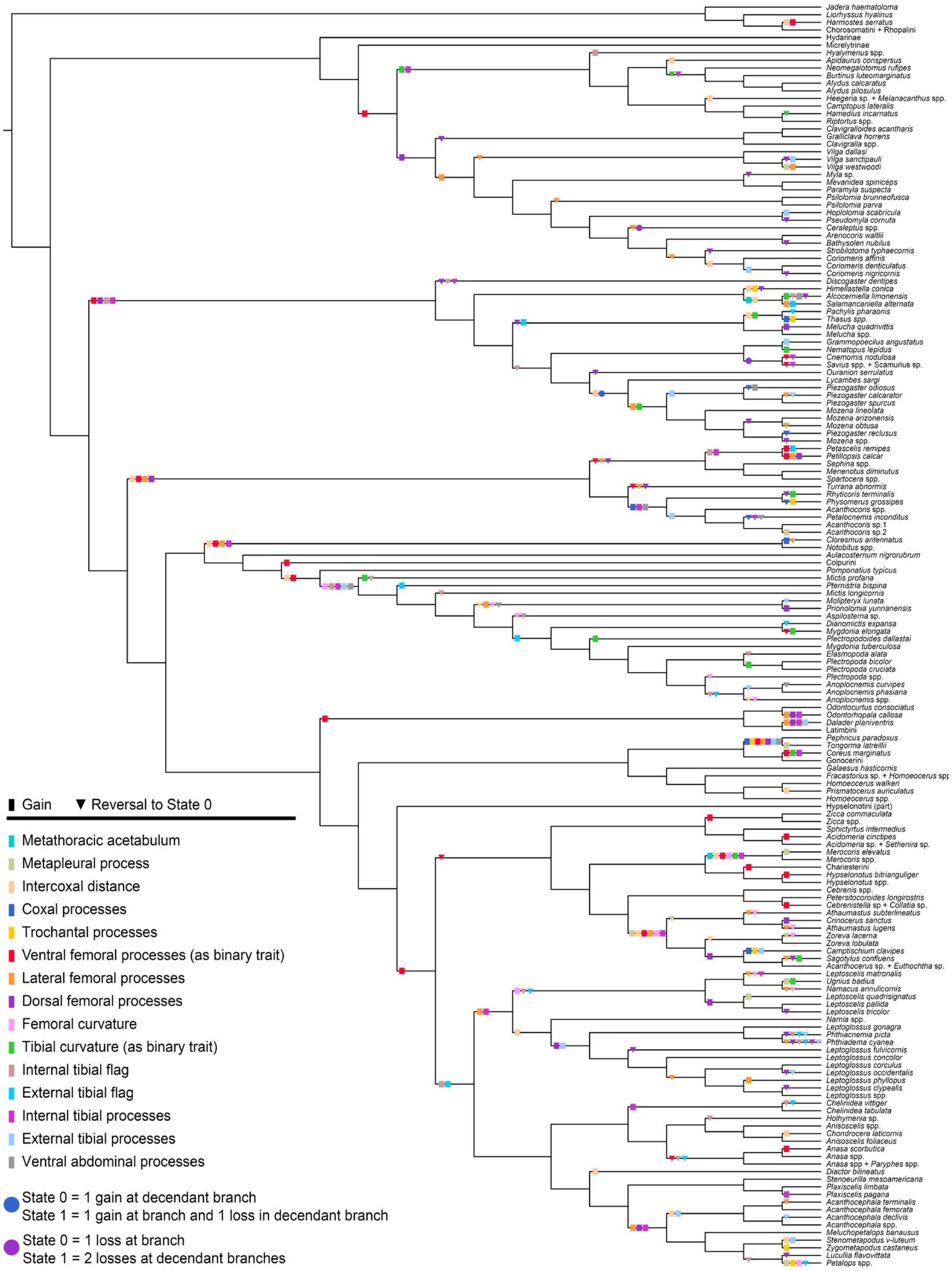


Fig. S8. Summary of gains and losses across all male hind leg components. The 50p ML ultrametric tree is displayed as a cladogram, and sister taxa with the same sets of component states have been collapsed into a single terminal for visualization. For simplicity in summarizing general component transitions, multistate Components #6 and #10 are shown as binary states.

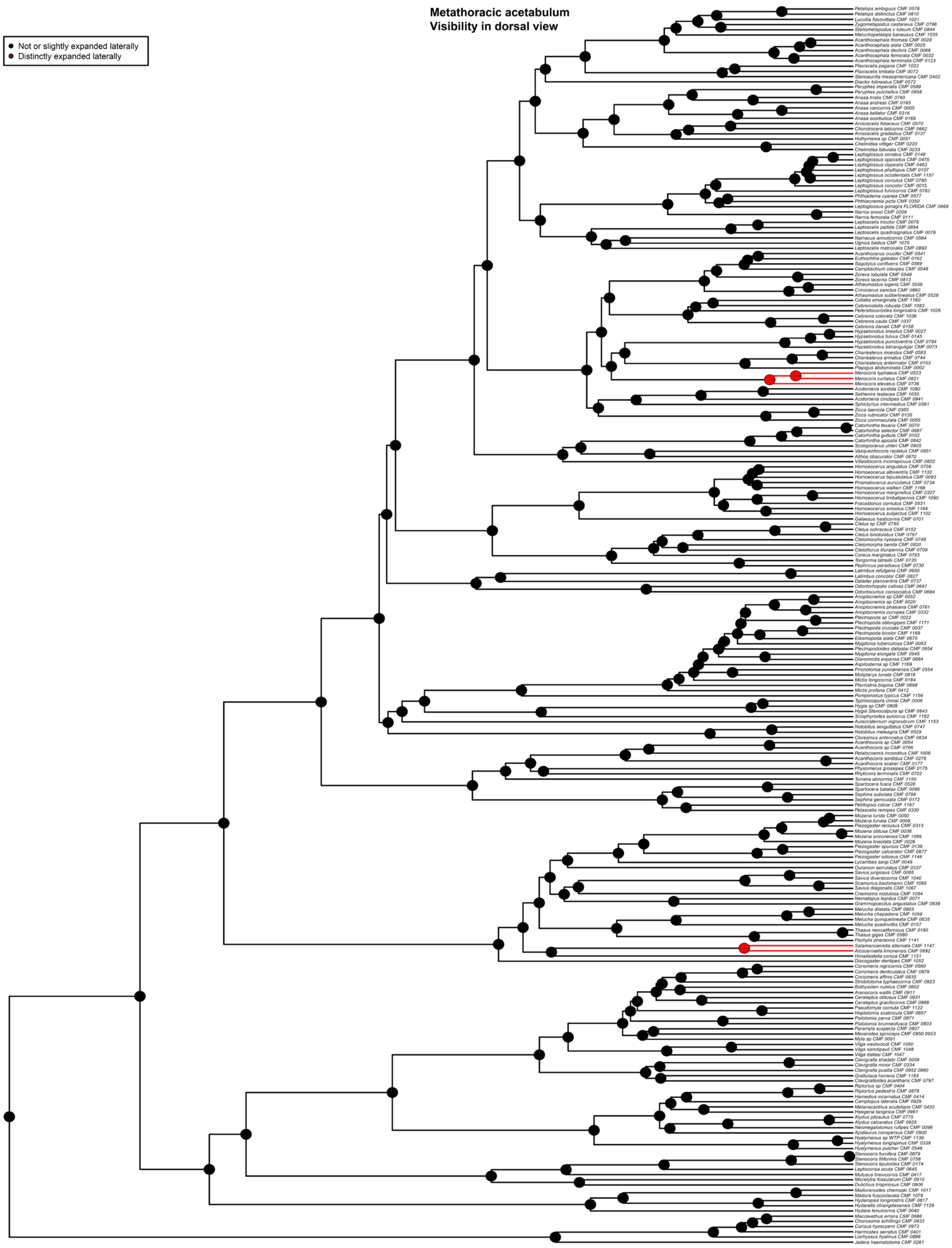


Fig. S9. Ancestral state estimates (ASE) based on the 50p ML ultrametric tree and equal rates (ER) model for Component #1. Taxa with missing data for Component #1 are pruned from the tree for analysis. Pie charts show the likeliest states for a given node (State 0 = black, State 1 = red), with branches similarly colored to represent the most likely state.

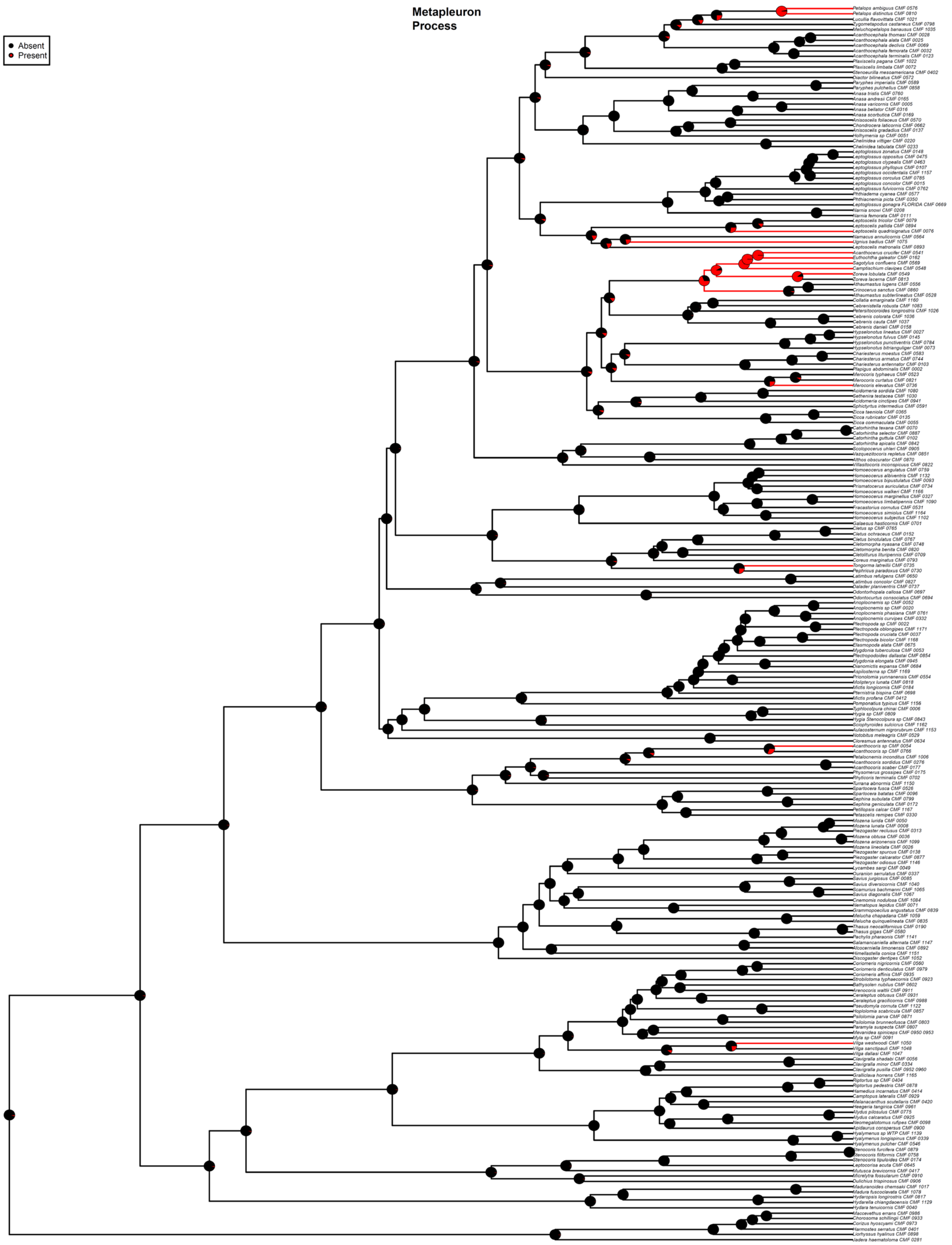


Fig. S10. ASE based on the 50p ML ultrametric tree and all rate different (ARD) model for Component #2. Taxa with missing data for Component #2 are pruned from the tree for analysis. Pie charts show the likeliest states for a given node (State 0 = black, State 1 = red), with branches similarly colored to represent the most likely state.

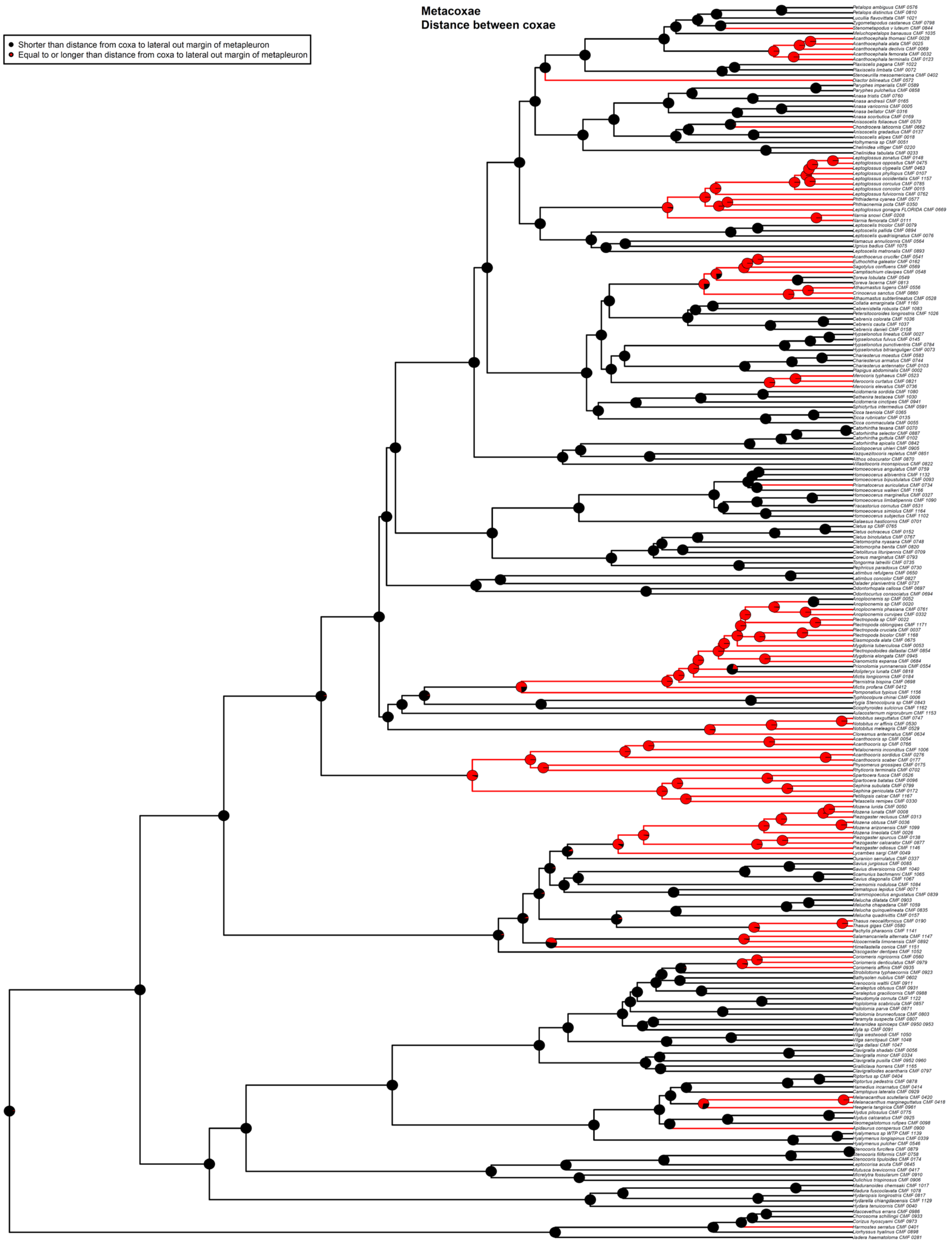


Fig. S11. ASE based on the 50p ML ultrametric tree and ER model for Component #3. Taxa with missing data for Component #3 are pruned from the tree for analysis. Pie charts show the likeliest states for a given node (State 0 = black, State 1 = red), with branches similarly colored to represent the most likely state.

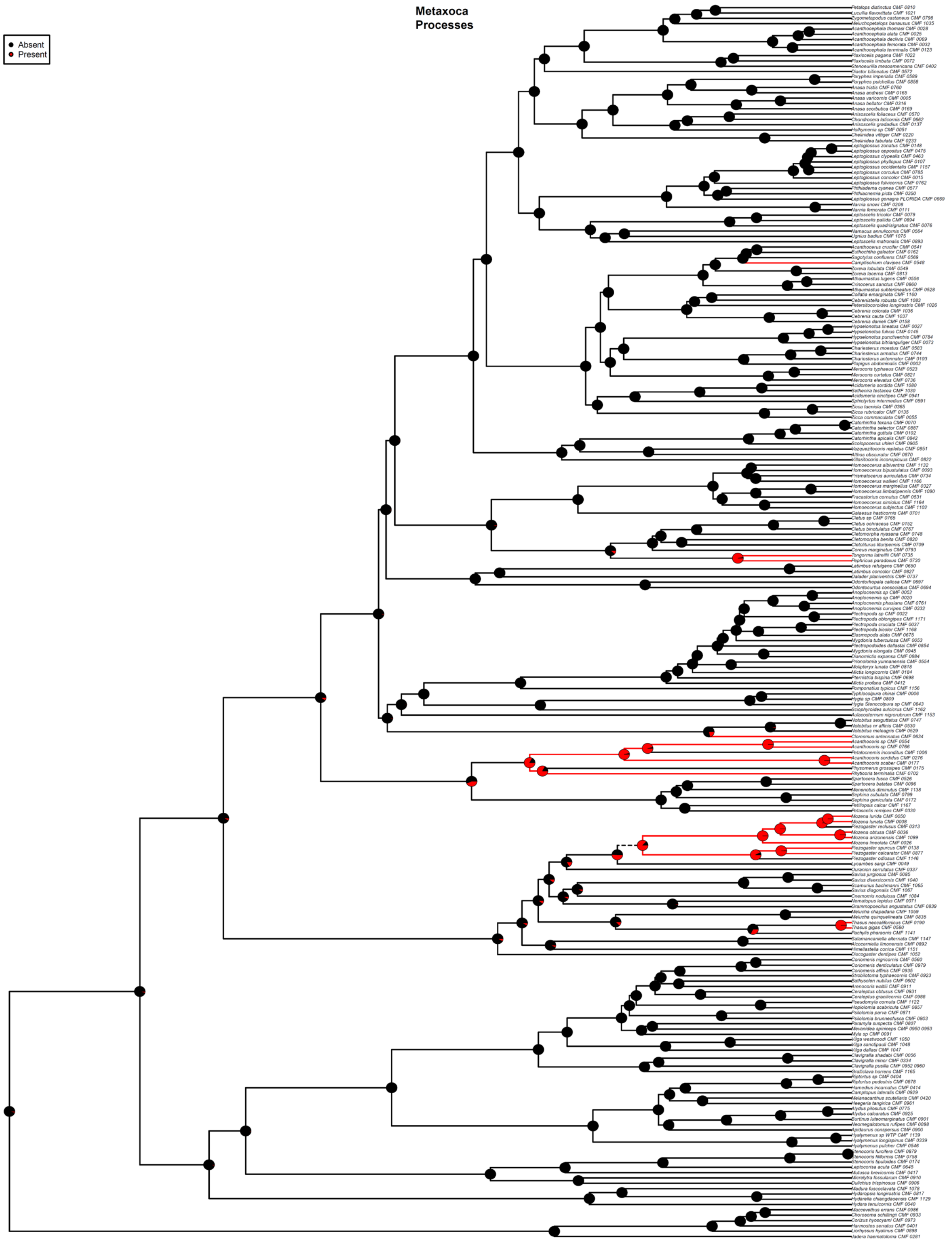


Fig. S12. ASE based on the 50p ML ultrametric tree and ARD model for Component #4. Taxa with missing data for Component #4 are pruned from the tree for analysis. Pie charts show the likeliest states for a given node (State 0 = black, State 1 = red), with branches similarly colored to represent the most likely state.

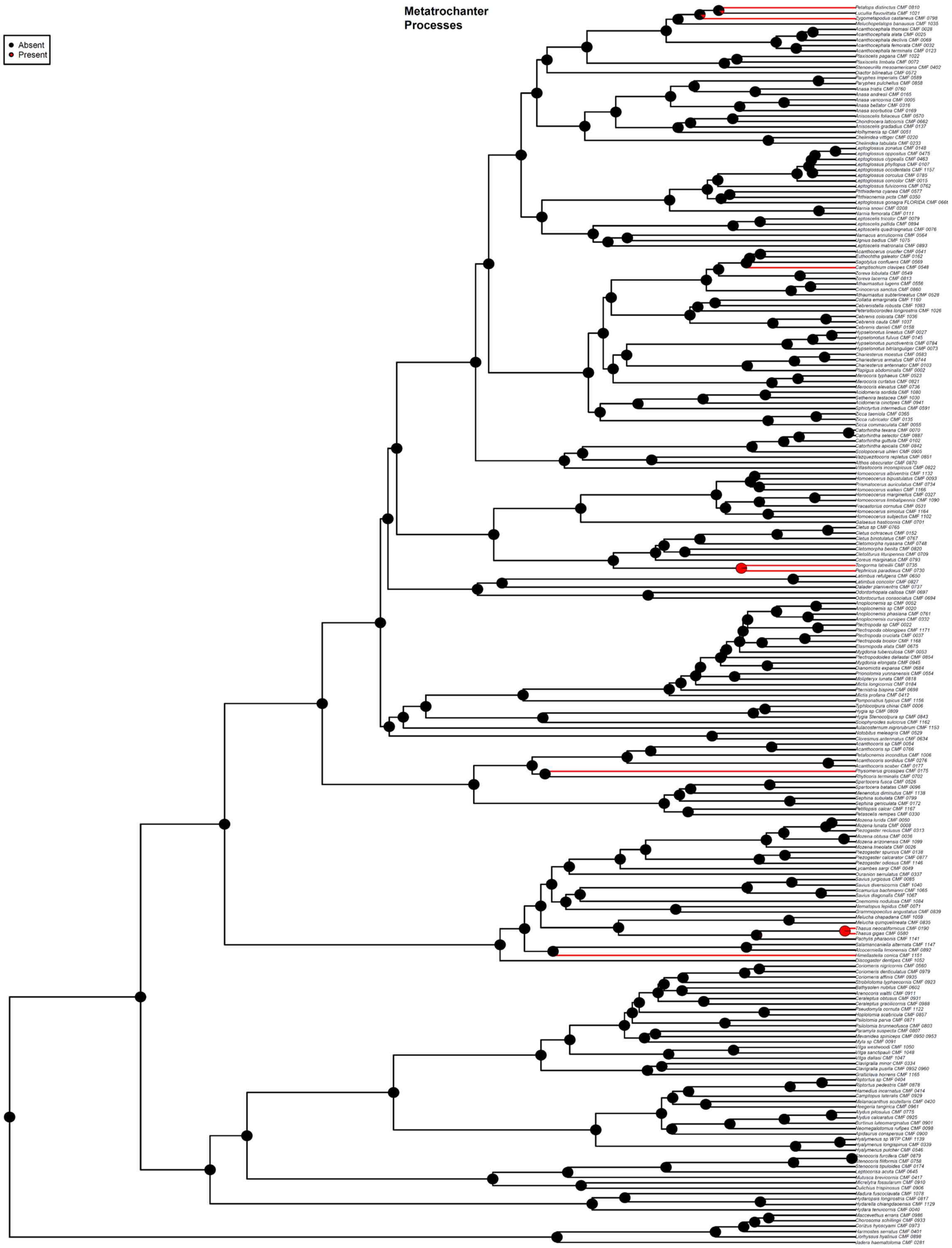


Fig. S13. ASE based on the 50p ML ultrametric tree and ER model for Component #5. Taxa with missing data for Component #5 are pruned from the tree for analysis. Pie charts show the likeliest states for a given node (State 0 = black, State 1 = red), with branches similarly colored to represent the most likely state.

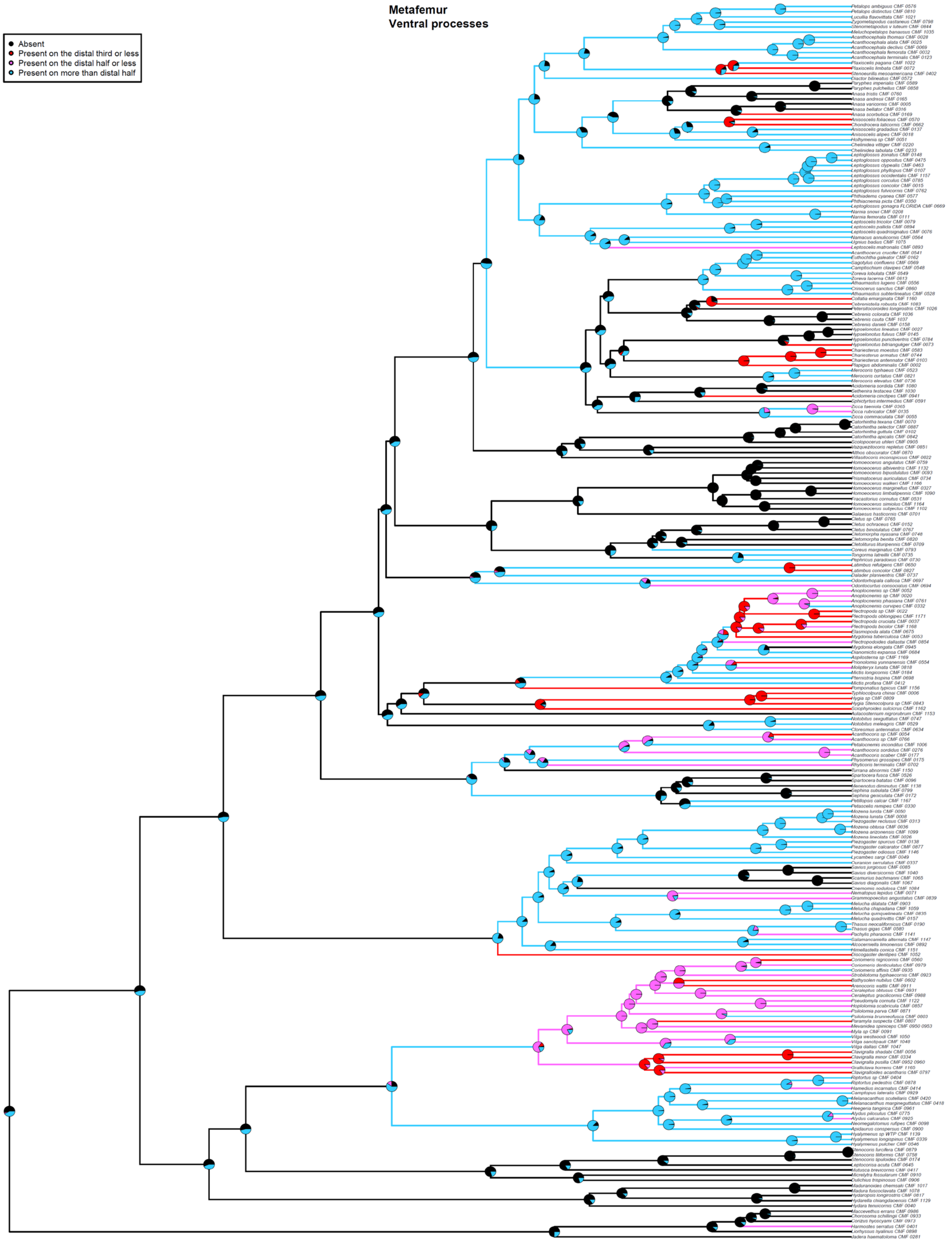


Fig. S14. ASE based on the 50p ML ultrametric tree and symmetric rates (SYM) model for Component #6. Taxa with missing data for Component #6 are pruned from the tree for analysis. Pie charts show the likeliest states for a given node (State 0 = black, State 1 = red, State 2 = magenta, State 3 = blue), with branches similarly colored to represent the most likely state.

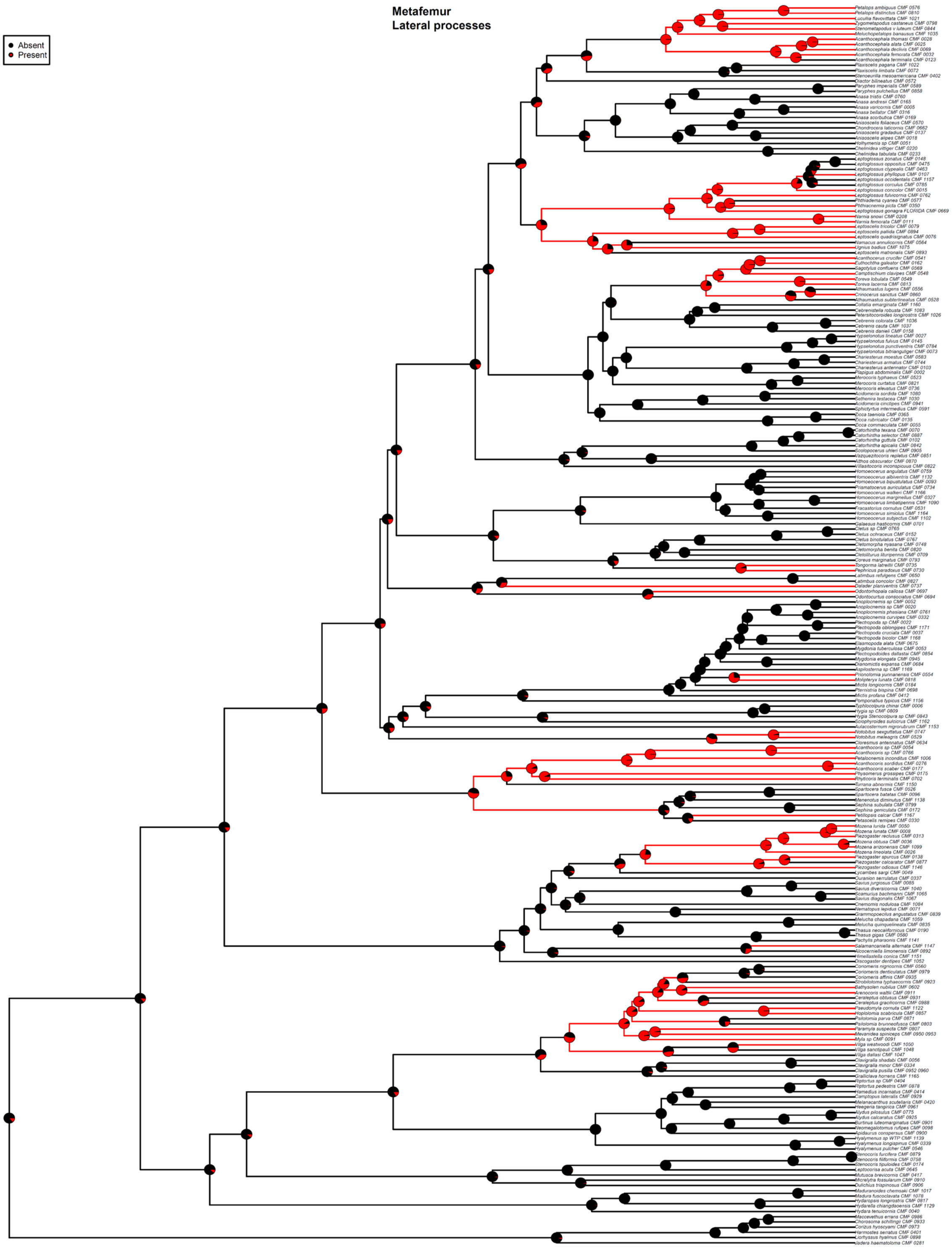


Fig. S15. ASE based on the 50p ML ultrametric tree and ARD model for Component #7. Taxa with missing data for Component #7 are pruned from the tree for analysis. Pie charts show the likeliest states for a given node (State 0 = black, State 1 = red), with branches similarly colored to represent the most likely state.

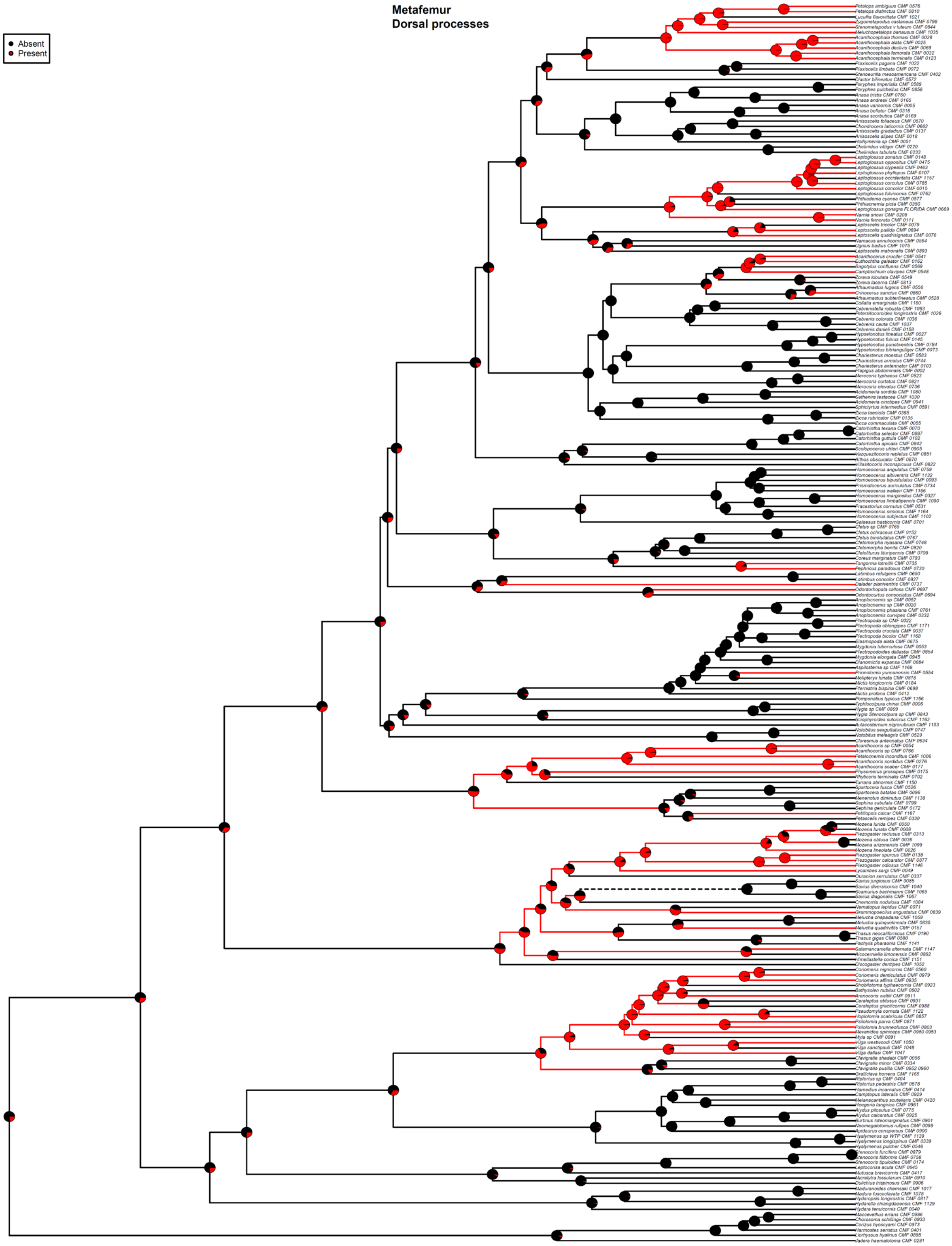


Fig. S16. ASE based on the 50p ML ultrametric tree and ARD model for Component #8. Taxa with missing data for Component #8 are pruned from the tree for analysis. Pie charts show the likeliest states for a given node (State 0 = black, State 1 = red), with branches similarly colored to represent the most likely state.

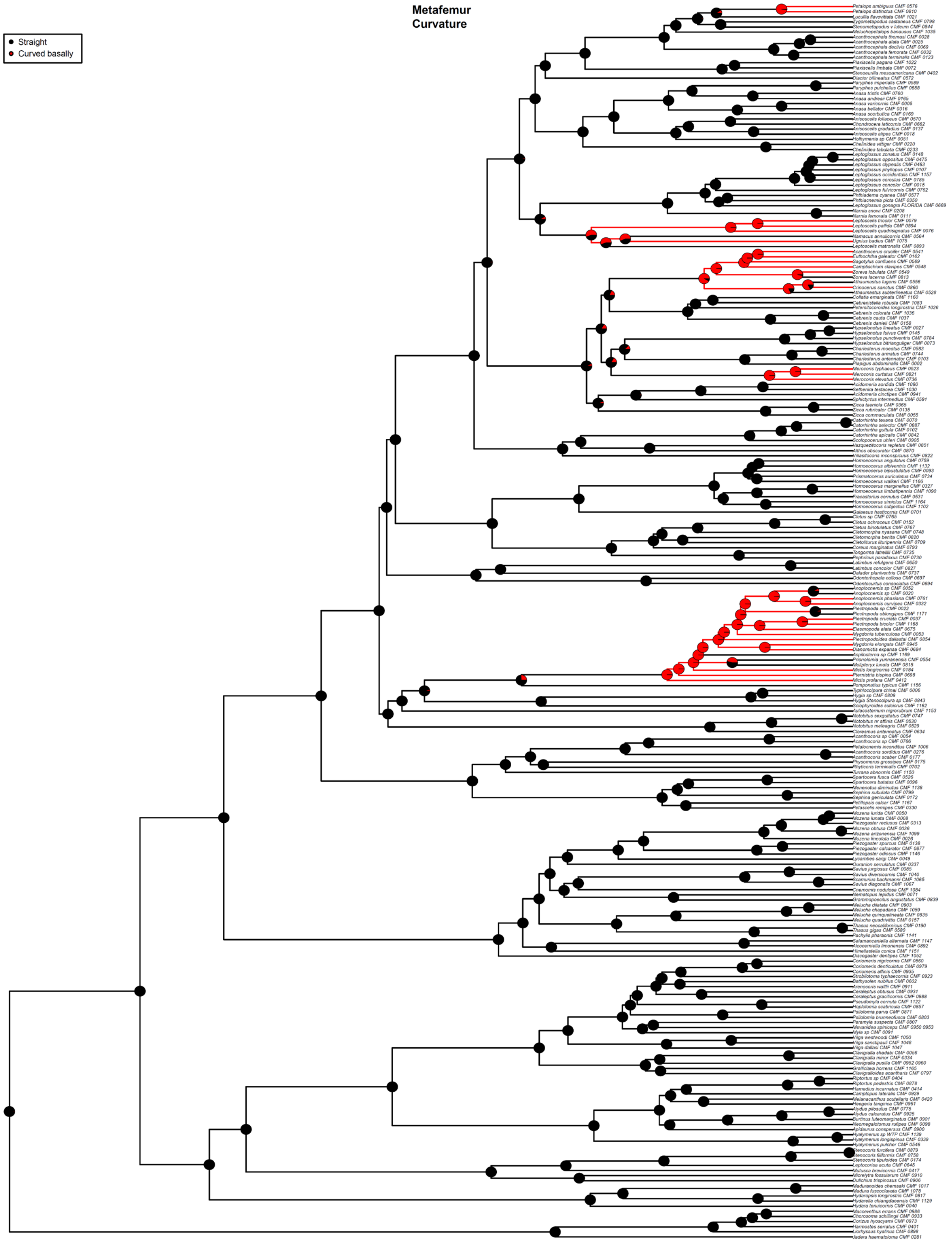


Fig. S17. ASE based on the 50p ML ultrametric tree and ARD model for Component #9. Taxa with missing data for Component #9 are pruned from the tree for analysis. Pie charts show the likeliest states for a given node (State 0 = black, State 1 = red), with branches similarly colored to represent the most likely state.

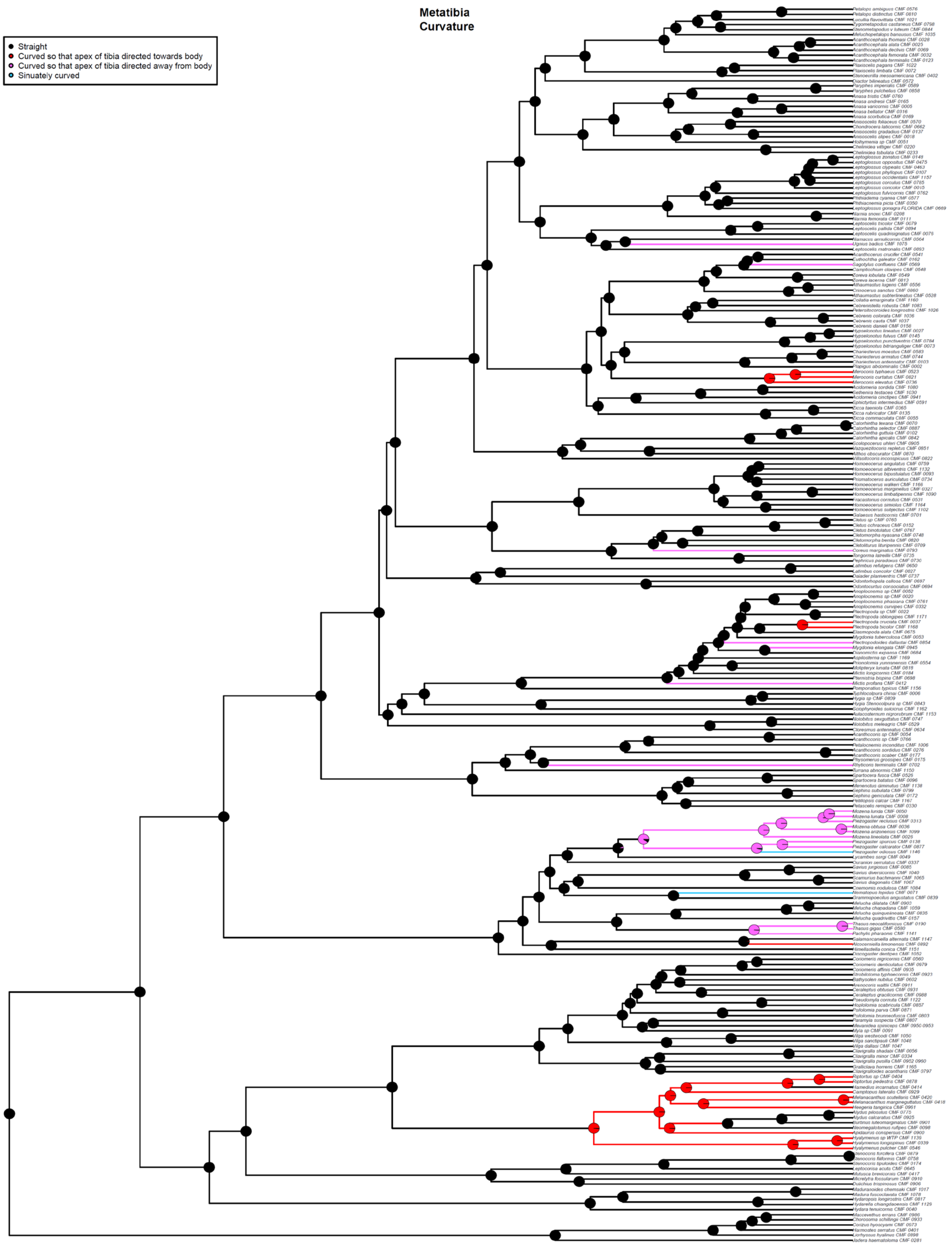


Fig. S18. ASE based on the 50p ML ultrametric tree and ER model for Component #10. Taxa with missing data for Component #10 are pruned from the tree for analysis. Pie charts show the likeliest states for a given node (State 0 = black, State 1 = red, State 2 = magenta, State 3 = blue), with branches similarly colored to represent the most likely state.

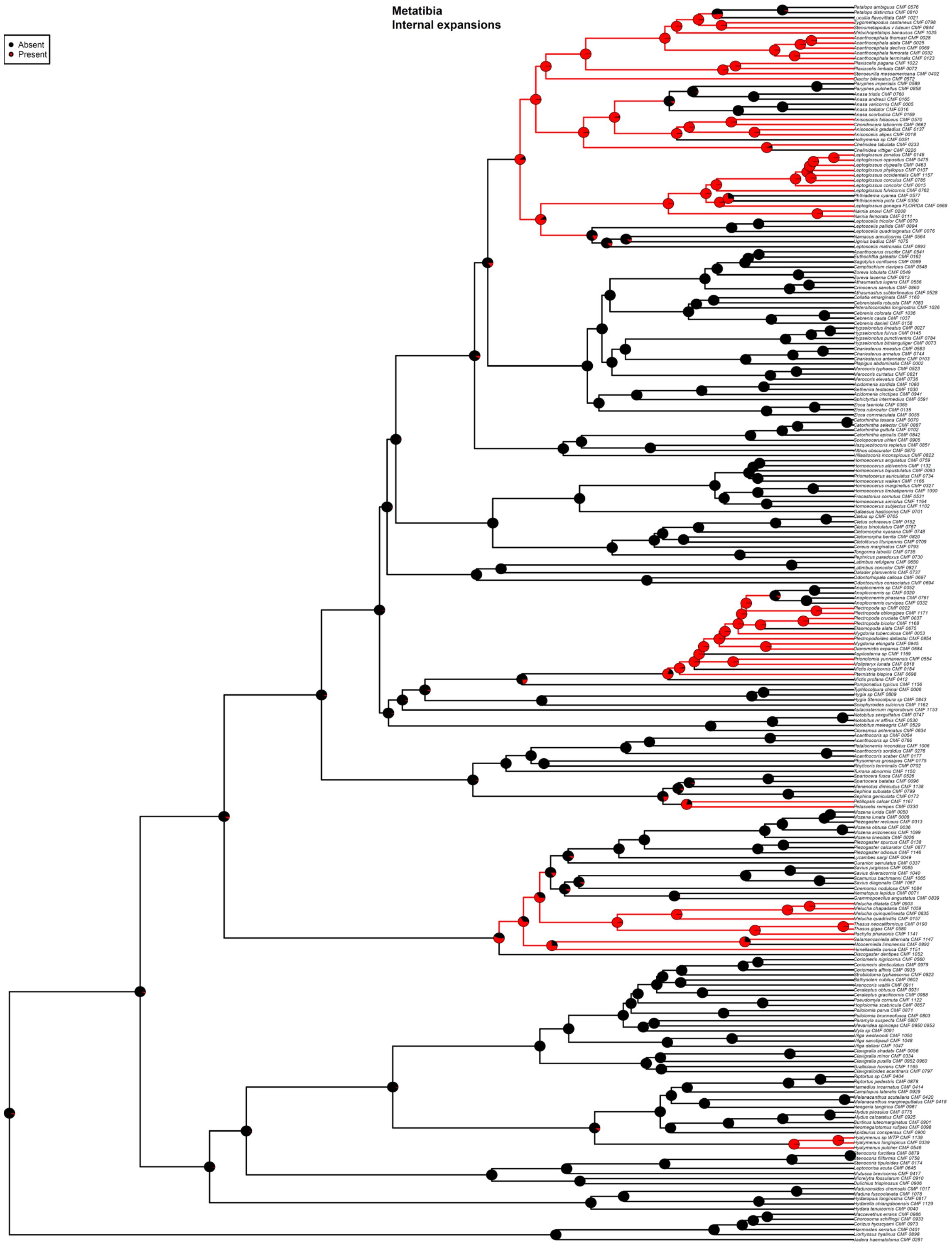


Fig. S19. ASE based on the 50p ML ultrametric tree and ARD model for Component #11. Taxa with missing data for Component #11 are pruned from the tree for analysis. Pie charts show the likeliest states for a given node (State 0 = black, State 1 = red), with branches similarly colored to represent the most likely state.

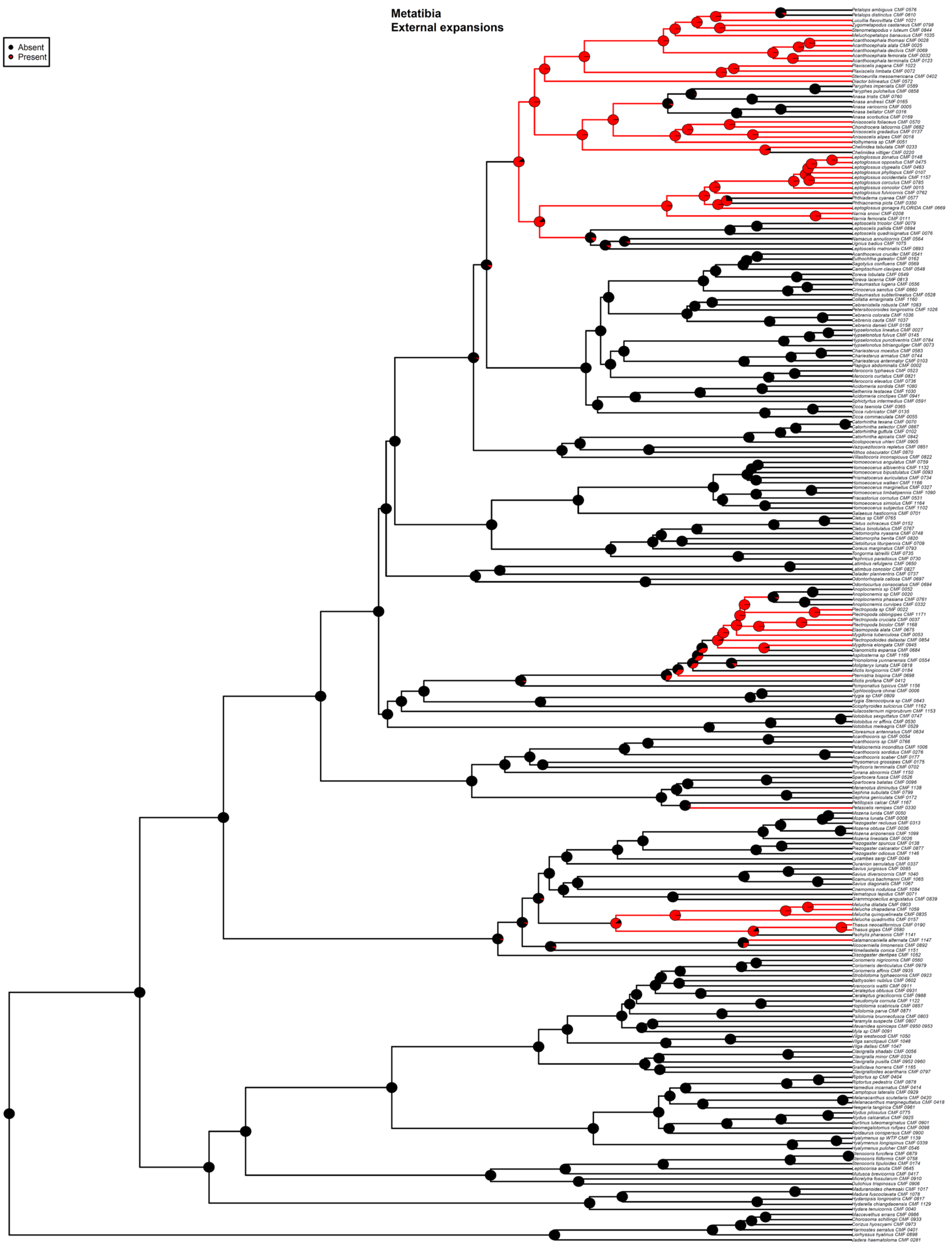


Fig. S20. ASE based on the 50p ML ultrametric tree and ARD model for Component #12. Taxa with missing data for Component #12 are pruned from the tree for analysis. Pie charts show the likeliest states for a given node (State 0 = black, State 1 = red), with branches similarly colored to represent the most likely state.

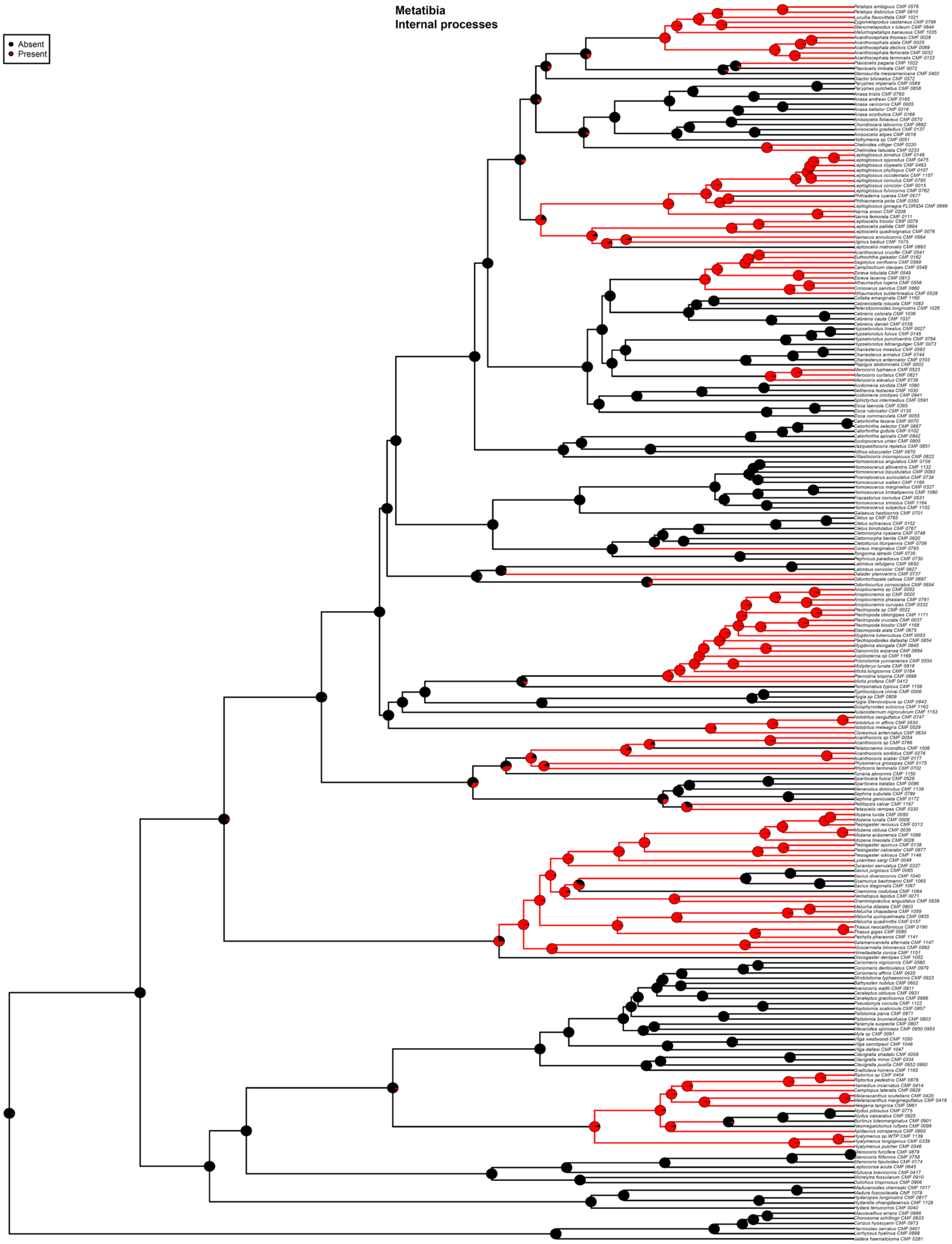


Fig. S21. ASE based on the 50p ML ultrametric tree and ER model for Component #13. Taxa with missing data for Component #13 are pruned from the tree for analysis. Pie charts show the likeliest states for a given node (State 0 = black, State 1 = red), with branches similarly colored to represent the most likely state.

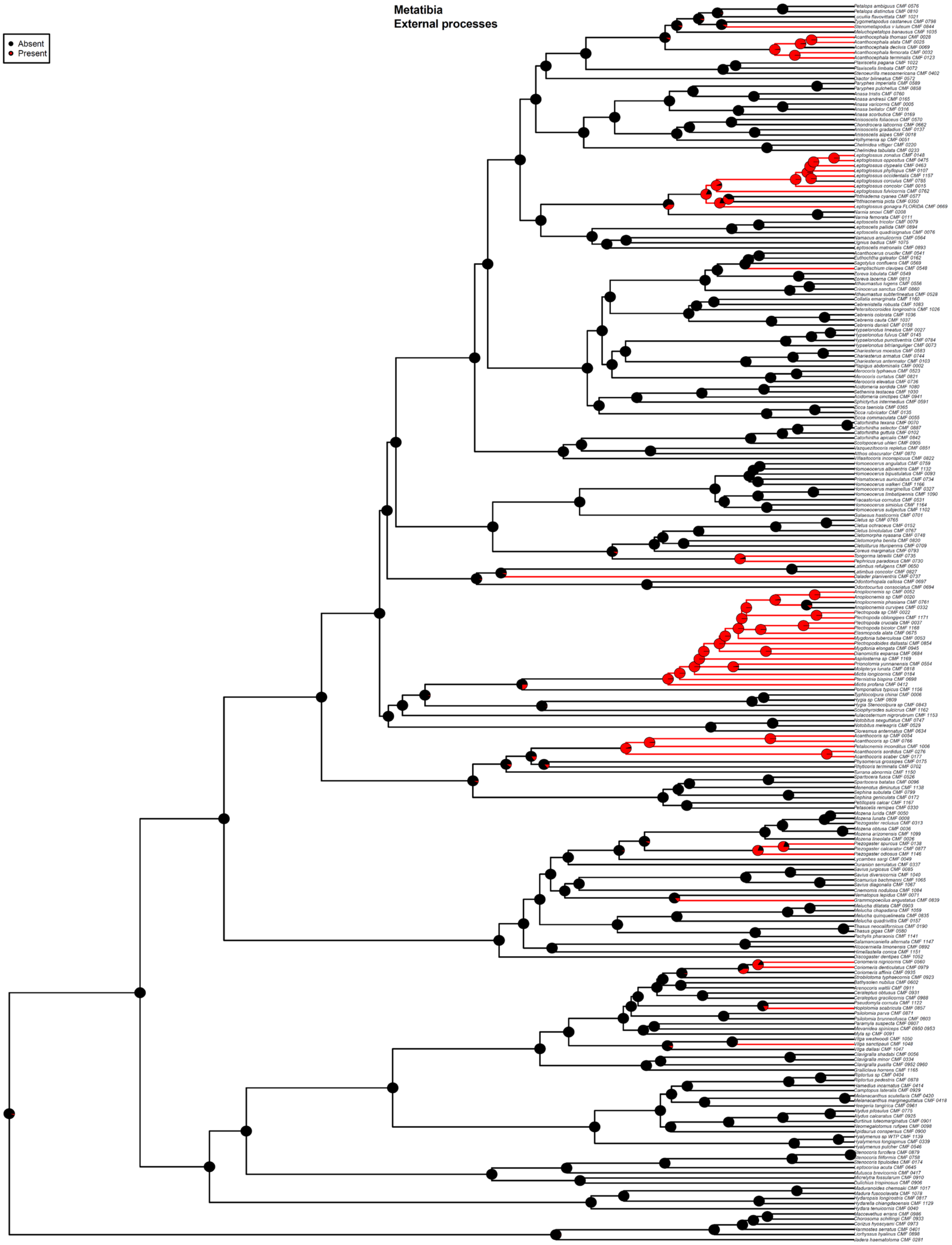


Fig. S22. ASE based on the 50p ML ultrametric tree and ARD model for Component #14. Taxa with missing data for Component #14 are pruned from the tree for analysis. Pie charts show the likeliest states for a given node (State 0 = black, State 1 = red), with branches similarly colored to represent the most likely state.

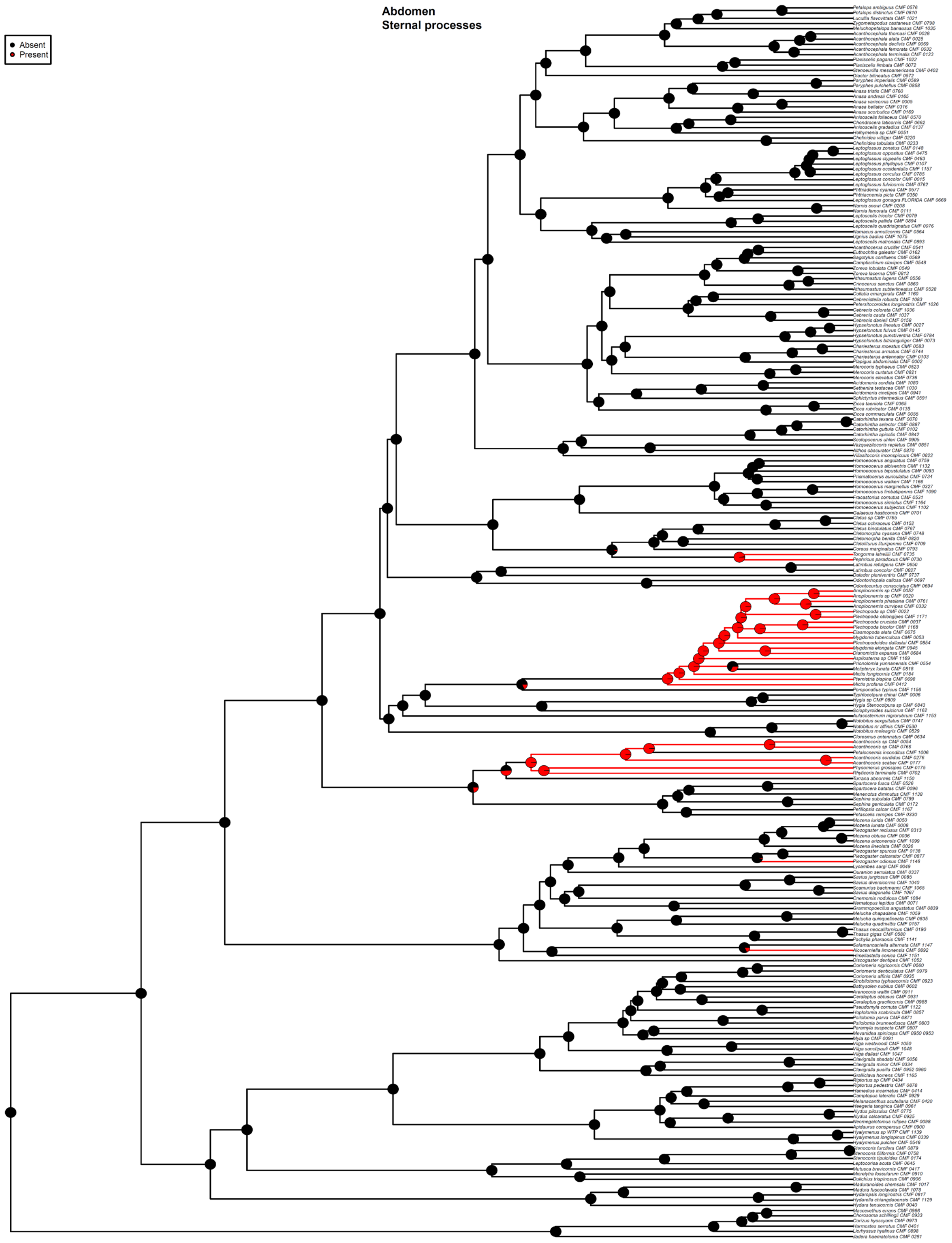


Fig. S23. ASE based on the 50p ML ultrametric tree and ARD model for Component #15. Taxa with missing data for Component #15 are pruned from the tree for analysis. Pie charts show the likeliest states for a given node (State 0 = black, State 1 = red), with branches similarly colored to represent the most likely state.

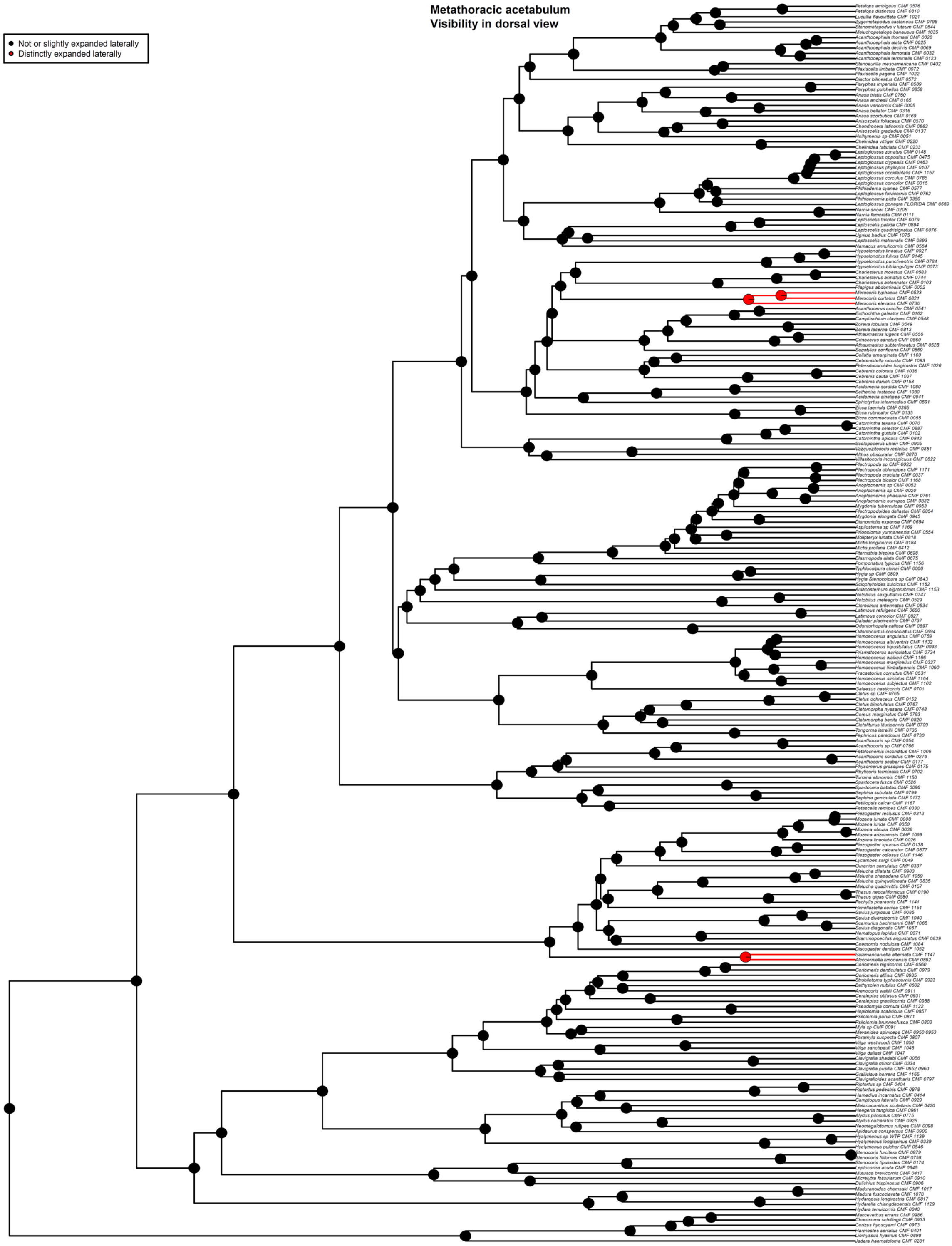


Fig. S24. ASE based on the 50p MSC ultrametric tree and ER model for Component #1. Taxa with missing data for Component #1 are pruned from the tree for analysis. Pie charts show the likeliest states for a given node (State 0 = black, State 1 = red), with branches similarly colored to represent the most likely state.

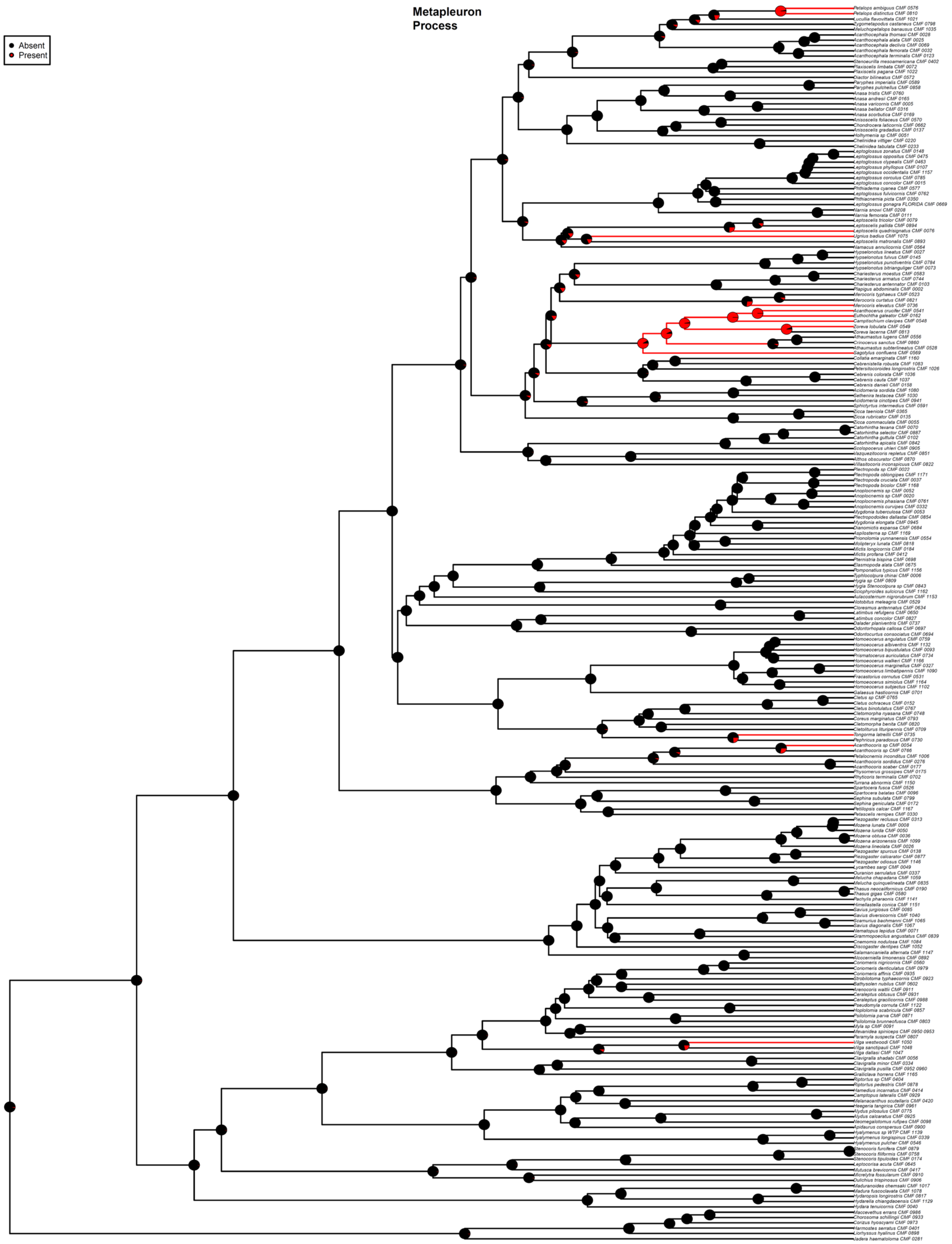


Fig. S25. ASE based on the 50p MSC ultrametric tree and ARD model for Component #2. Taxa with missing data for Component #2 are pruned from the tree for analysis. Pie charts show the likeliest states for a given node (State 0 = black, State 1 = red), with branches similarly colored to represent the most likely state.

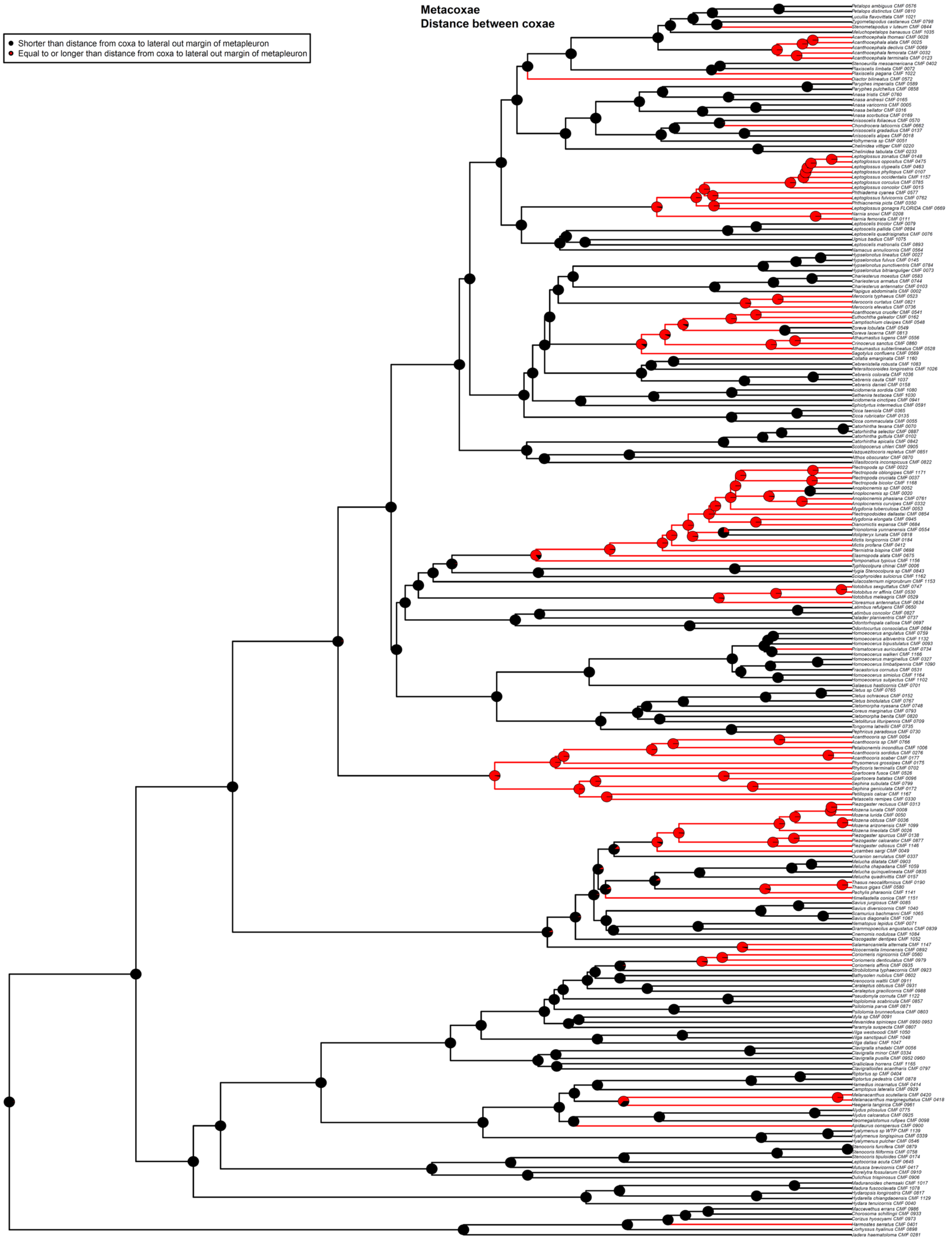


Fig. S26. ASE based on the 50p MSC ultrametric tree and ER model for Component #3. Taxa with missing data for Component #3 are pruned from the tree for analysis. Pie charts show the likeliest states for a given node (State 0 = black, State 1 = red), with branches similarly colored to represent the most likely state.

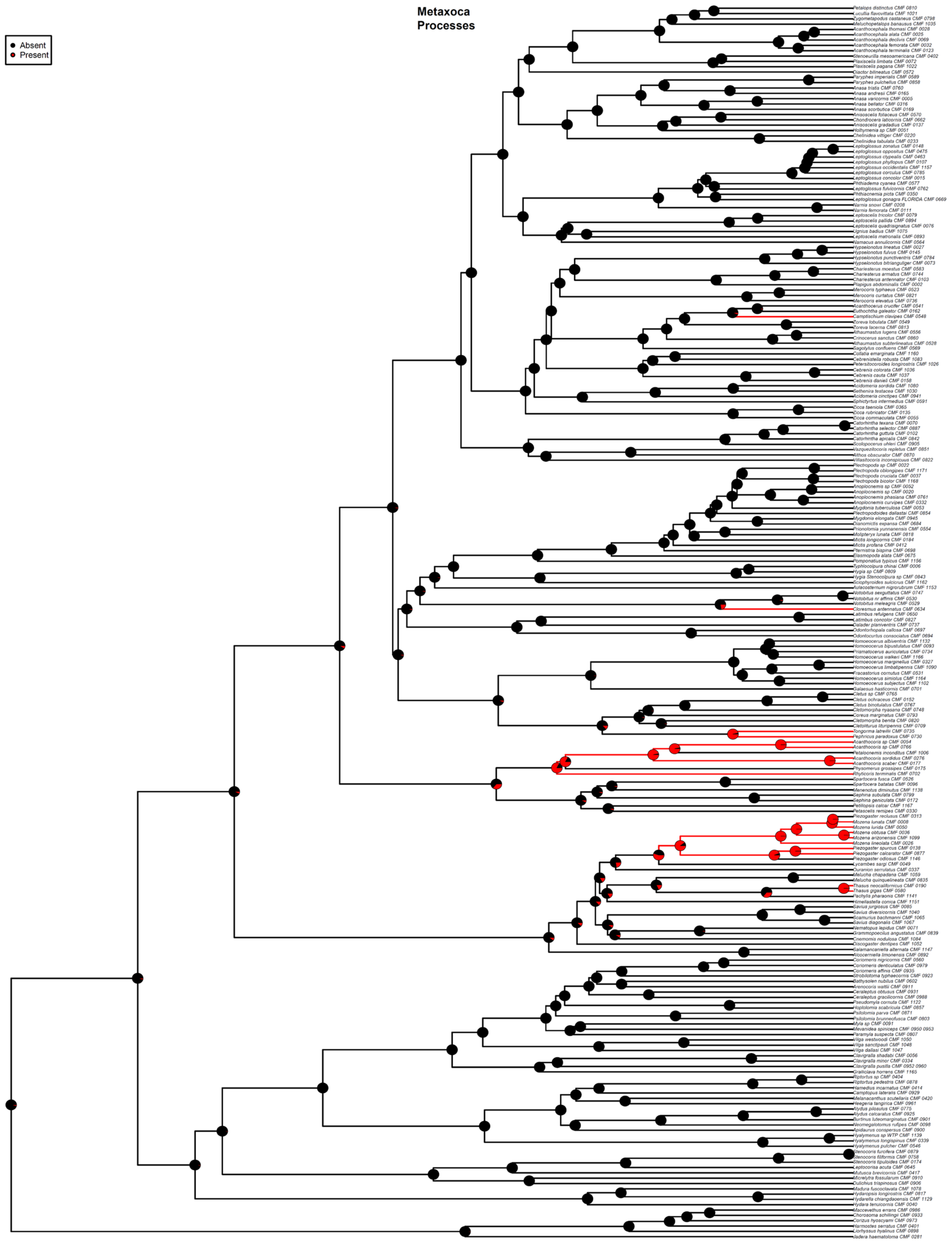


Fig. S27. ASE based on the 50p MSC ultrametric tree and ARD model for Component #4. Taxa with missing data for Component #4 are pruned from the tree for analysis. Pie charts show the likeliest states for a given node (State 0 = black, State 1 = red), with branches similarly colored to represent the most likely state.

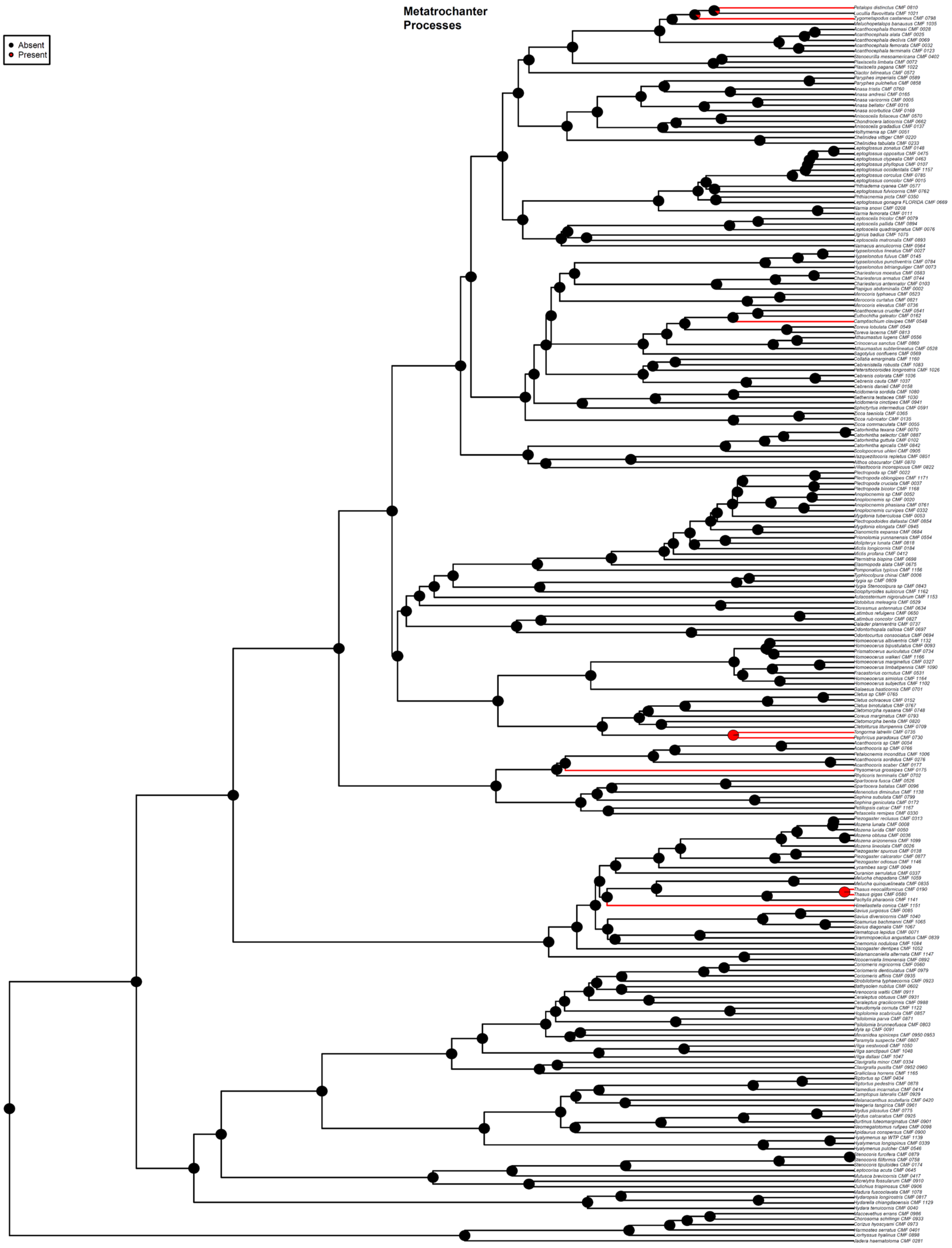


Fig. S28. ASE based on the 50p MSC ultrametric tree and ER model for Component #5. Taxa with missing data for Component #5 are pruned from the tree for analysis. Pie charts show the likeliest states for a given node (State 0 = black, State 1 = red), with branches similarly colored to represent the most likely state.

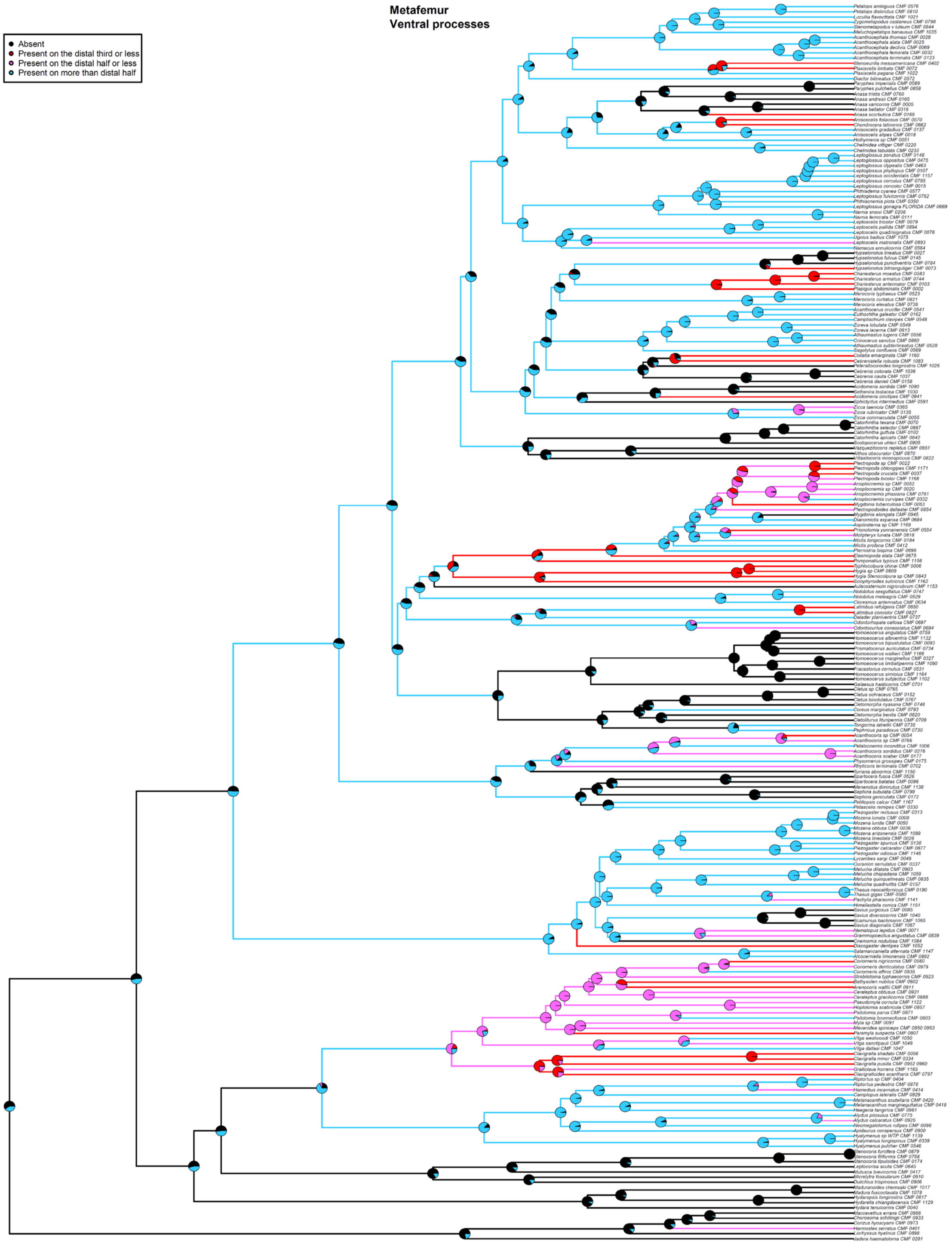


Fig. S29. ASE based on the 50p MSC ultrametric tree and SYM model for Component #6. Taxa with missing data for Component #6 are pruned from the tree for analysis. Pie charts show the likeliest states for a given node (State 0 = black, State 1 = red, State 2 = magenta, State 3 = blue), with branches similarly colored to represent the most likely state.

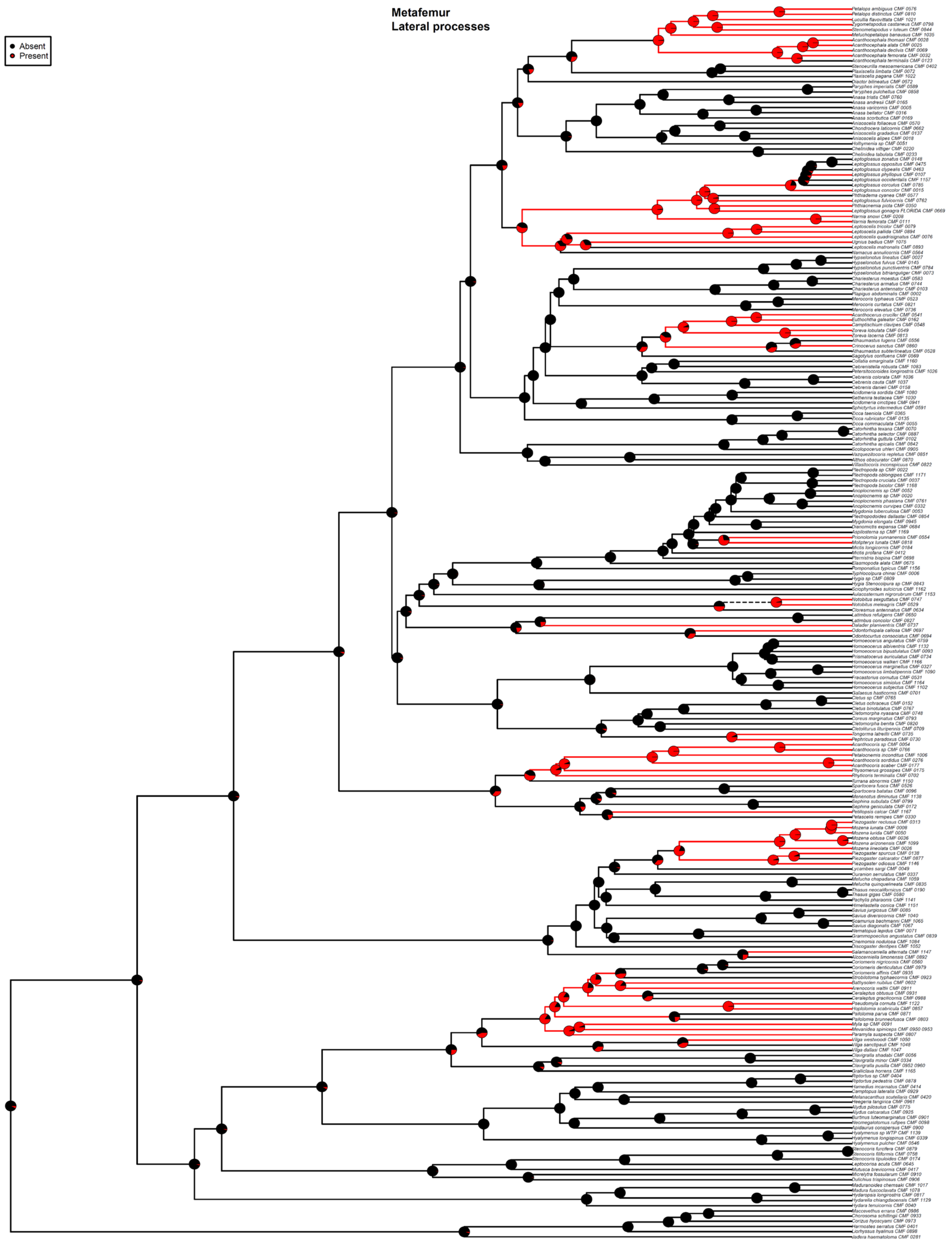


Fig. S30. ASE based on the 50p MSC ultrametric tree and ARD model for Component #7. Taxa with missing data for Component #7 are pruned from the tree for analysis. Pie charts show the likeliest states for a given node (State 0 = black, State 1 = red), with branches similarly colored to represent the most likely state.

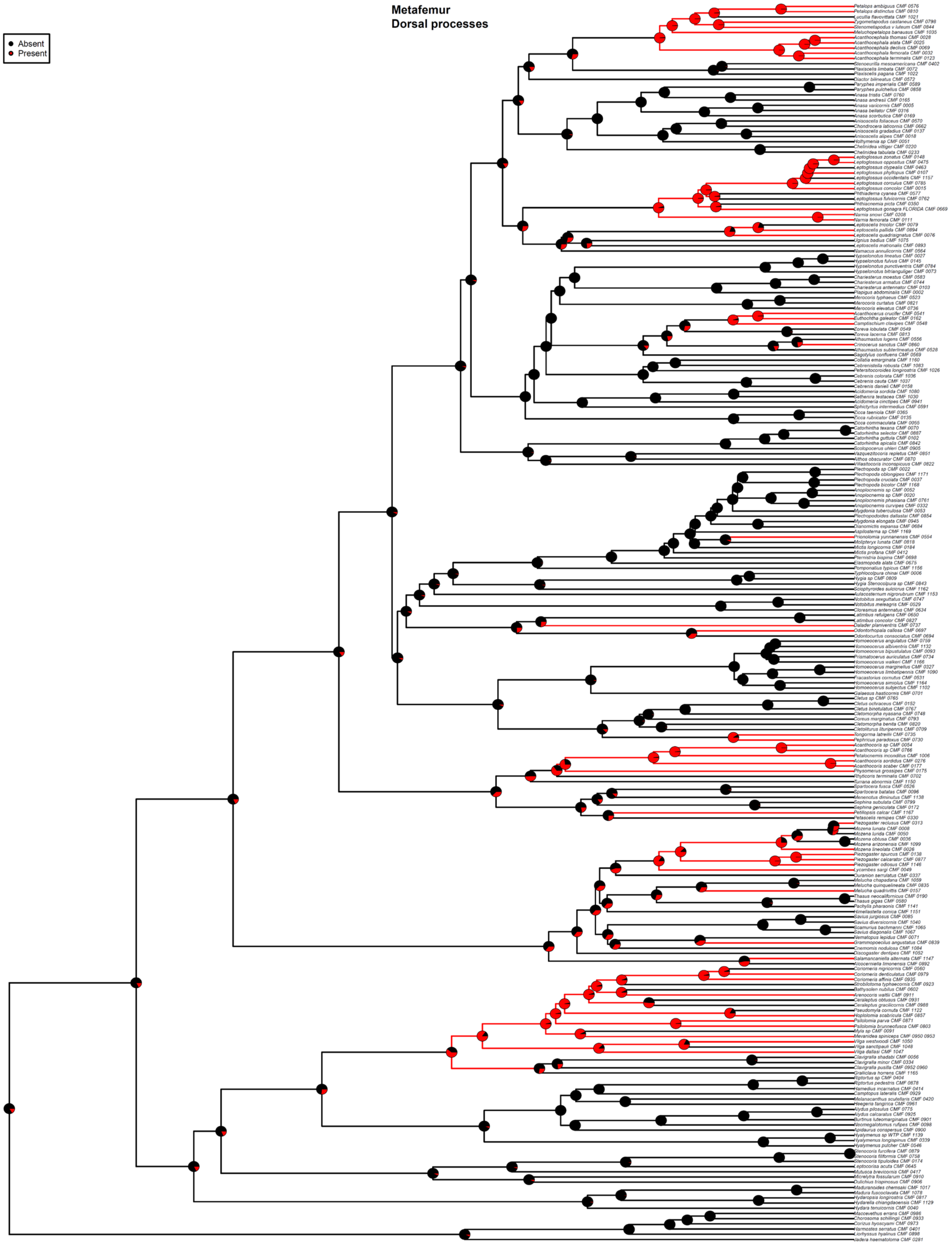


Fig. S31. ASE based on the 50p MSC ultrametric tree and ARD model for Component #8. Taxa with missing data for Component #8 are pruned from the tree for analysis. Pie charts show the likeliest states for a given node (State 0 = black, State 1 = red), with branches similarly colored to represent the most likely state.

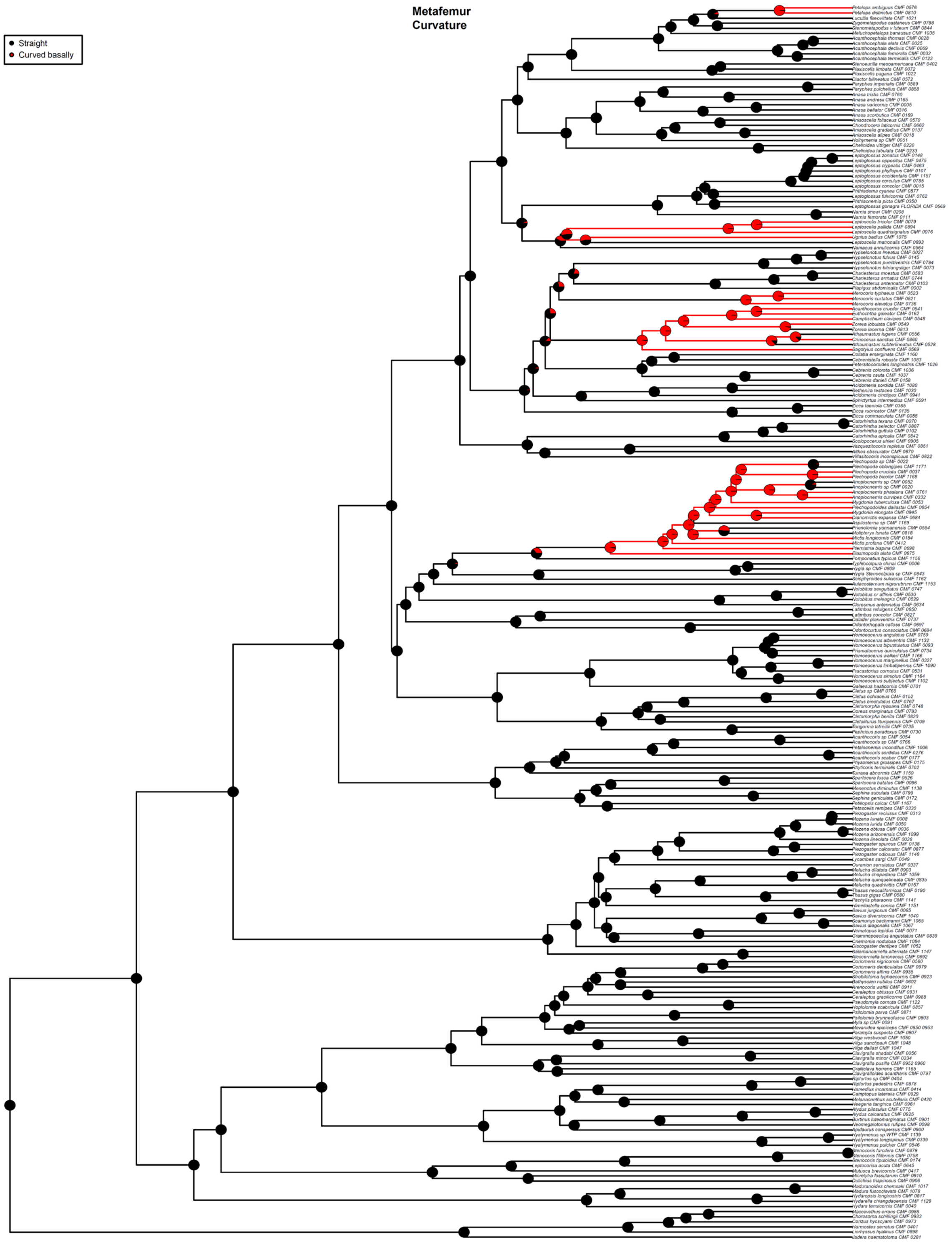


Fig. S32. ASE based on the 50p MSC ultrametric tree and ARD model for Component #9. Taxa with missing data for Component #9 are pruned from the tree for analysis. Pie charts show the likeliest states for a given node (State 0 = black, State 1 = red), with branches similarly colored to represent the most likely state.

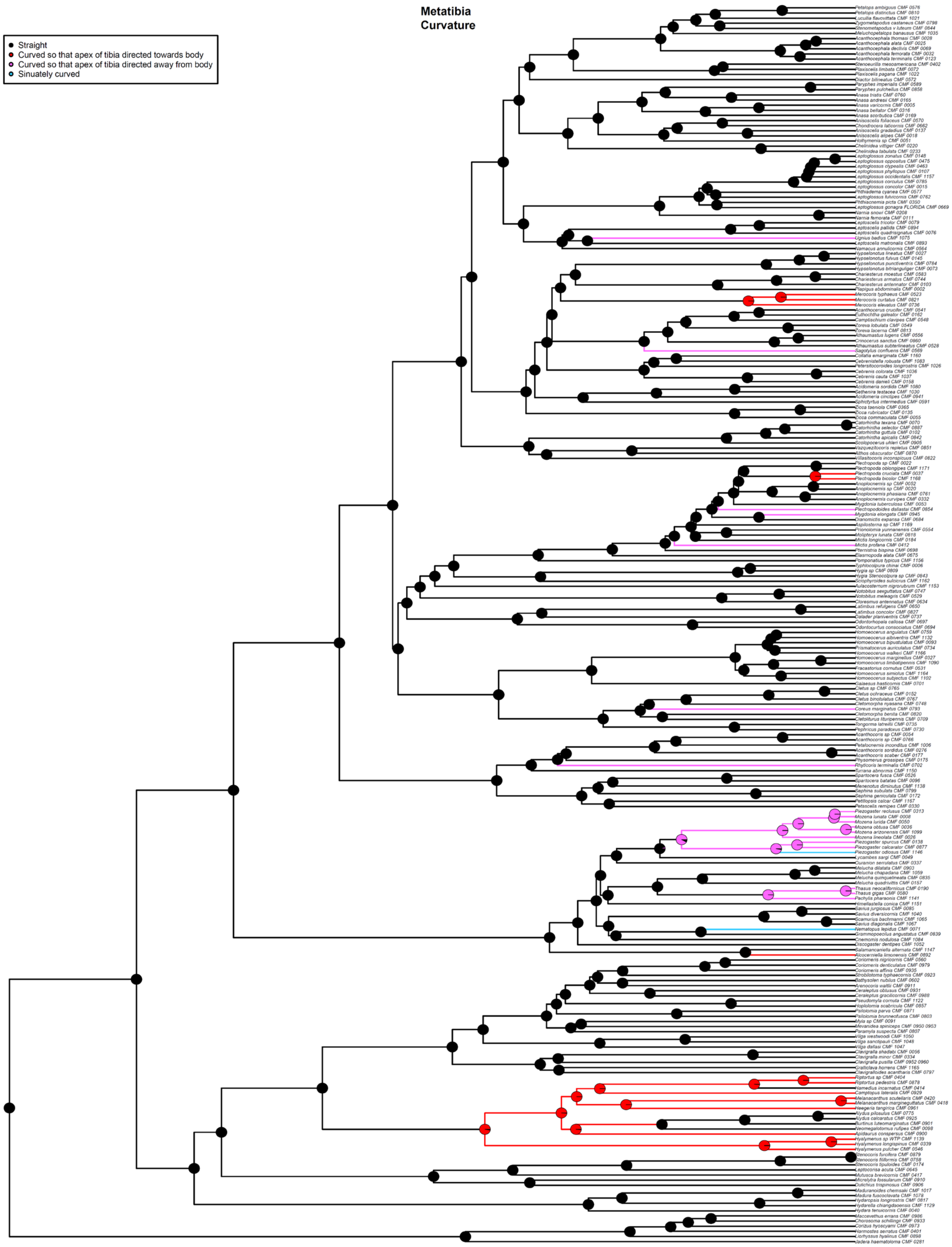


Fig. S33. ASE based on the 50p MSC ultrametric tree and ER model for Component #10. Taxa with missing data for Component #10 are pruned from the tree for analysis. Pie charts show the likeliest states for a given node (State 0 = black, State 1 = red, State 2 = magenta, State 3 = blue), with branches similarly colored to represent the most likely state.

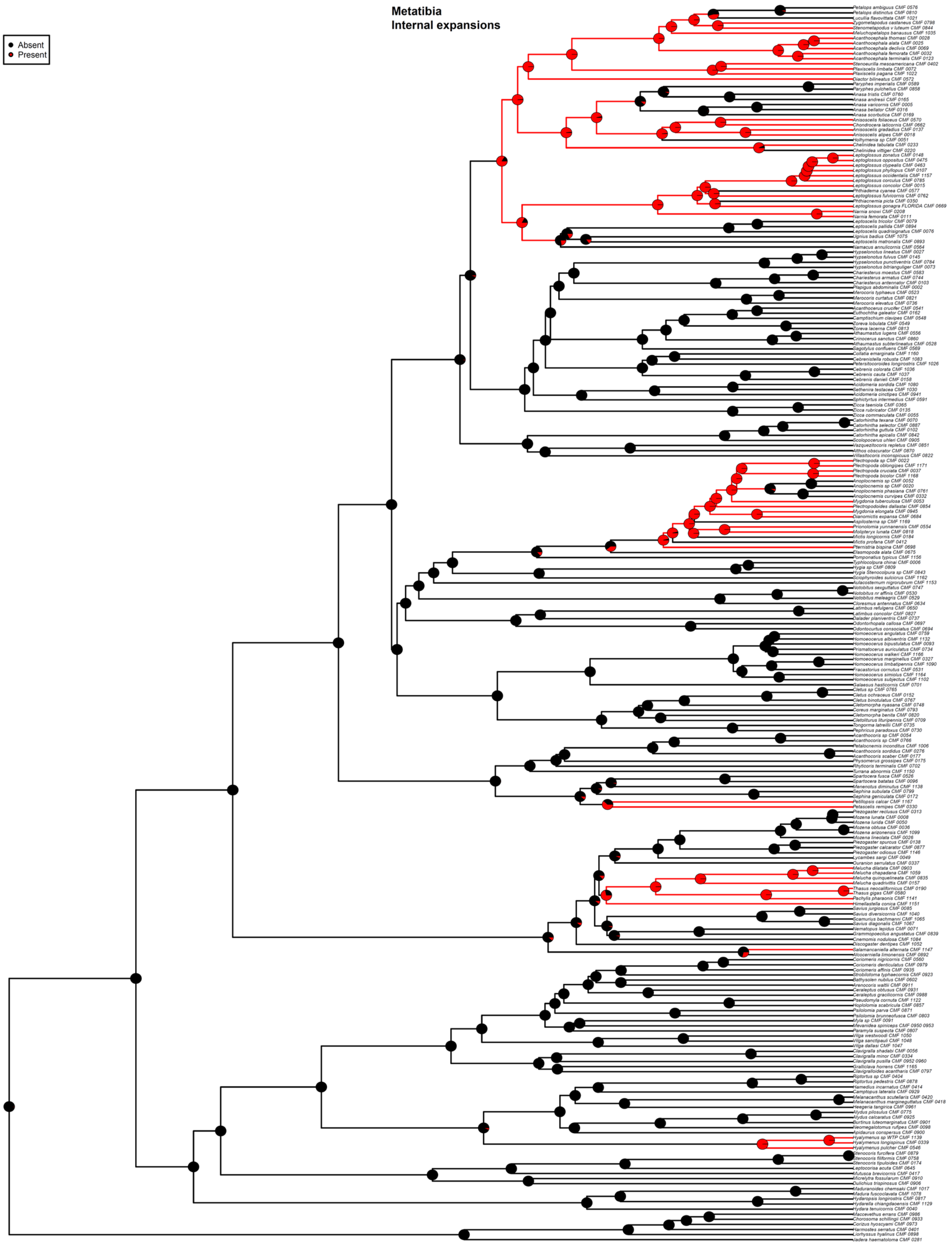


Fig. S34. ASE based on the 50p MSC ultrametric tree and ARD model for Component #11. Taxa with missing data for Component #11 are pruned from the tree for analysis. Pie charts show the likeliest states for a given node (State 0 = black, State 1 = red), with branches similarly colored to represent the most likely state.

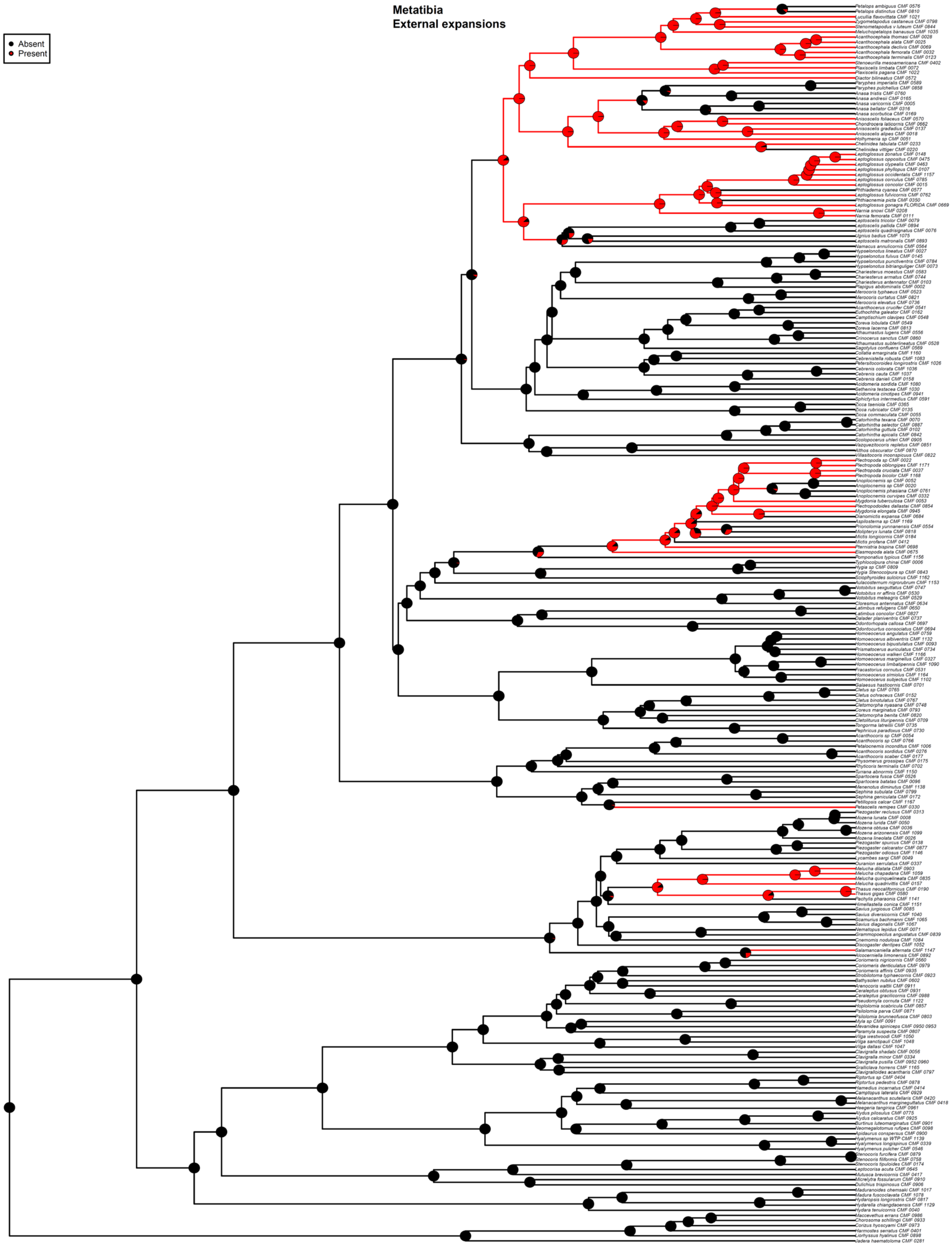


Fig. S35. ASE based on the 50p MSC ultrametric tree and ARD model for Component #12. Taxa with missing data for Component #12 are pruned from the tree for analysis. Pie charts show the likeliest states for a given node (State 0 = black, State 1 = red), with branches similarly colored to represent the most likely state.

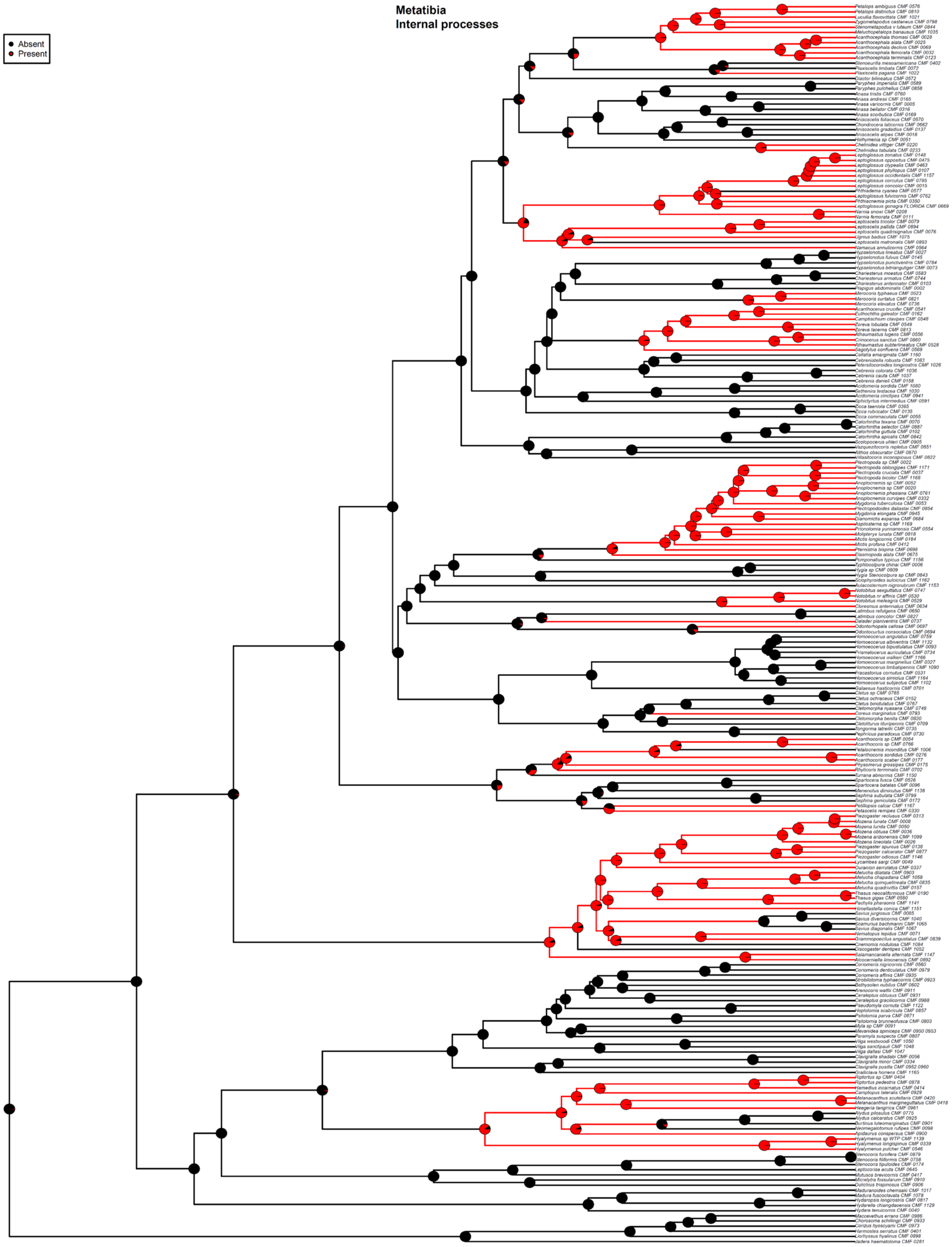


Fig. S36. ASE based on the 50p MSC ultrametric tree and ER model for Component #13. Taxa with missing data for Component #13 are pruned from the tree for analysis. Pie charts show the likeliest states for a given node (State 0 = black, State 1 = red), with branches similarly colored to represent the most likely state.

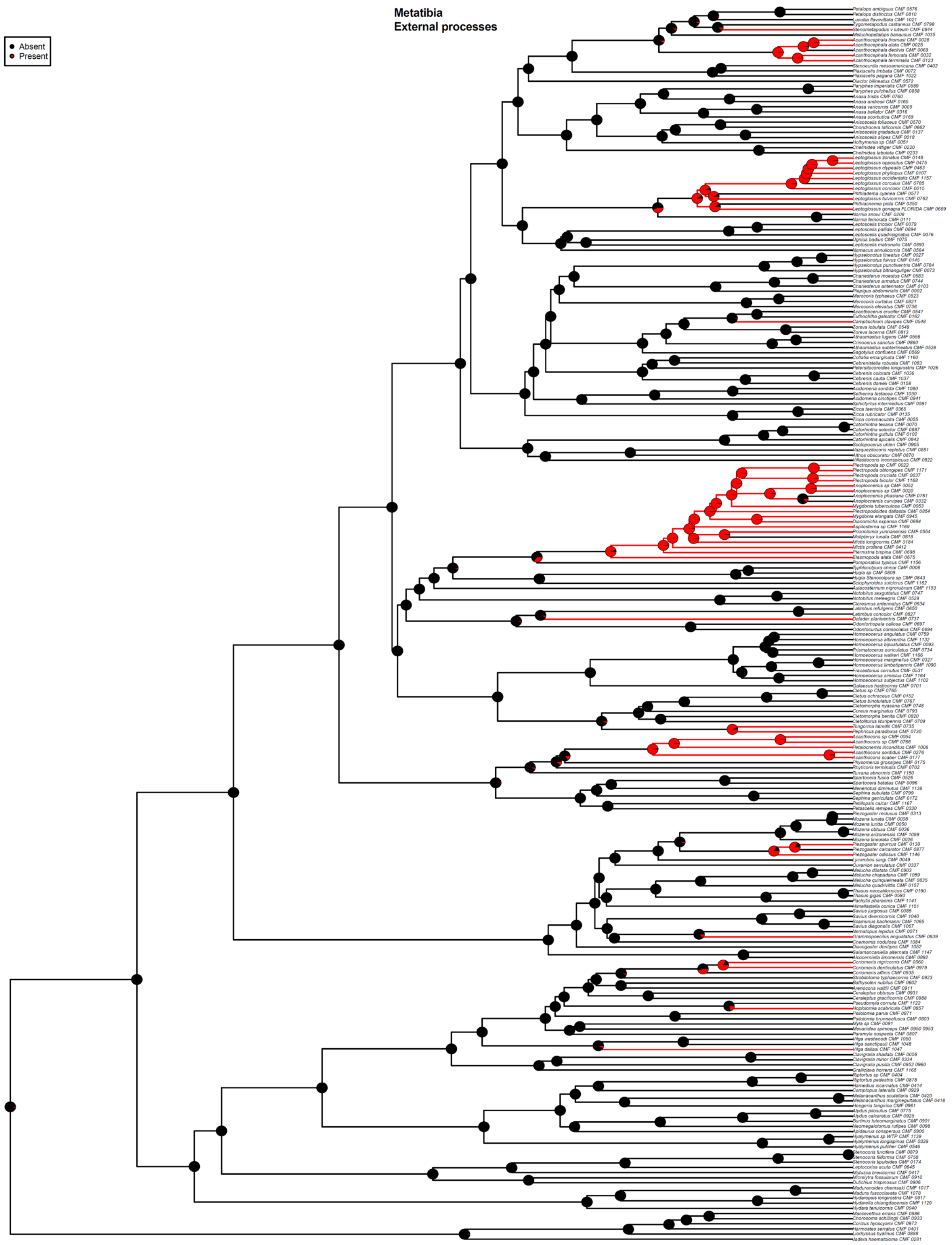


Fig. S37. ASE based on the 50p MSC ultrametric tree and ARD model for Component #14. Taxa with missing data for Component #14 are pruned from the tree for analysis. Pie charts show the likeliest states for a given node (State 0 = black, State 1 = red), with branches similarly colored to represent the most likely state.

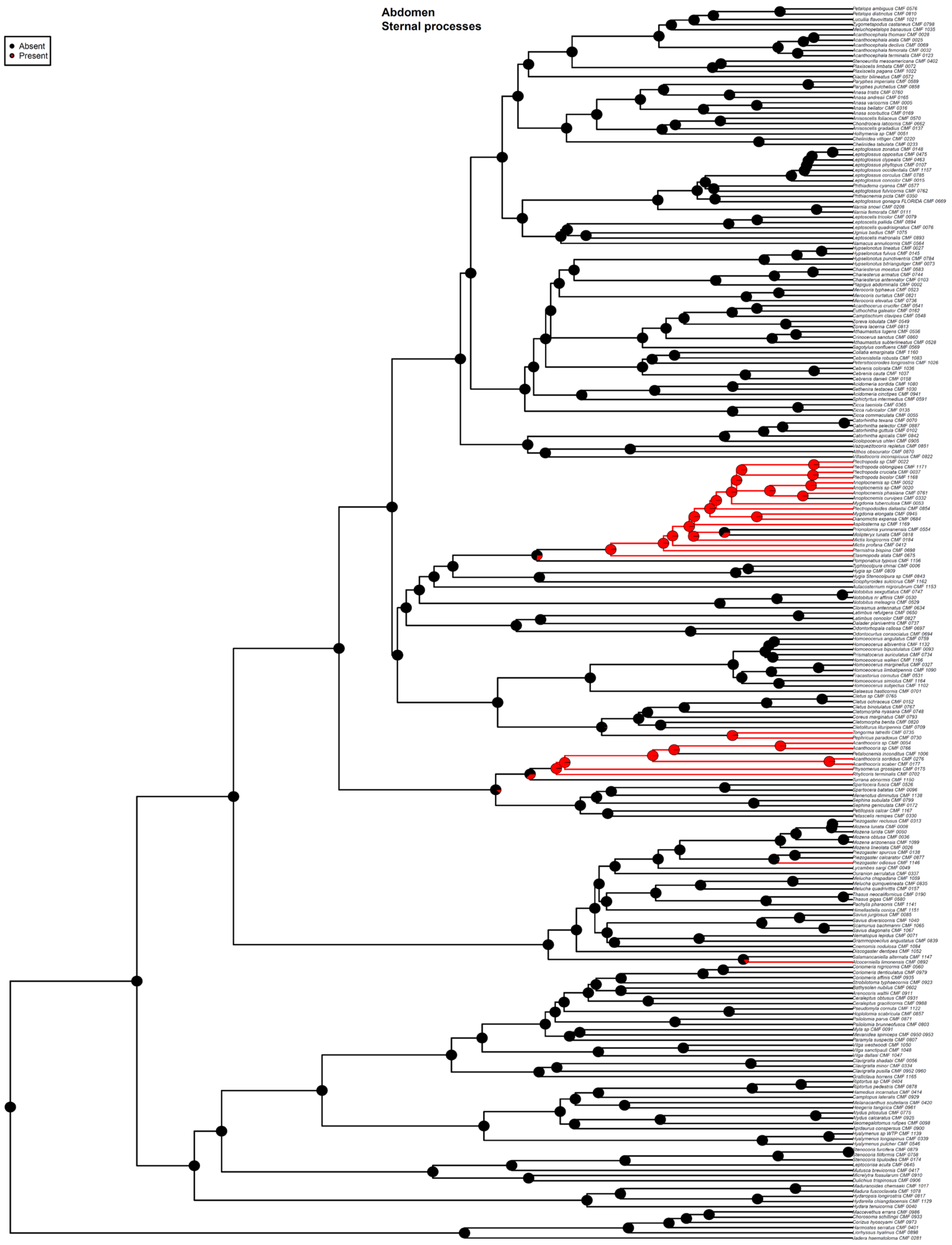


Fig. S38. ASE based on the 50p MSC ultrametric tree and ARD model for Component #15. Taxa with missing data for Component #15 are pruned from the tree for analysis. Pie charts show the likeliest states for a given node (State 0 = black, State 1 = red), with branches similarly colored to represent the most likely state.

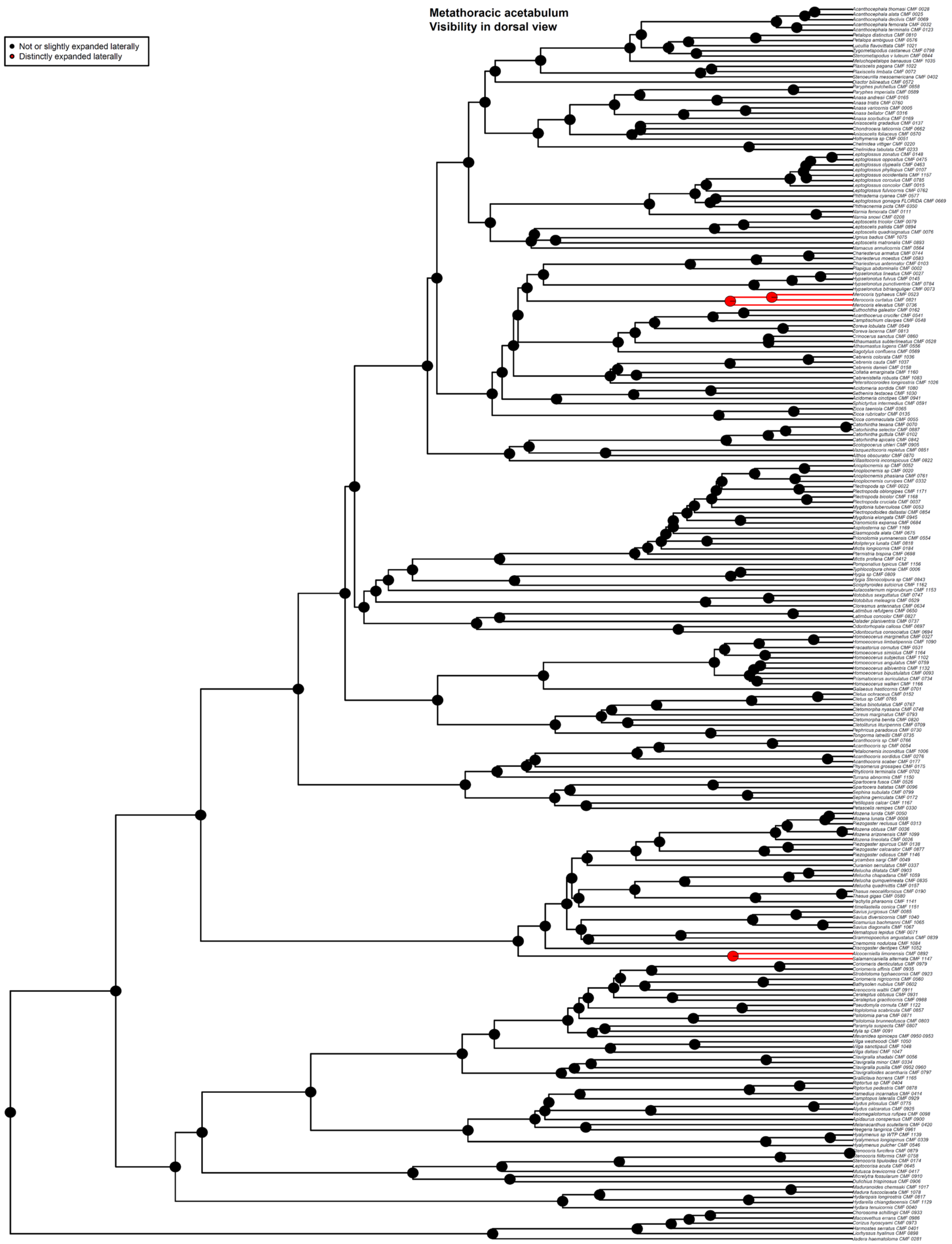


Fig. S39. ASE based on the 50p25mi MSC ultrametric tree and ER model for Component #1. Taxa with missing data for Component #1 are pruned from the tree for analysis. Pie charts show the likeliest states for a given node (State 0 = black, State 1 = red), with branches similarly colored to represent the most likely state.

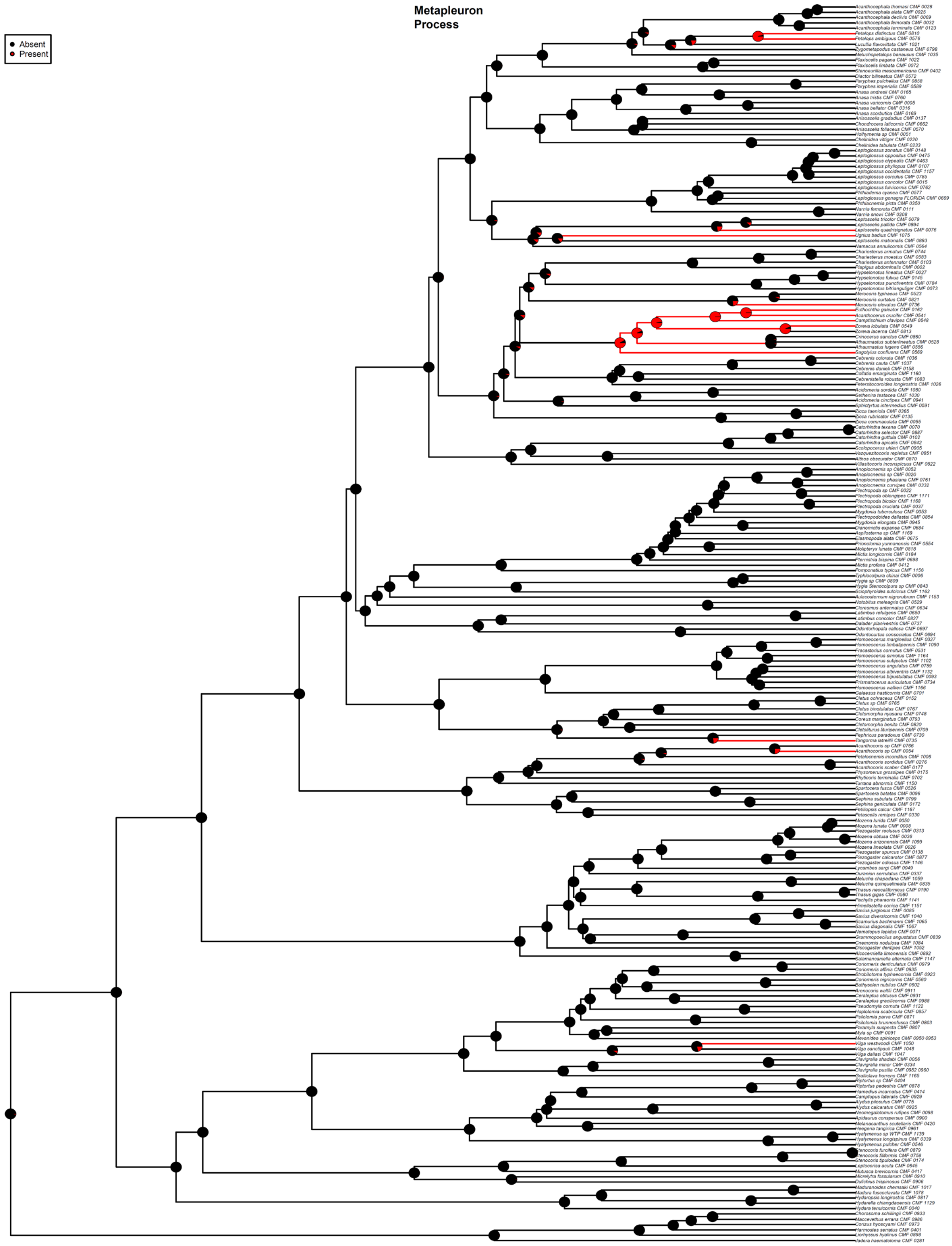


Fig. S40. ASE based on the 50p25mi MSC ultrametric tree and ARD model for Component #2. Taxa with missing data for Component #2 are pruned from the tree for analysis. Pie charts show the likeliest states for a given node (State 0 = black, State 1 = red), with branches similarly colored to represent the most likely state.

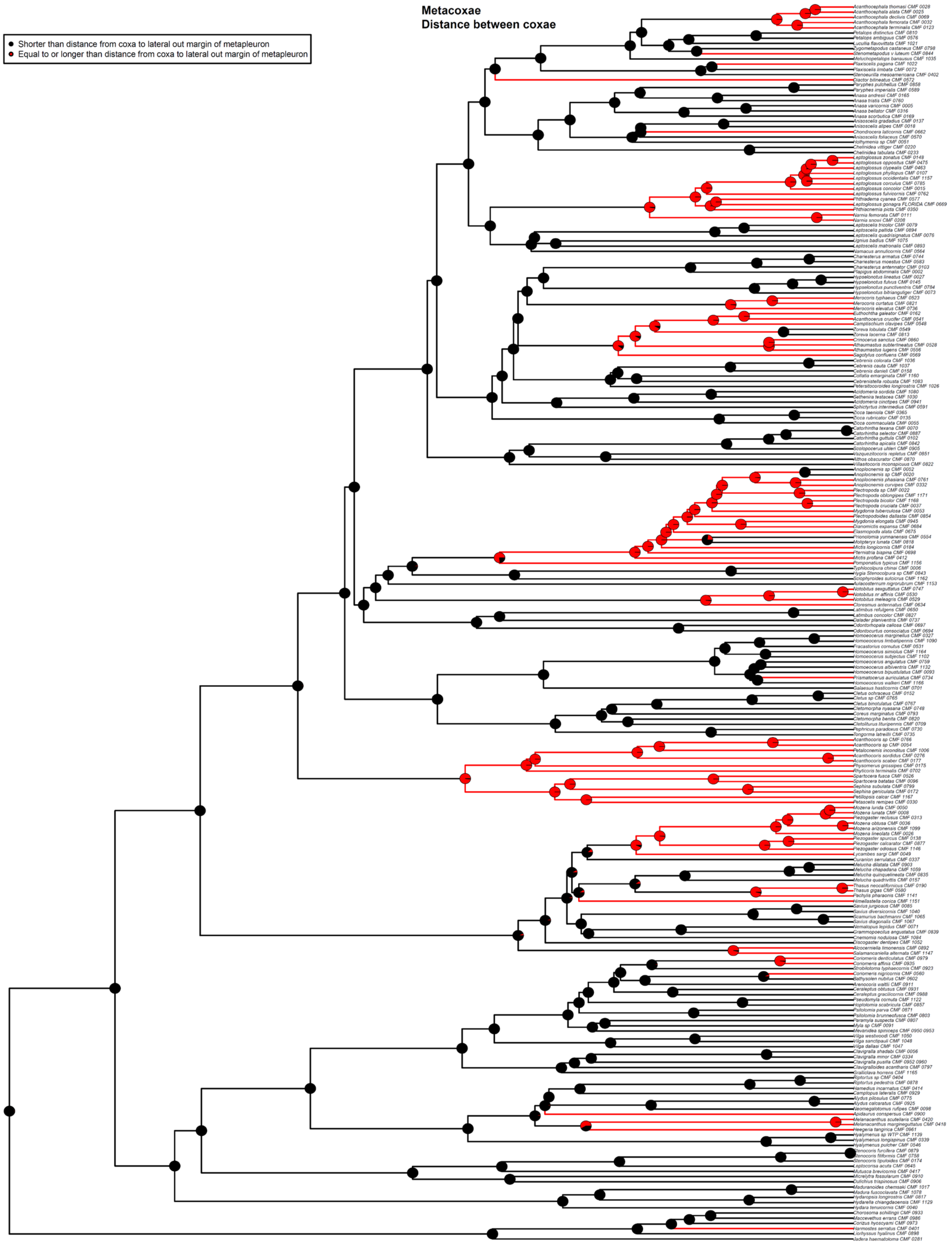


Fig. S41. ASE based on the 50p25mi MSC ultrametric tree and ER model for Component #3. Taxa with missing data for Component #3 are pruned from the tree for analysis. Pie charts show the likeliest states for a given node (State 0 = black, State 1 = red), with branches similarly colored to represent the most likely state.

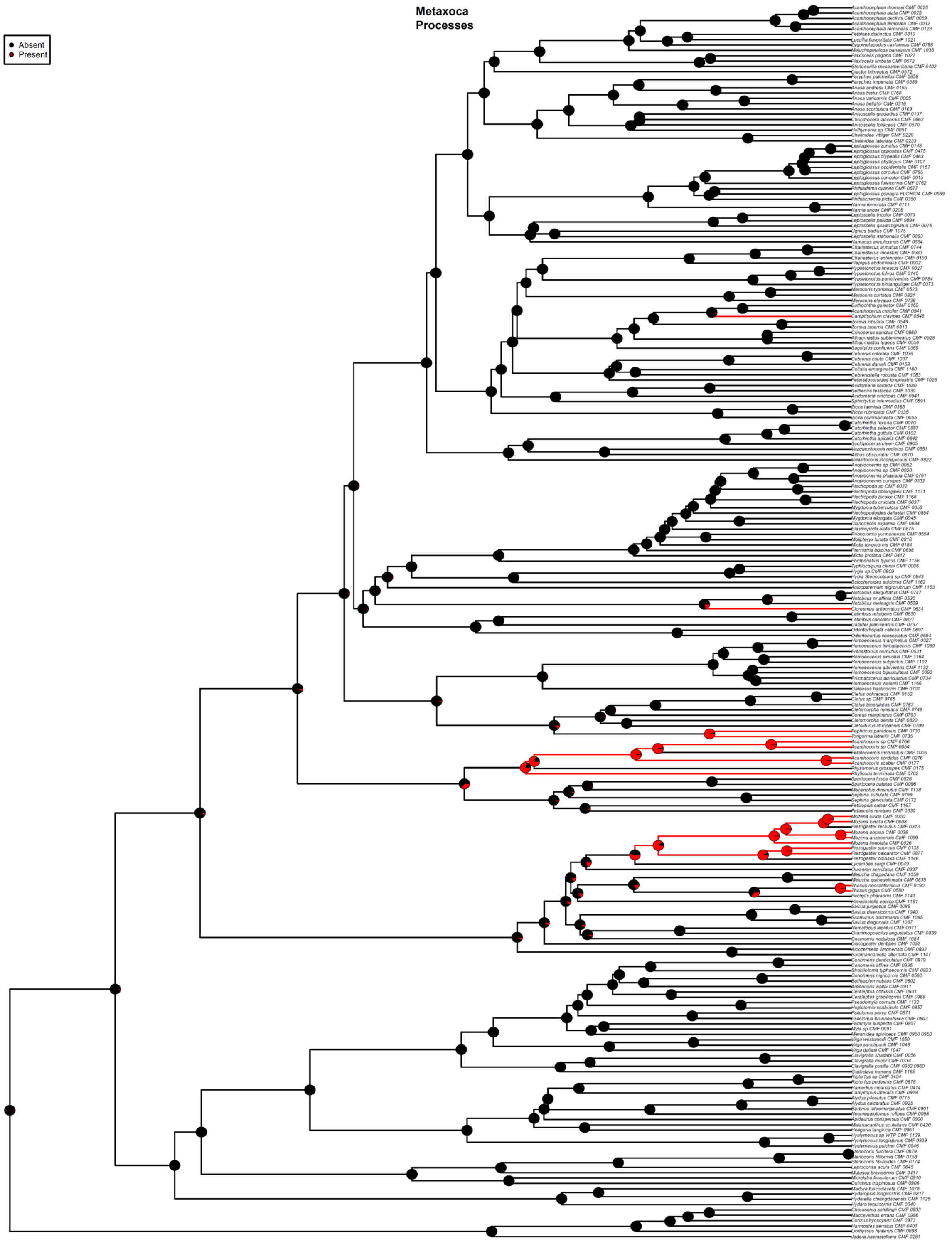


Fig. S42. ASE based on the 50p25mi MSC ultrametric tree and ARD model for Component #4. Taxa with missing data for Component #4 are pruned from the tree for analysis. Pie charts show the likeliest states for a given node (State 0 = black, State 1 = red), with branches similarly colored to represent the most likely state.

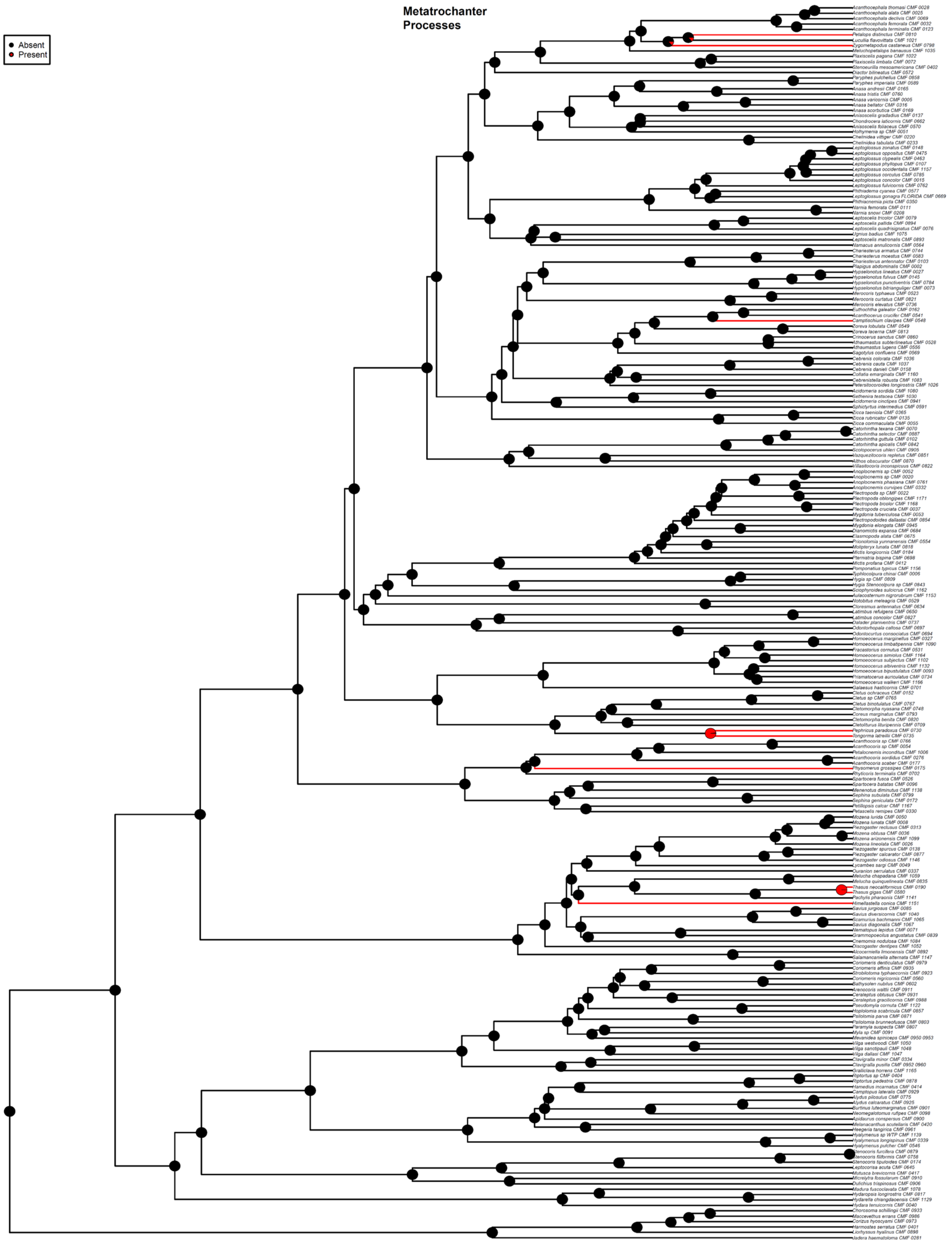


Fig. S43. ASE based on the 50p25mi MSC ultrametric tree and ER model for Component #5. Taxa with missing data for Component #5 are pruned from the tree for analysis. Pie charts show the likeliest states for a given node (State 0 = black, State 1 = red), with branches similarly colored to represent the most likely state.

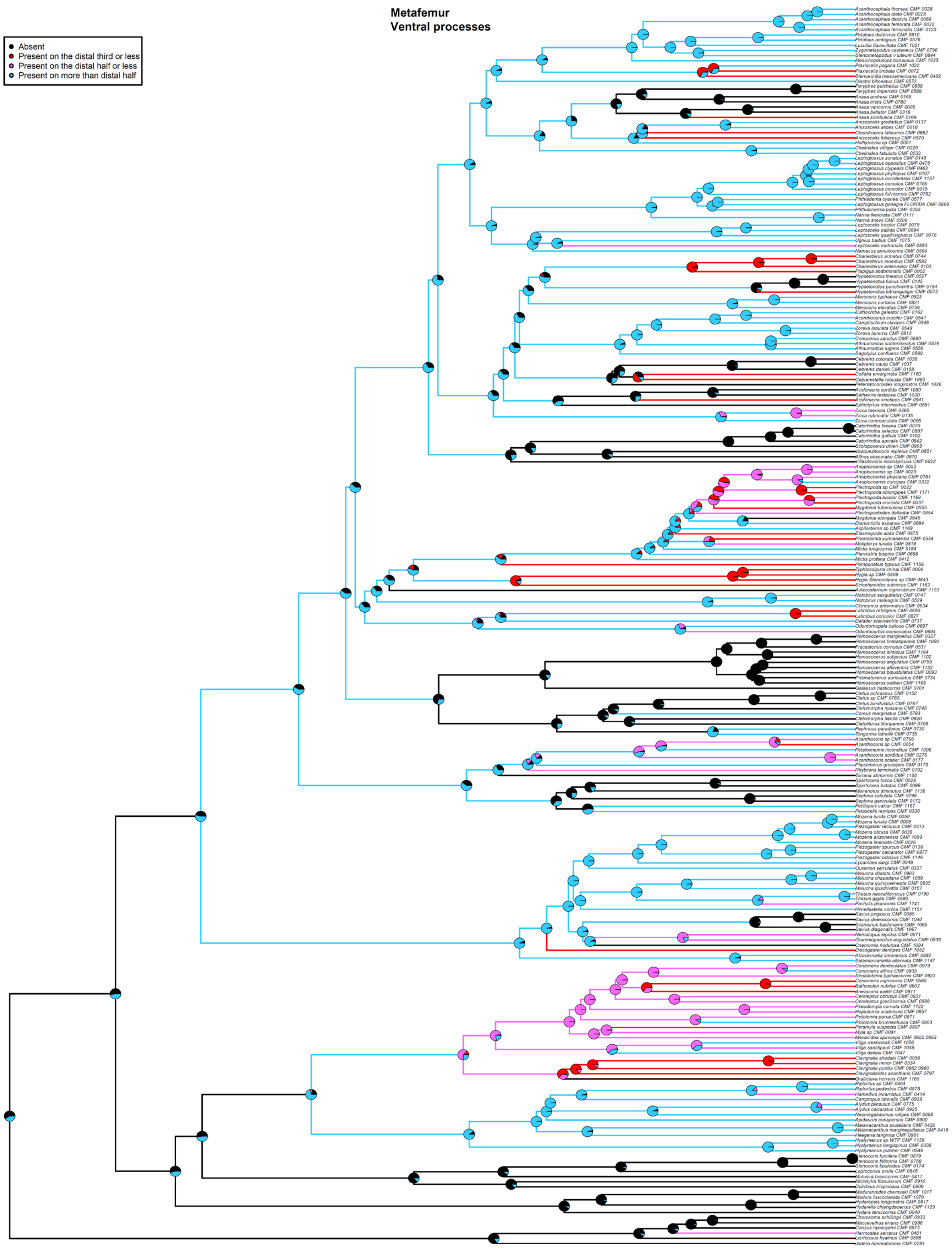


Fig. S44. ASE based on the 50p25mi MSC ultrametric tree and SYM model for Component #6. Taxa with missing data for Component #6 are pruned from the tree for analysis. Pie charts show the likeliest states for a given node (State 0 = black, State 1 = red, State 2 = magenta, State 3 = blue), with branches similarly colored to represent the most likely state.

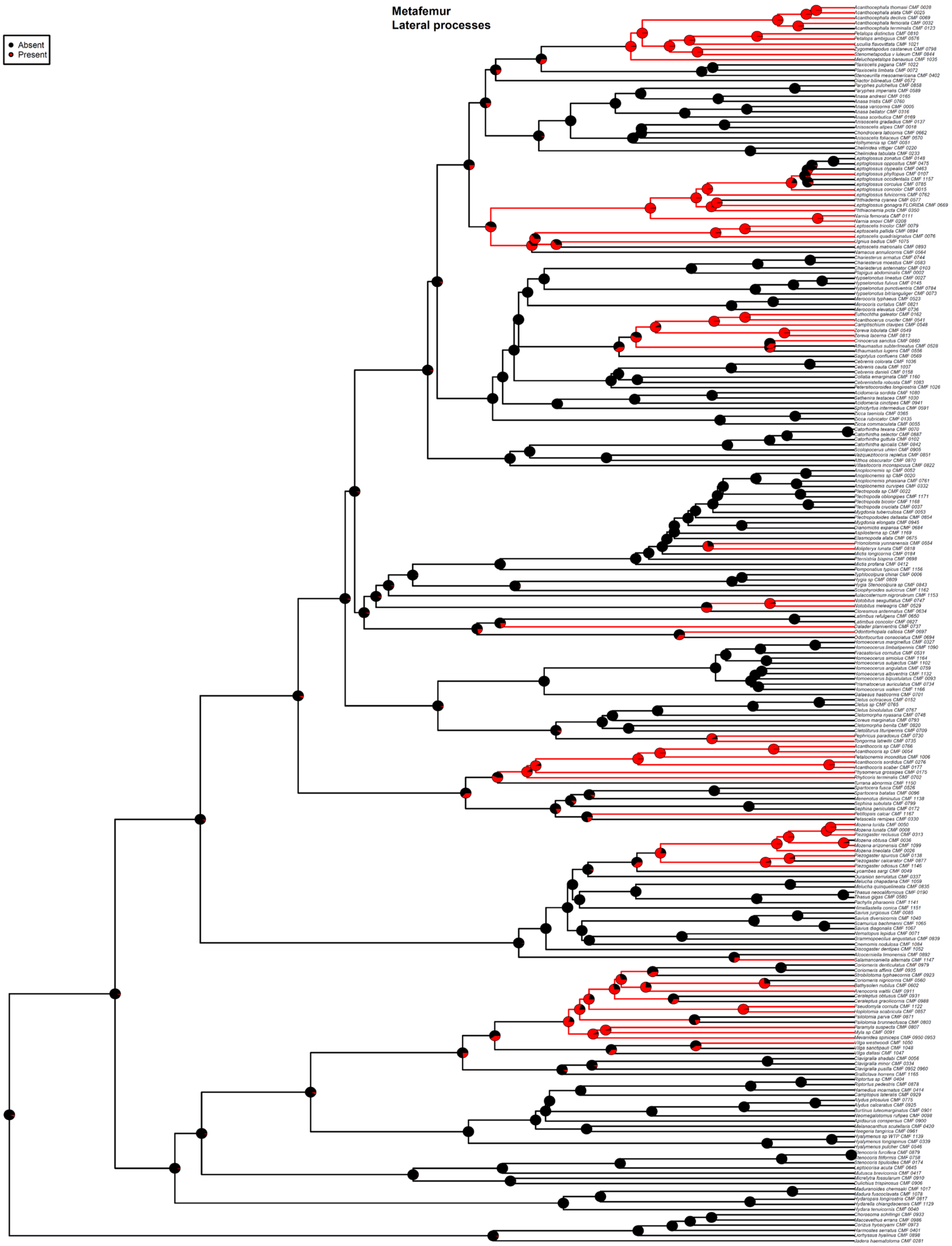


Fig. S45. ASE based on the 50p25mi MSC ultrametric tree and ARD model for Component #7. Taxa with missing data for Component #7 are pruned from the tree for analysis. Pie charts show the likeliest states for a given node (State 0 = black, State 1 = red), with branches similarly colored to represent the most likely state.

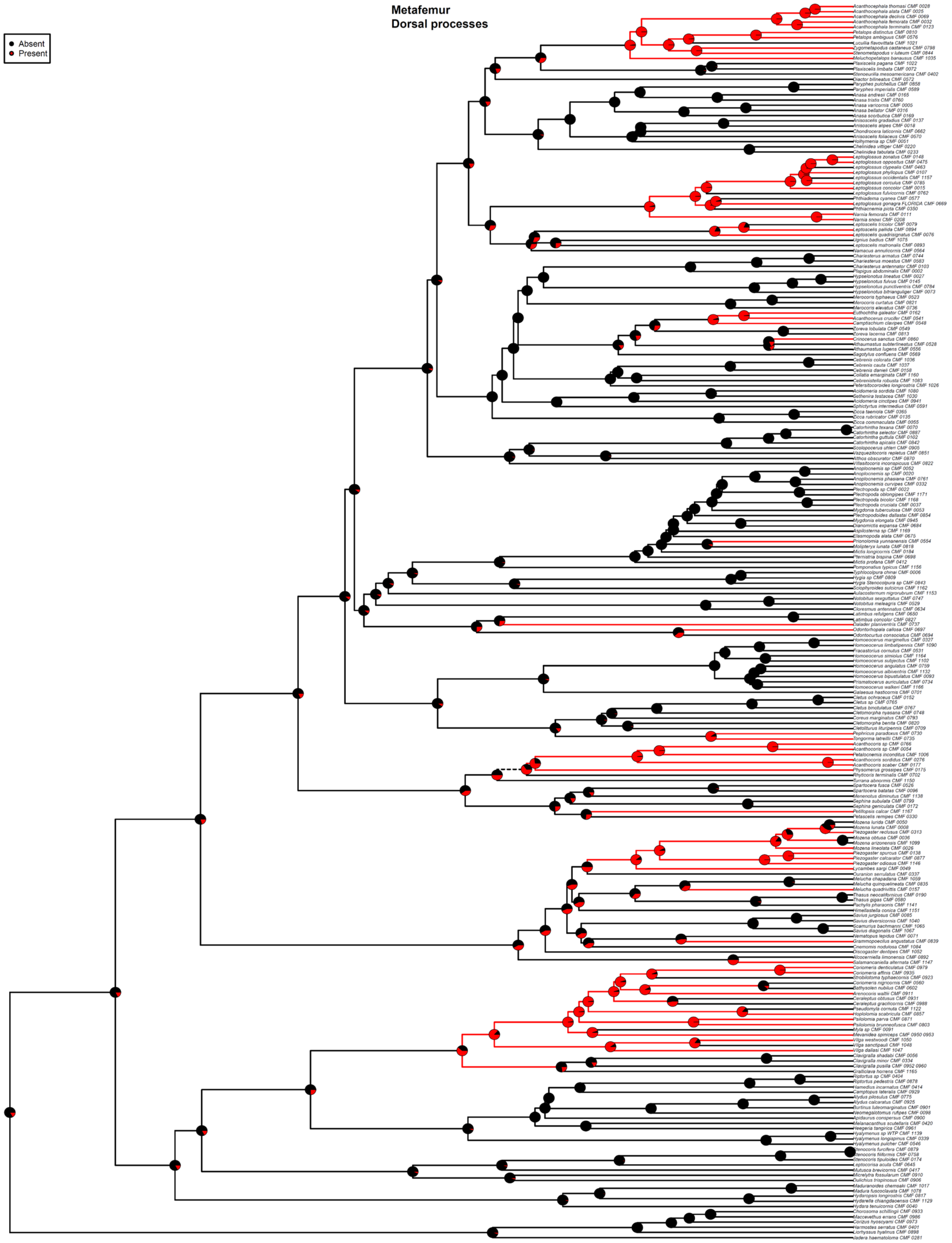


Fig. S46. ASE based on the 50p25mi MSC ultrametric tree and ARD model for Component #8. Taxa with missing data for Component #8 are pruned from the tree for analysis. Pie charts show the likeliest states for a given node (State 0 = black, State 1 = red), with branches similarly colored to represent the most likely state.

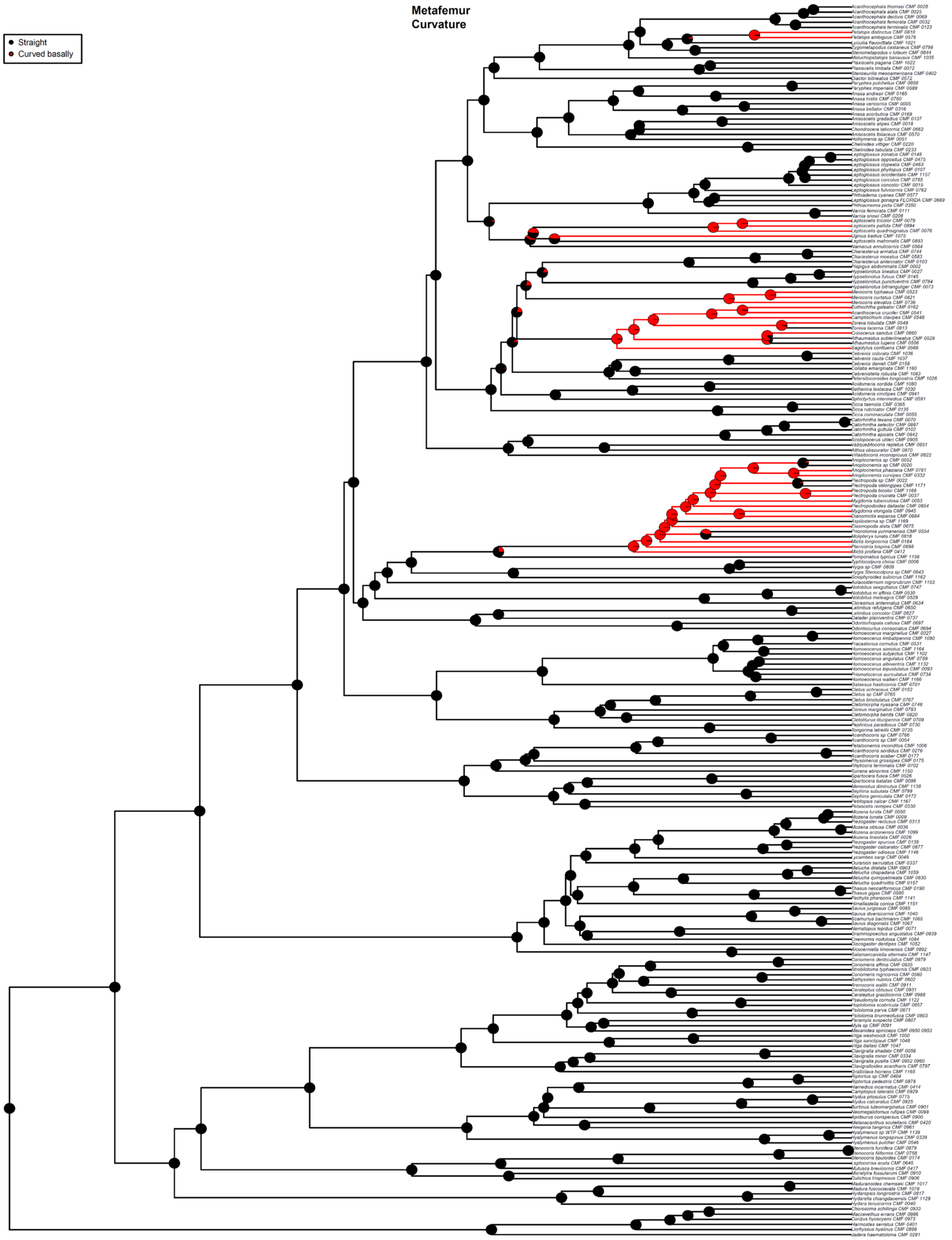


Fig. S47. ASE based on the 50p25mi MSC ultrametric tree and ARD model for Component #9. Taxa with missing data for Component #9 are pruned from the tree for analysis. Pie charts show the likeliest states for a given node (State 0 = black, State 1 = red), with branches similarly colored to represent the most likely state.

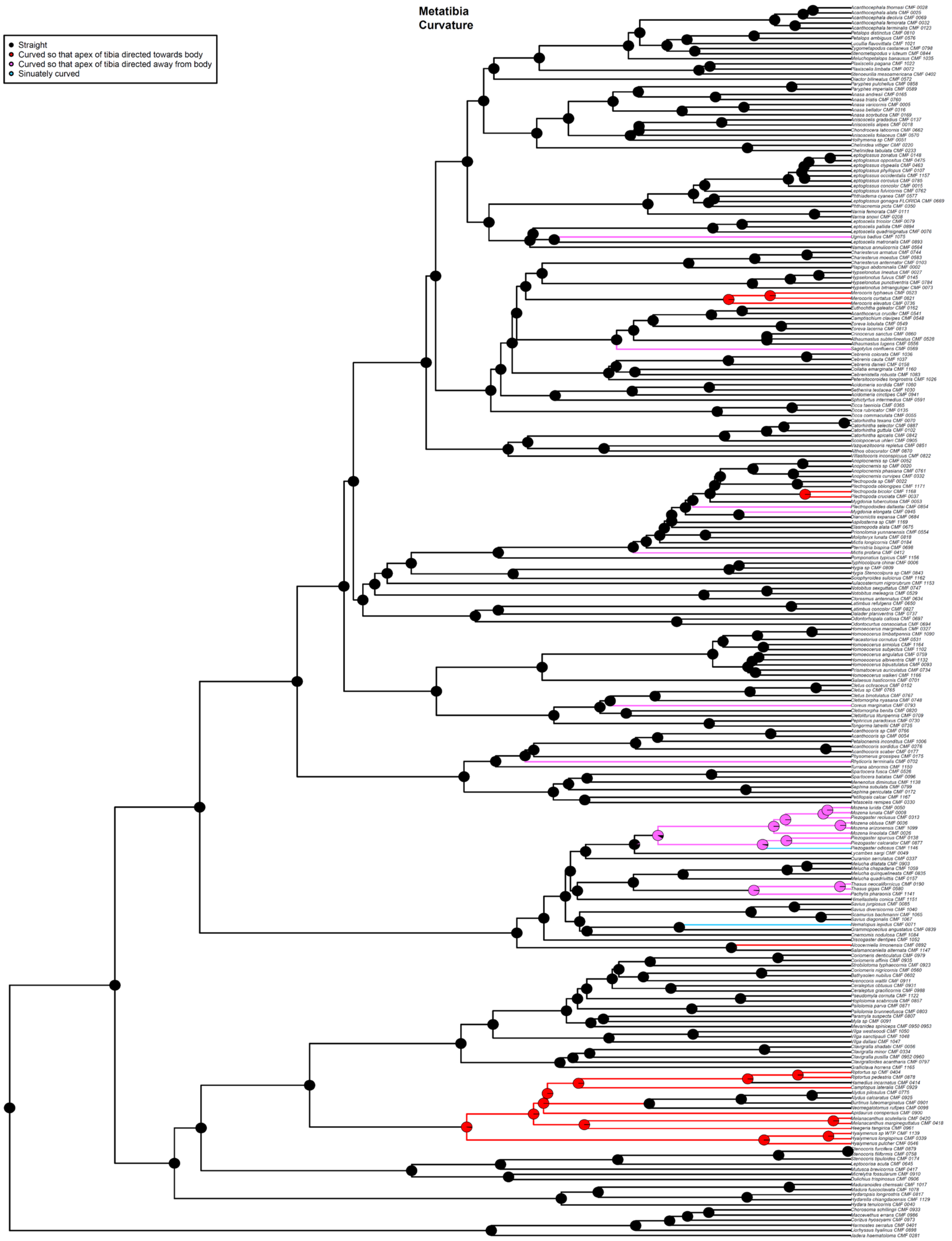


Fig. S48. ASE based on the 50p25mi MSC ultrametric tree and ER model for Component #10. Taxa with missing data for Component #10 are pruned from the tree for analysis. Pie charts show the likeliest states for a given node (State 0 = black, State 1 = red, State 2 = magenta, State 3 = blue), with branches similarly colored to represent the most likely state.

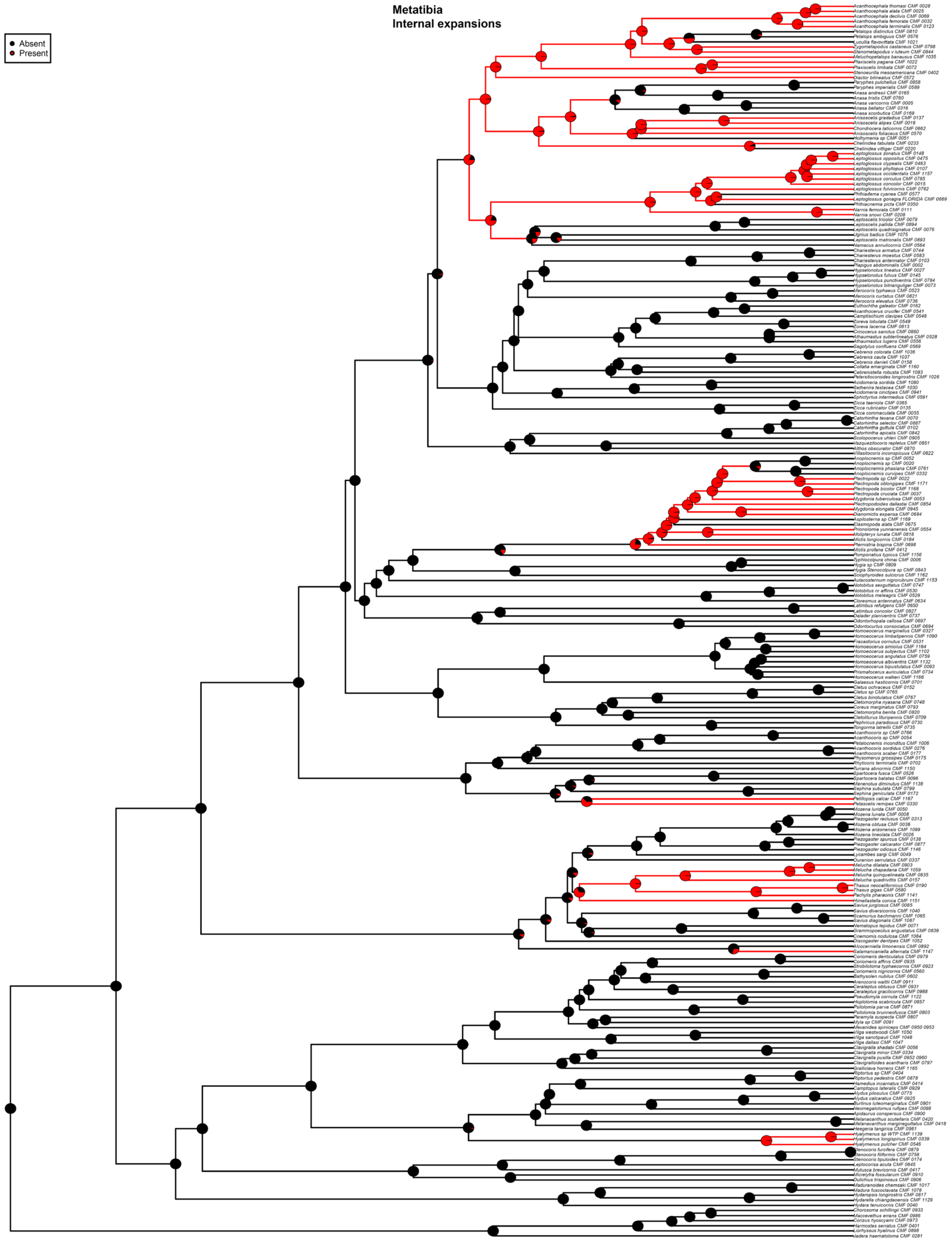


Fig. S49. ASE based on the 50p25mi MSC ultrametric tree and ARD model for Component #11. Taxa with missing data for Component #11 are pruned from the tree for analysis. Pie charts show the likeliest states for a given node (State 0 = black, State 1 = red), with branches similarly colored to represent the most likely state.

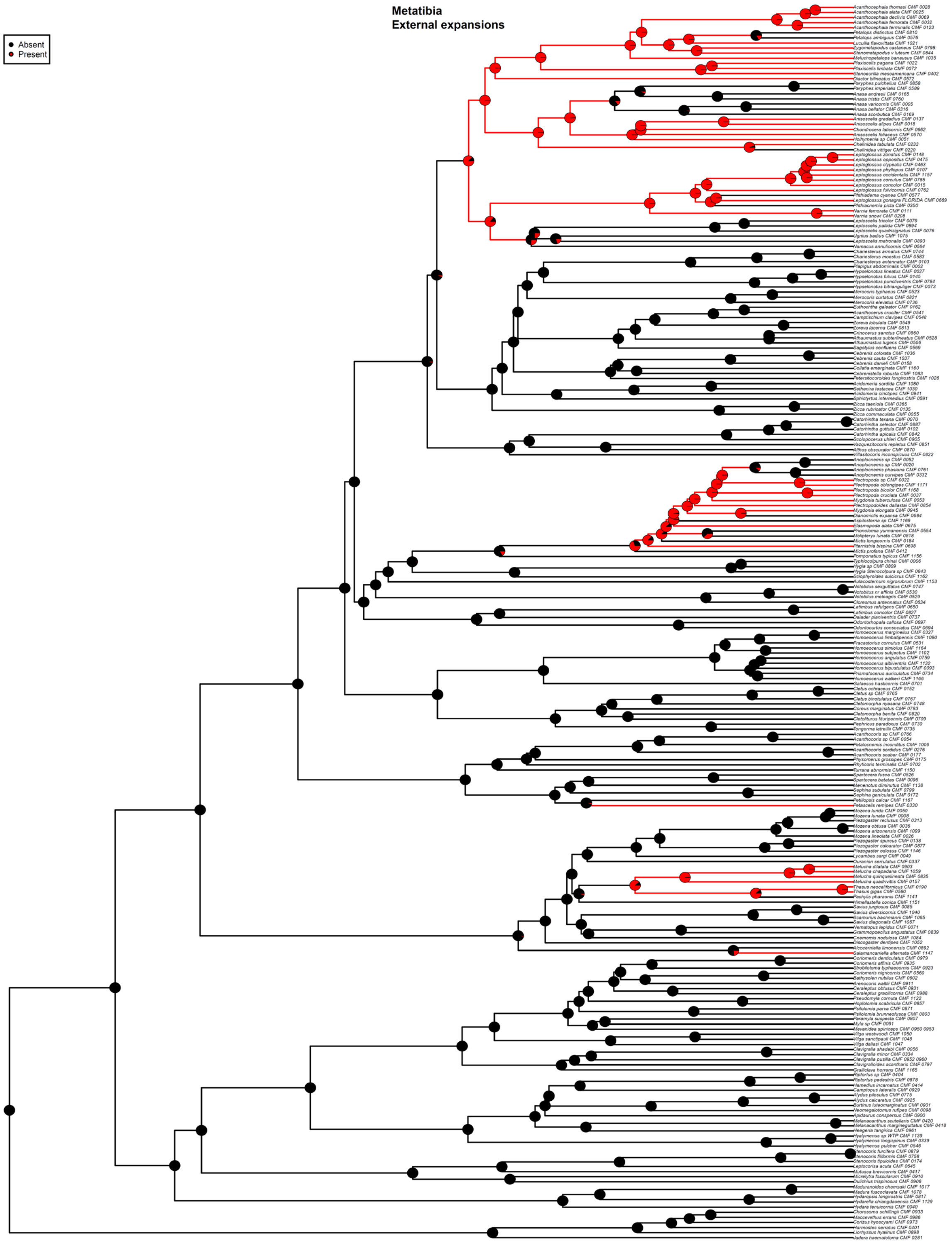


Fig. S50. ASE based on the 50p25mi MSC ultrametric tree and ARD model for Component #12. Taxa with missing data for Component #12 are pruned from the tree for analysis. Pie charts show the likeliest states for a given node (State 0 = black, State 1 = red), with branches similarly colored to represent the most likely state.

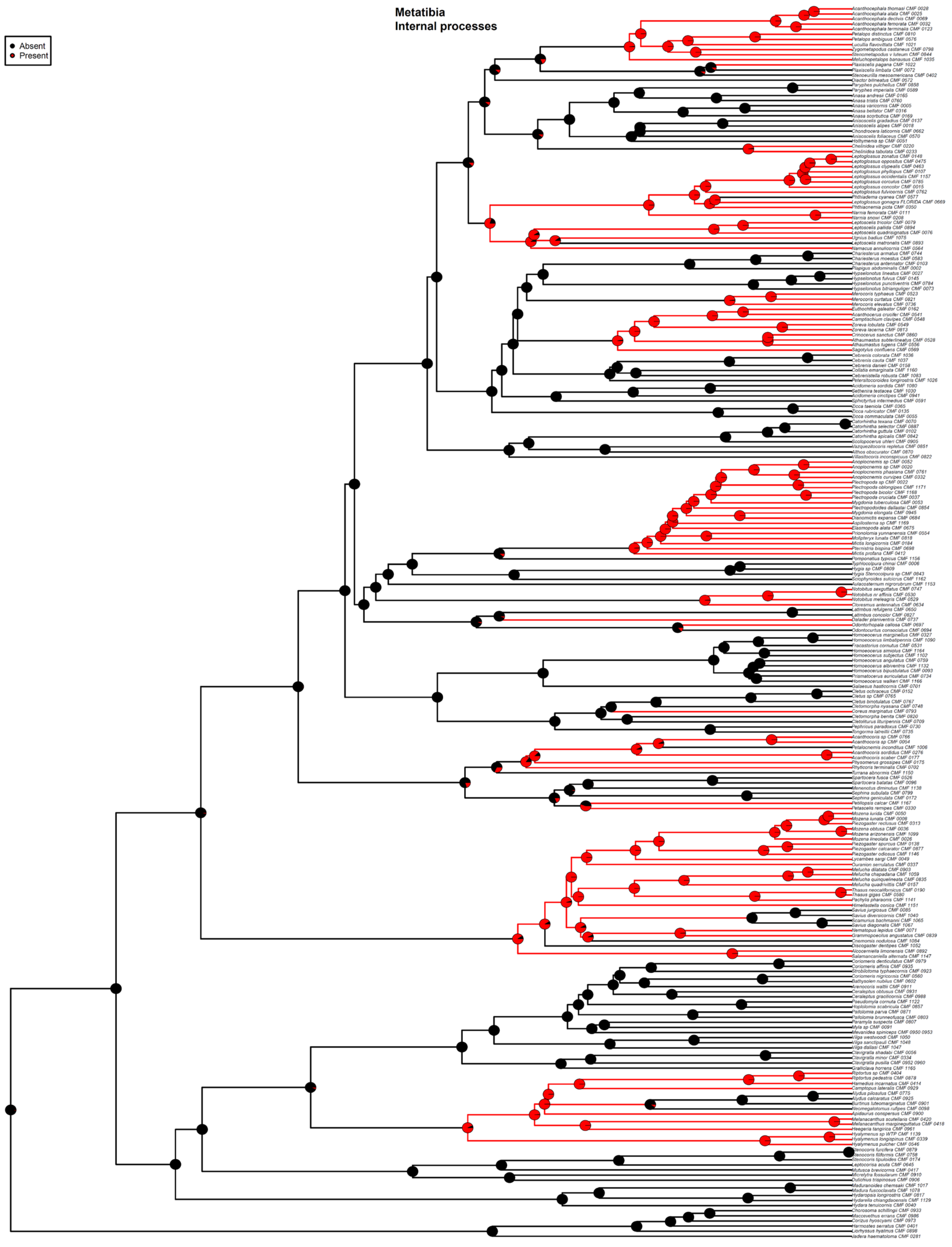


Fig. S51. ASE based on the 50p25mi MSC ultrametric tree and ER model for Component #13. Taxa with missing data for Component #13 are pruned from the tree for analysis. Pie charts show the likeliest states for a given node (State 0 = black, State 1 = red), with branches similarly colored to represent the most likely state.

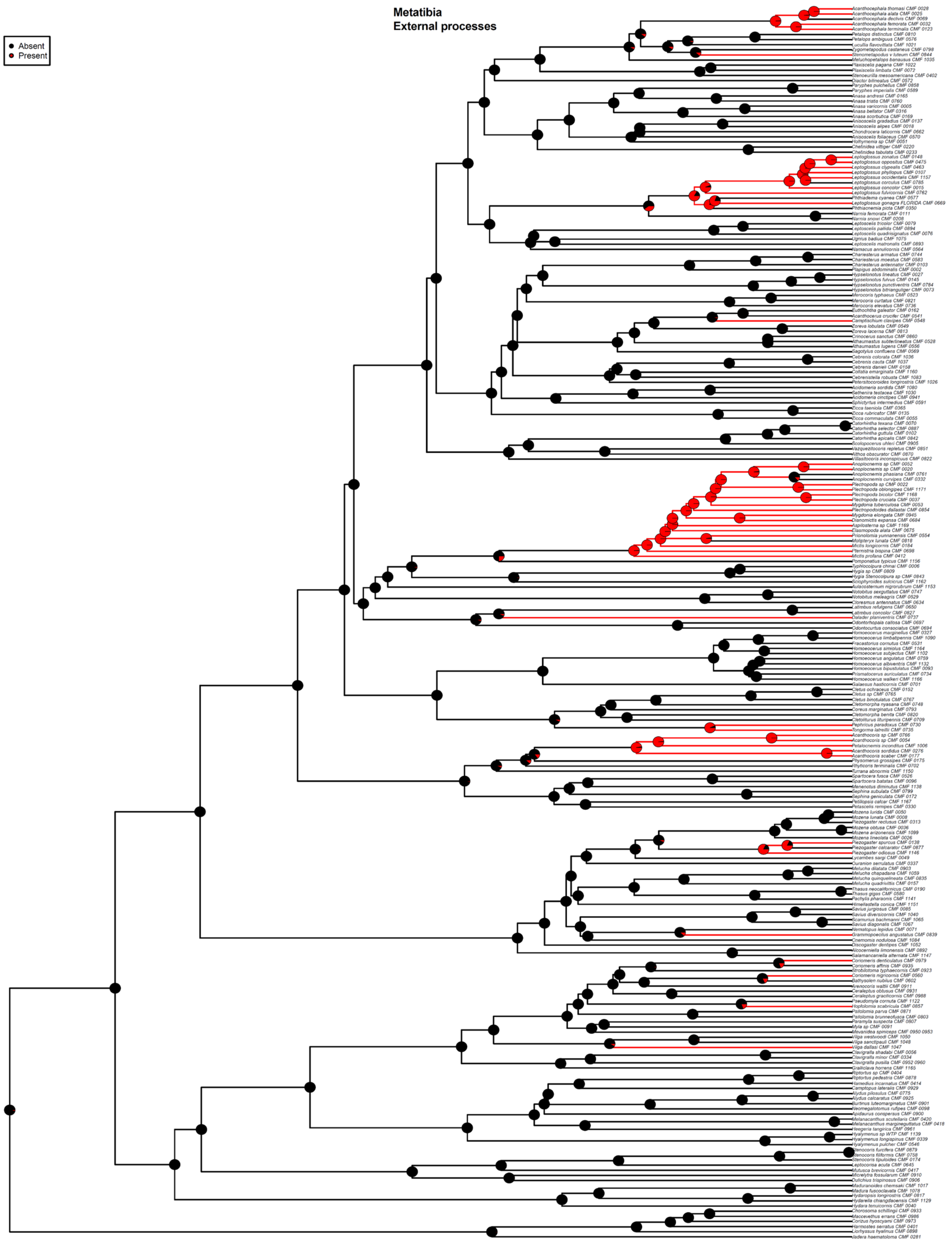


Fig. S52. ASE based on the 50p25mi MSC ultrametric tree and ARD model for Component #14. Taxa with missing data for Component #14 are pruned from the tree for analysis. Pie charts show the likeliest states for a given node (State 0 = black, State 1 = red), with branches similarly colored to represent the most likely state.

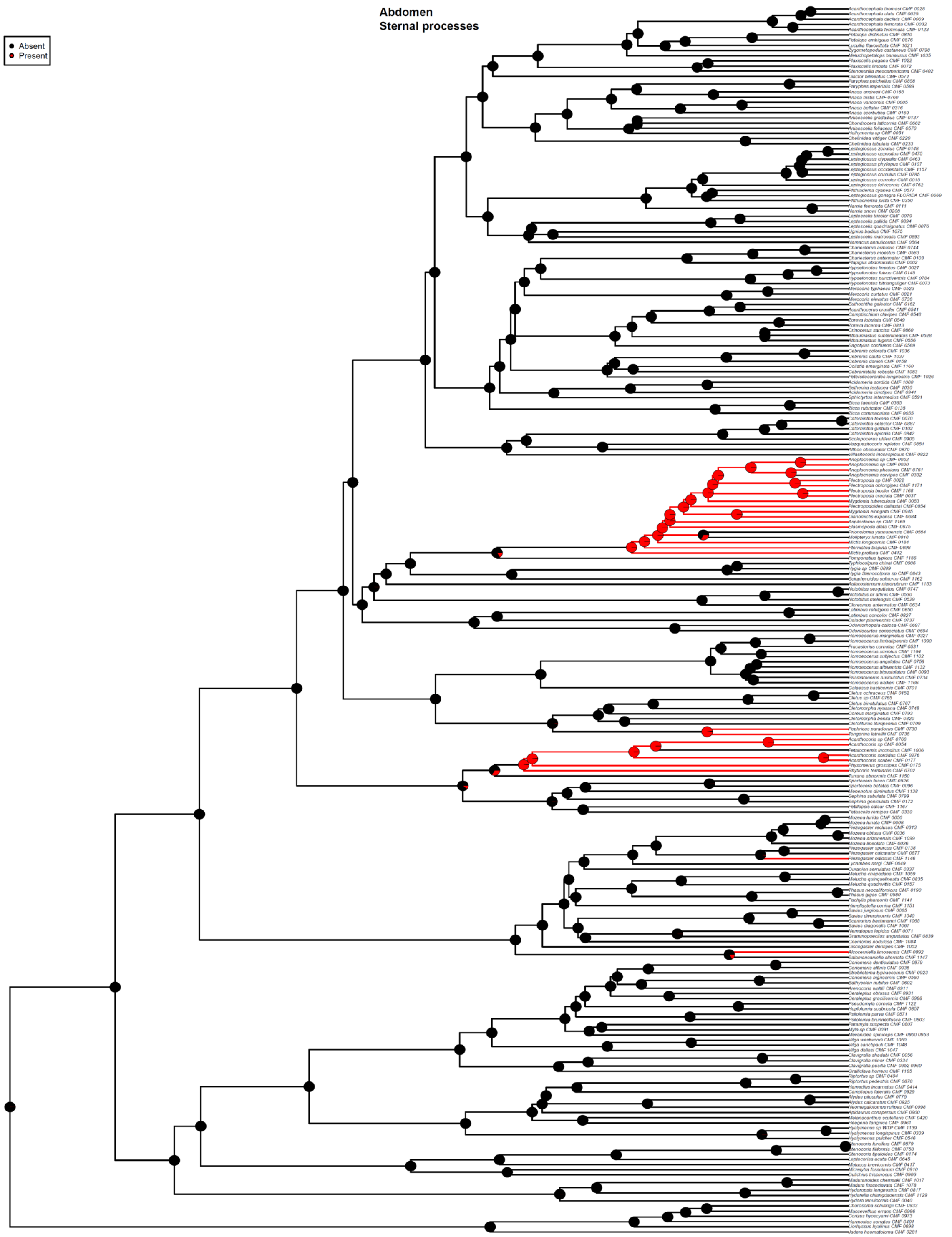


Fig. S53. ASE based on the 50p25mi MSC ultrametric tree and ARD model for Component #15. Taxa with missing data for Component #15 are pruned from the tree for analysis. Pie charts show the likeliest states for a given node (State 0 = black, State 1 = red), with branches similarly colored to represent the most likely state.

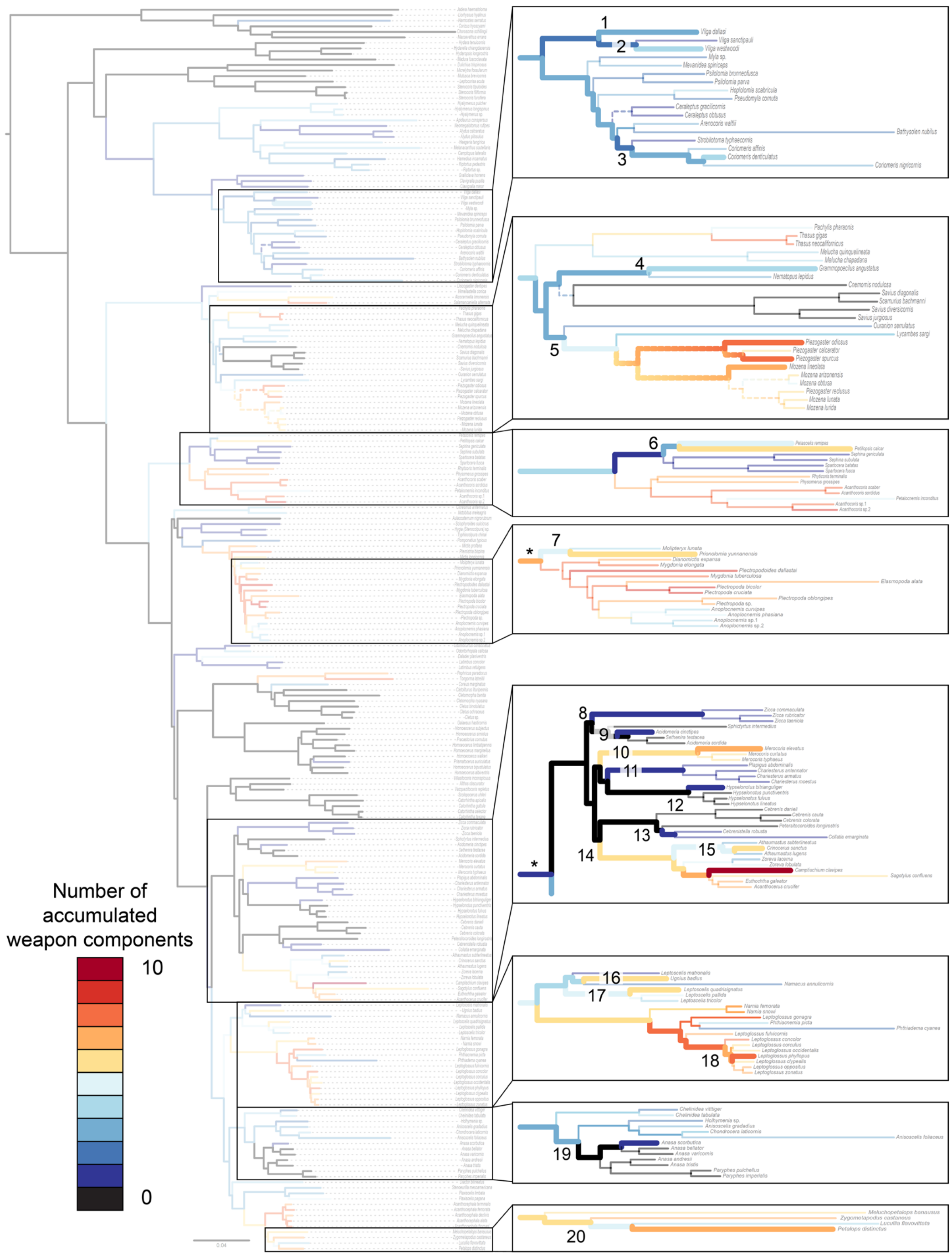


Fig. S54. Summary of the total number of component states accumulated on branches of the 50p ML phylogram, with insets at the right highlight areas of the tree where a cyclical pattern of increasing and decreasing elaboration were observed (each instance indicated by a number and thickened branches). Dashed lines indicate branches affected by at least one component having an ambiguous ancestral state; in this case, a color gradient is given to represent the range of the total number of components along the branch. Branches denoted with an asterisk in insets at the right have been modified (i.e. arbitrarily lengthened and not to scale) for visualization.

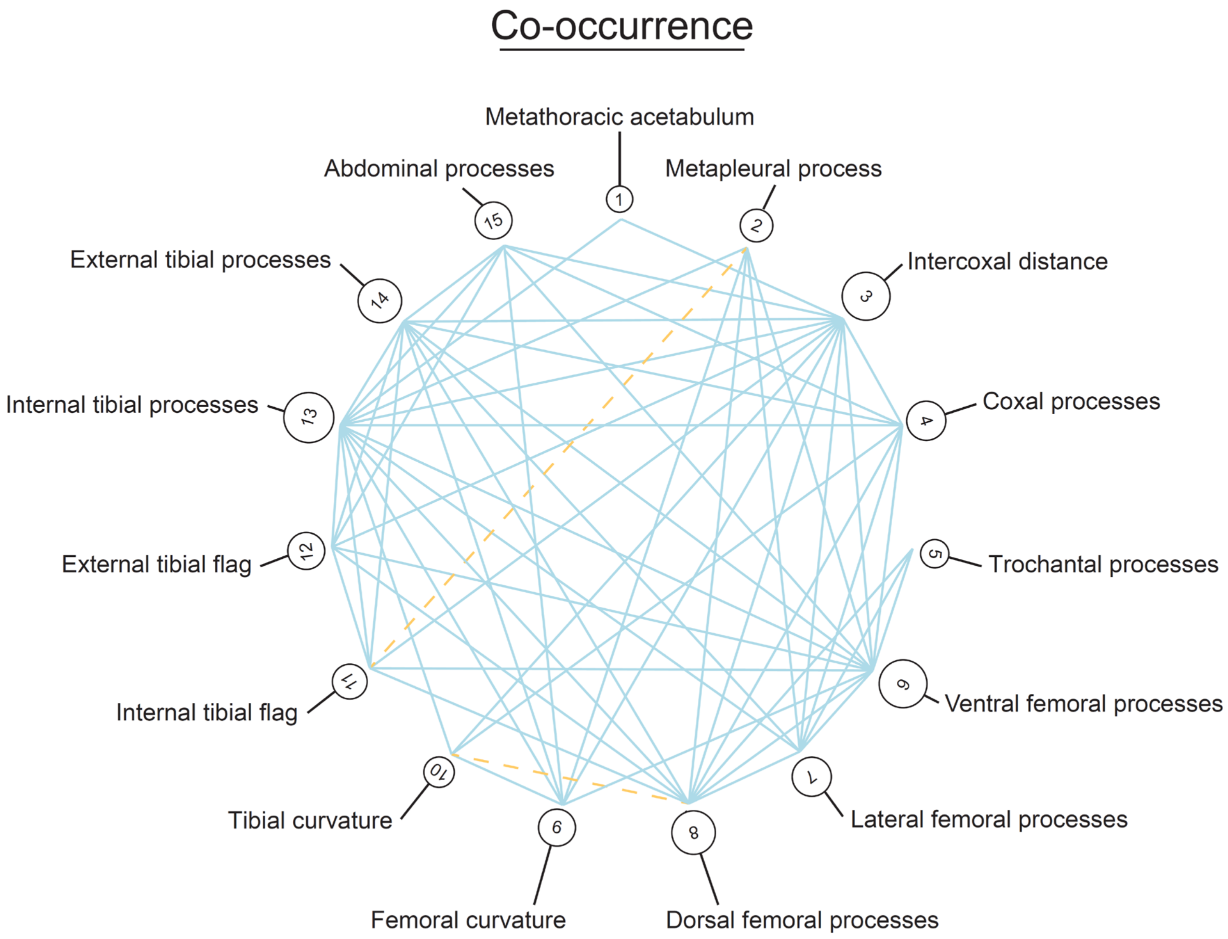


Fig. S55. Trait co-occurrences across 15 weapon components. Components with significant positive associations are shown in solid blue lines, while those with negative associations are shown in dashed yellow lines. The size of circles around character numbers reflects the total number of significant associations.

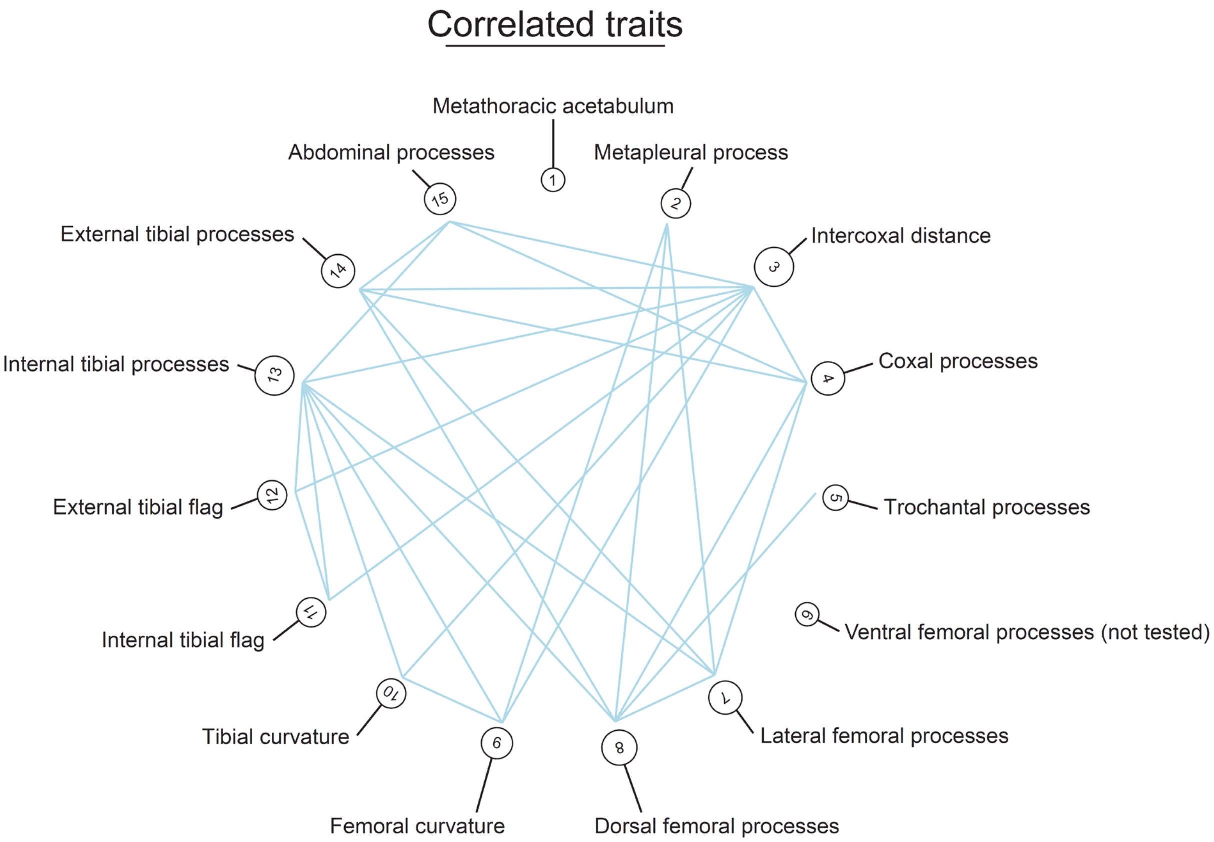


Figure S56. Correlated components based on the 50p MSC ultrametric tree and trait co-occurrence results. The size of circles around character numbers reflects the total number of significant associations.

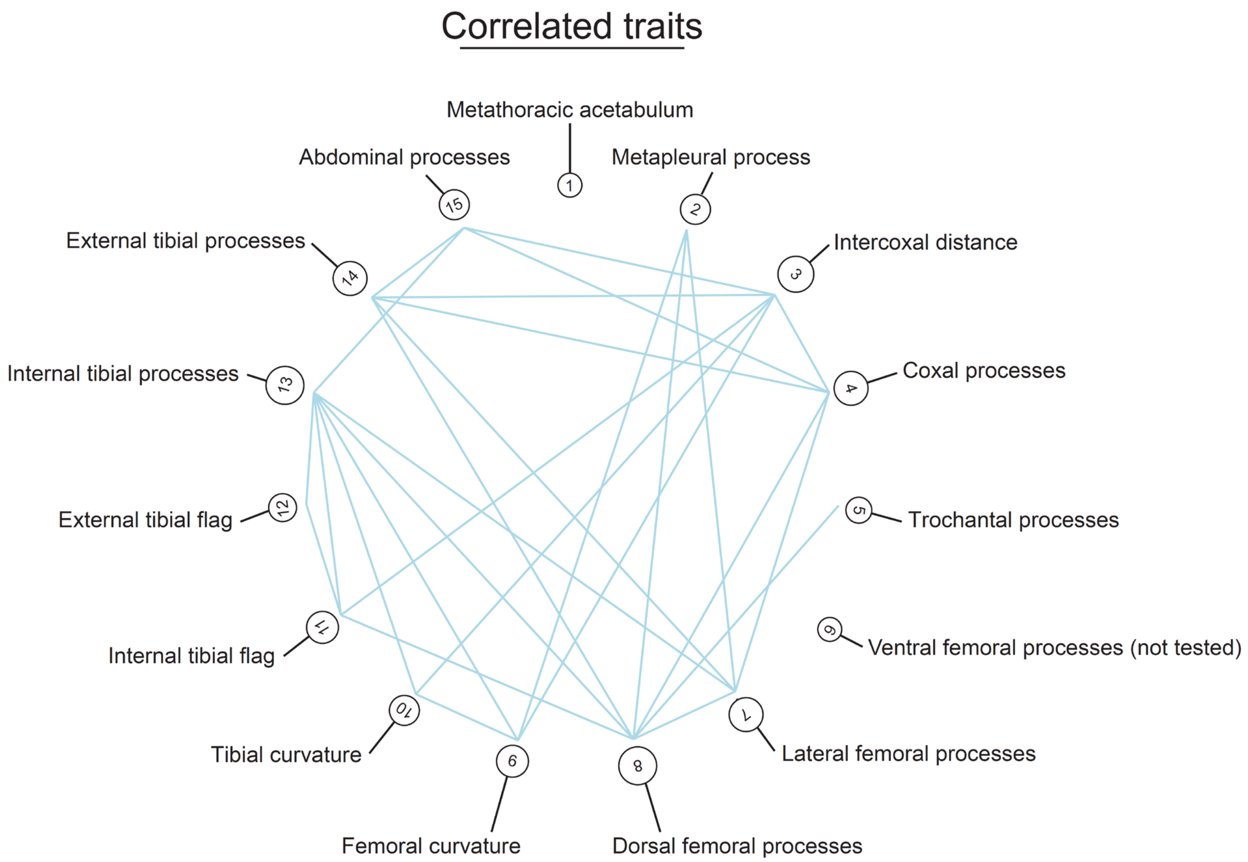


Figure S57. Correlated components based on the 50p25mi MSC ultrametric tree and trait co-occurrence results. The size of circles around character numbers reflects the total number of significant associations.
